# Unique genetic signatures of local adaptation over space and time for diapause, an ecologically relevant complex trait, in *Drosophila melanogaster*

**DOI:** 10.1101/2020.05.06.081281

**Authors:** Priscilla A. Erickson, Cory A. Weller, Daniel Y. Song, Alyssa S. Bangerter, Paul Schmidt, Alan O. Bergland

## Abstract

Organisms living in seasonally variable environments utilize cues such as light and temperature to induce plastic responses, enabling them to exploit favorable seasons and avoid unfavorable ones. Local adapation can result in variation in seasonal responses, but the genetic basis and evolutionary history of this variation remains elusive. Many insects, including *Drosophila melanogaster,* are able to undergo an arrest of reproductive development (diapause) in response to unfavorable conditions. In *D. melanogaster*, the ability to diapause is more common in high latitude populations, where flies endure harsher winters, and in the spring, reflecting differential survivorship of overwintering populations. Using a novel hybrid swarm-based genome wide association study, we examined the genetic basis and evolutionary history of ovarian diapause. We exposed outbred females to different temperatures and day lengths, characterized ovarian development for over 2800 flies, and reconstructed their complete, phased genomes. We found that diapause, scored at two different developmental cutoffs, has modest heritability, and we identified hundreds of SNPs associated with each of the two phenotypes. Alleles associated with one of the diapause phenotypes tend to be more common at higher latitudes, but these alleles do not show predictable seasonal variation. The collective signal of many small-effect, clinally varying SNPs can plausibly explain latitudinal variation in diapause seen in North America. Alleles associated with diapause are segregating at relatively high frequencies in Zambia, suggesting that variation in diapause relies on ancestral polymorphisms, and both pro- and anti-diapause alleles have experienced selection in North America. Finally, we utilized outdoor mesocosms to track diapause under natural conditions. We found that hybrid swarms reared outdoors evolved increased propensity for diapause in late fall, whereas indoor control populations experienced no such change. Our results indicate that diapause is a complex, quantitative trait with different evolutionary patterns across time and space.

**Author Summary:** Animals exhibit diverse strategies to cope with unfavorable conditions in temperate, seasonally varying environments. The model fly, *Drosophila melanogaster*, can enter a physiological state known as diapause under winter-like conditions. Diapause is characterized by an absence of egg maturation in females and is thought to conserve energy for survival during stressful times. The ability to diapause is more common in flies from higher latitudes and in offspring from flies that have recently overwintered. Therefore, diapause has been thought to be a recent adaptation to temperate climates. We identified hundreds of genetic variants that affect diapause and found that some vary predictably across latitudes in North America. We found little signal of repeated seasonality in diapause-associated genetic variants, but our populations evolved an increased ability to diapause in the winter when they were exposed to natural conditions. Combined, our results suggest that diapause-associated variants evolve differently across space and time. We find little evidence that diapause evolved recently in temperate environments; rather, SNPs associated with diapause tend to be quite common in Zambia, suggesting that diapause may promote survival under stresses other than cold. Our results provide future targets for research into the genetic underpinnings of this complex, ecologically relevant trait.

## Introduction

Organisms exhibit diverse strategies to survive environments that vary in space and time. Populations can undergo local adaptation, producing genotypic combinations that optimize fitness under present environmental conditions but may be less fit in other environments [1]. In temperate, seasonal locales, environmental signals are often used to anticipate the onset of the unfavorable season and induce plastic responses [2–4]. Insects exhibit a spectacular array of plastic responses to changes in season that can affect morphology [5–7], behavior [8], and developmental progression [9–11]. The ability to reprogram or arrest development, potentially for a fixed period of time (*diapause*), allows insects to weather unfavorable conditions in a hardy state of low metabolism [9–13]. Entry into diapause can result in tradeoffs between immediate survival and future growth, longevity, and reproductive success [14–16]. Due to these tradeoffs, some species exhibit local adaptation of their diapause response, producing dramatic differences in life history patterns across space and time [17–22]. Therefore, diapause is a plastic response to environmental changes, but the genetic ability to diapause can also differ across environments as a result of local adaptation. Intraspecific differences in diapause strategies offer an opportunity to study the spatiotemporal variation and evolutionary history of alleles contributing to a critical life history trait.

A number of reproductive life history traits vary in the genetic model organism *Drosophila melanogaster* [18, 23]. Female *D. melanogaster* are capable of arresting reproductive development and oogenesis when exposed to short day lengths (10 hours light: 14 hours dark, or 10L:14D) and low temperatures (10-14°C) soon after eclosion [24]. These winter-like conditions induce a change in hormonal signaling that prevents reproductive development [25, 26]. Physiologically, dormancy is regulated by hormones including insulin [27–29], juvenile hormone [25,30,31], and ecdysteroids [30, 32], as well as dopamine and serotonin [33]. *D. melanogaster*’s ovarian dormancy is accompanied by metabolic changes [34, 35] and is enhanced by starvation conditions [36]. Dormancy is coincident with transcriptional changes in up to half of all genes in *D. melanogaster* [37–39] and in other drosophilid species [40–42]. Male *D. melanogaster* also arrest spermatogenesis and undergo physiological changes under unfavorable conditions [43]. Dormancy subsequently diverts limited energy to survival rather than to reproduction at the onset of winter, perhaps allowing flies to overwinter *in situ* [44, 45] or in local refugia [46]. Whether this dormancy state is a true diapause or a state of quiescence is still open to debate [11], however it is clearly a substantial physiological reprogramming of the normal female egg production program that phenocopies diapause strategies of temperate endemic drosophilids [47–51]. Hereafter, we refer to *D. melanogaster’*s ovarian dormancy as diapause, keeping with established terminology in the field [for example, 18,24,26,46].

Clinal variation in the severity of winter is correlated with clinal variation in the ability to enter diapause in *D. melanogaster*. Flies from northern locations in North America, such as Maine, are more likely to be capable of entering diapause than those from more southern locations, such as Florida [18]. Additionally, the ability to enter diapause varies seasonally: the offspring of flies captured in the early spring (the survivors of winter, or their direct descendants) have a greater propensity for diapause than the offspring of flies captured in late summer [19], the descendants of lineages that prospered during favorable conditions [53]. Whether the same genetic loci underlie similar spatial and temporal evolution of phenotypes such as diapause remains unknown at a genome-wide level (but see [54]).

Adaptive evolutionary change in diapause propensity over ∼15 generations [55] during the growing season suggests the existence of tradeoffs related to diapause; while diapause is advantageous in unfavorable conditions, fitness costs occur when diapause is unnecessary [12]. In the absence of diapause-inducing conditions, strains capable of diapause have lower early reproductive success, putting them at a disadvantage when conditions are ideal for rapid reproduction. However, strains able to diapause tend to live longer, have greater reproductive success later in life, and are more tolerant of cold and starvation [15]. Laboratory selection experiments offer further evidence for tradeoffs: outbred flies reared under alternating cold and starvation stress in the lab evolve an increased genetic propensity to diapause, whereas flies reared under benign lab conditions evolve a decreased propensity for diapause [19], as do flies experimentally selected for heat tolerance [56]. Taken together, these finding suggest diapause is an ecologically relevant trait with tradeoffs that underlie local adaptation across both space and time.

In addition to being genetically polymorphic, *D. melanogaster*’s diapause is shallow relative to the diapause of other insects, including other drosophilids [57, 58]. Females may spontaneously resume ovarian development after ∼6 weeks even when diapause-inducing conditions persist [24] (but see [59]), and diapause is rapidly broken if temperature or day length is increased [24]. In a short-lived species with many generations per year, this weak and facultative diapause strategy may be advantageous to allow individuals to quickly reenter reproductive mode soon after conditions become favorable. On the other hand, the relatively recent colonization of temperate habitats [53–57] led to the suggestion that seasonal diapause may be a recently evolved trait [24, 60], so the weak diapause of *D. melanogaster* might reflect its incipient evolution. Phenotypic plasticity, such as diapause or behavioral modification, is predicted to evolve when the environment varies more rapidly than generation time, whereas fixed alternative strategies are favored when environmental variation occurs more slowly than generation time [61]. However, in temperate *D. melanogaster*, the scale of temporal environmental variation is similar to generation time (weeks), which may explain the reversibility of diapause and the substantial genetic variation in diapause induction over seasonal timescales.

Despite our extensive knowledge of the natural history and physiological basis of diapause in *D. melanogaster* and other insects, we still know little about the identity and evolutionary history of polymorphisms underlying variation in this critical life history trait [62]. Herein, we sought to identify genetic variants underlying diapause via a genome wide association study (GWAS) and to link these variants to patterns of global polymorphism in *D. melanogaster*. After identifying hundreds of single nucleotide polymorphisms (SNPs) with small effects on diapause, we specifically addressed two questions about the evolutionary history of these alleles. First, do SNPs underlying diapause show predictable patterns of genetic variation across latitudes and seasons, with pro-diapause alleles more common in northern latitudes and in the spring? Second, are alleles associated with diapause present in ancestral populations, and have they experienced recent selective sweeps in North America? Our results suggest that the evolution of diapause across spatial gradients may be distinct from its evolution across seasons: while diapause-associated alleles are weakly clinal, they do not tend to vary predictably over seasons across multiple populations. Furthermore, alleles controlling diapause represent ancestral genetic variation, suggesting they may play roles in seasonal, or even general, aspects of stress response in ancestral localities. The favorability of diapause-associated alleles under a variety of stressful conditions could explain the lack of repeated seasonality of these alleles as well as their presence in tropical climates with pronounced seasonality. Our results provide a roadmap to understand the role of small-effect alleles underlying the evolution of a complex, ecologically relevant trait that varies across space and time.

## Results

In order to characterize the genetic polymorphisms that underlie variation in diapause in *D. melanogaster*, we used a novel hybrid-swarm based mapping approach [63] employing sequenced, genetically diverse inbred lines collected around North America and the Caribbean (Fig 1A; S1 Fig; S1 Table). We intercrossed these lines to produce two outbred populations with recombinant genotypes (populations A and B, Fig 1B; S2 Fig) and then exposed these hybrid individuals to various diapause-inducing conditions in custom-built chambers (Fig 1C). We mapped the genetic basis of diapause by dissecting and genotyping 2,823 females (Fig 1D). We used the results of this GWAS to analyze patterns of variation in SNPs associated with diapause.

**Fig 1.**
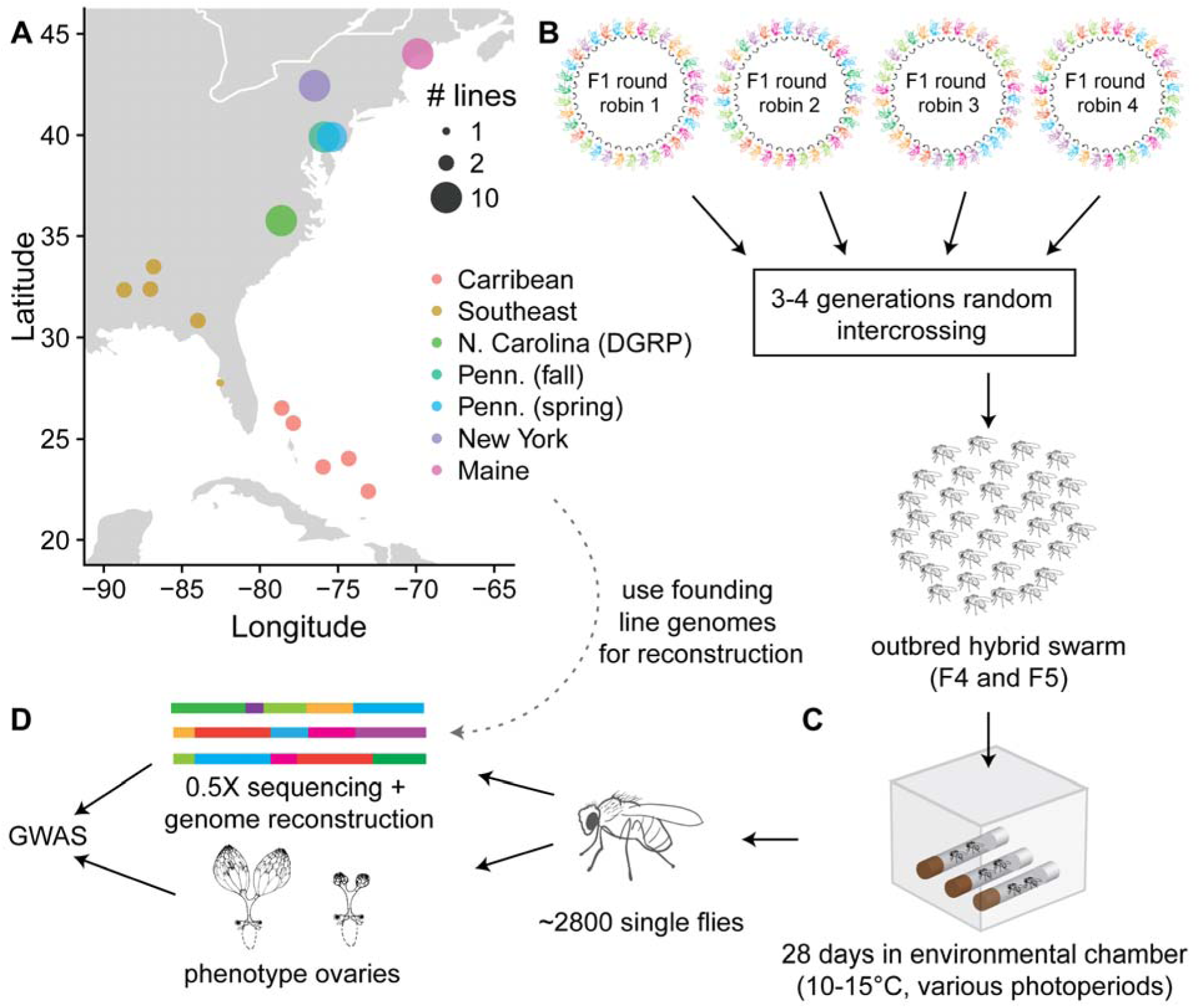
Experimental design for hybrid-swarm based association mapping of diapause. (A) A total of 68 sequenced, inbred lines from seven North American collections were used to initiate hybrid swarm crosses. The lines were divided into two groups of 34 (populations A and B) so that each group had equal representation from each collection. Map generated using [64]. (B) Within each population, randomly ordered round-robin crosses were established. The F1 adults were released into cages and propagated with non-overlapping generations. C) Virgin females from the F4 and F5 generations were collected and placed in environmental chambers with varying photoperiods and temperatures for 28 days. D) Individual flies were dissected to phenotype diapause. DNA was extracted from the carcasses and individually sequenced to approximately 0.5X coverage. Sequencing reads were used to reconstruct full genome sequences and perform a genome-wide association study. Ovary drawing in D) modified from [65] under a CC-BY license.

### Temperature and nutrition affect diapause

We initially assessed ovarian development in F4 and F5 hybrid swarm individuals exposed to a range of cool temperatures (10-16 °C, S3 Fig) and photoperiods representing the approximate range of day lengths experienced by our northernmost Maine population. We recorded the stage of the most advanced non-stage 14 ovariole [66] (Fig 2A) and counted the number of mature eggs in each individual. We found a strong effect of temperature on ovarian development: higher temperatures led to more advanced ovarian development and more eggs (Fig 2A-B). When examining the most advanced pre-stage 14 ovariole in flies that had produced eggs, we also found that higher temperatures increased the proportion of individuals with advanced ovariole stages (Fig 2C).

**Fig 2.**
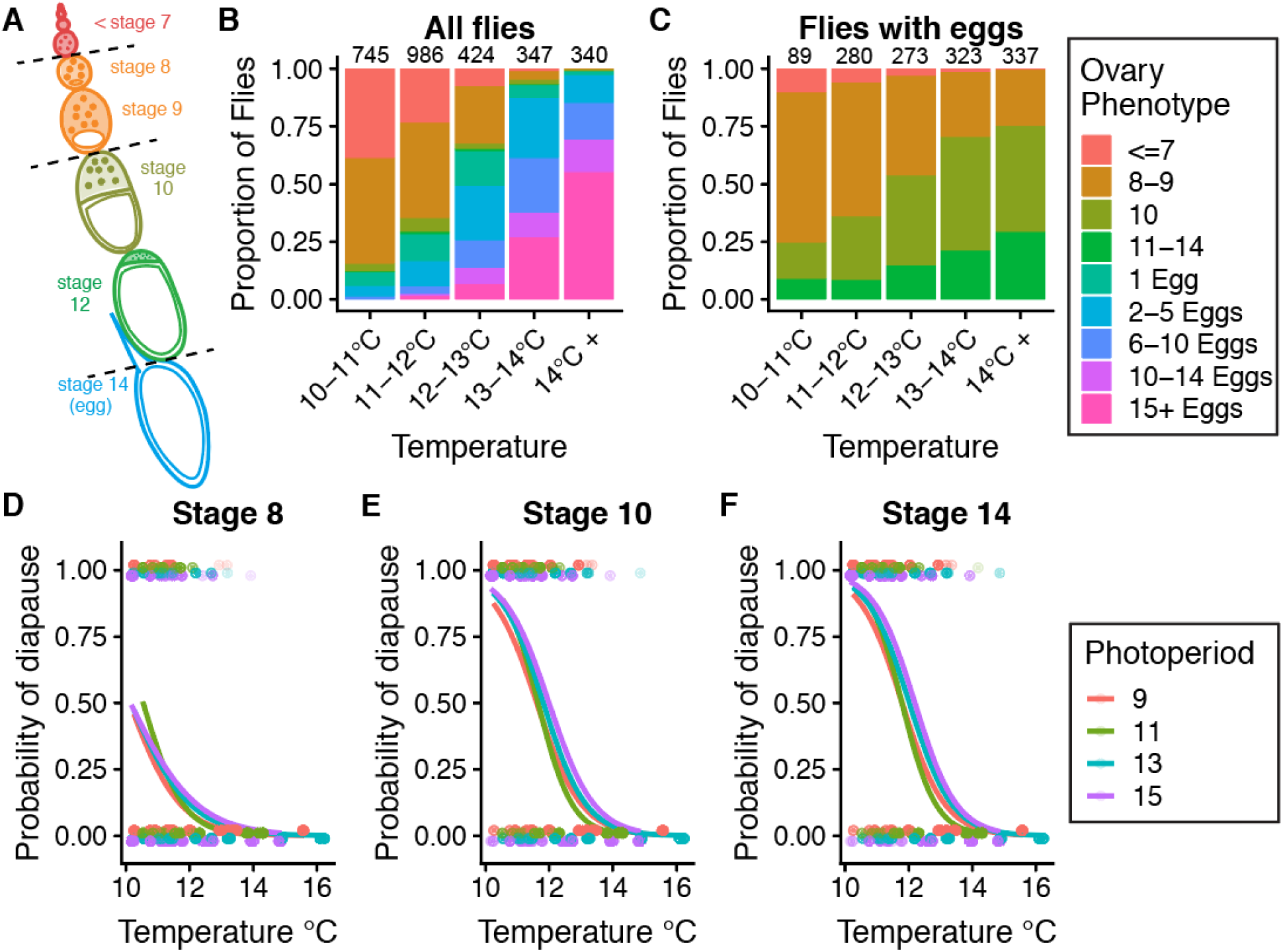
Effects of temperature and photoperiod on diapause in hybrid swarm populations. (A) Ovaries were scored based on the most advanced ovariole stage and the total number of eggs, according to King (1970) [66]. Dashed lines indicate three cutoffs for scoring diapause: stage 8, stage 10, and stage 14. Colors correspond to panels B and C. (B) The most advanced ovariole stage, proportion of flies with eggs, and the total number of eggs per individual all increase with increasing temperature. Numbers above bars indicate the total number of flies phenotyped in each temperature range. (C) Among flies with eggs, the stage of the most advanced egg chamber also increases with temperature. (D-F) Diapause incidence, scored at the stage 8, stage 10, or stage 14 cutoffs, decreases with increasing temperature (binomial general linear model, *P* < 2 x 10^-16^ for all phenotypes). Unexpectedly, longer photoperiods (shown as hours of light: hours of dark) result in increased diapause incidence (binomial *glm*, *P* = 0.002, *P* = 2.7 x 10^-7^, *P* = 5.0 x 10^-8^, respectively). Gray points represent individual fly phenotypes (diapause = 1, non-diapause = 0).

We initially scored diapause as an absence of development past three different ovariole stages [66]: stage 8 (no ovarioles past stage 7), stage 10 (no ovarioles past stage 9), and stage 14 (no mature eggs present) (Fig 2A). Diapause at stage 8 and stage 10 were modestly correlated (Pearson correlation, *R* = 0.49, *P* = 2 x 10^-172^) and diapause at stage 10 and stage 14 were strongly correlated (*R* = 0.92, *P* < 1 x 10^-200^). After accounting for differences caused by temperature, photoperiod had an effect on diapause in the opposite direction predicted: individuals exposed to long day (13L:11D or 15L:9D) light cycles tended to have higher diapause induction (Fig 2D-F), regardless of the ovariole stage cutoff used to determine diapause. Based on recent studies [35], previous work [18,19,52,59,67], and developmental evidence that a checkpoint exists between stages 9 and 10 in ovariole development [68–70], we chose to classify diapause either as an absence of ovariole development to stage 8 (Fig 2D) or stage 10 (Fig 2E) for the remaining genetic analysis.

We next investigated other factors besides temperature and photoperiod that may affect diapause. The F4 generation of our outbred populations had increased diapause incidence at all temperatures (S4A Fig; *P* < 2 x 10^-16^) relative to generation F5, potentially reflecting inadvertent rearing differences (such as larval density) between generations. Outbred populations A and B also showed significantly different diapause induction across temperatures (S4B Fig, *P* < 0.01), though the differences were much less pronounced than the effect of generation. In an experiment conducted in the F20 generation of the hybrid swarm, we found that feeding adults supplemental live yeast substantially reduced diapause incidence across a range of temperatures, further suggesting that adult nutrition and temperature contribute to diapause induction (S5 Fig, *P* < 1 x 10^-10^).

Lastly, we tested for an influence of *Wolbachia* infection on diapause. We quantified the proportion of sequencing reads mapping to the *Wolbachia* genome for each hybrid individual to infer *Wolbachia* infection status. We found that *Wolbachia* decreased the likelihood of diapause at stage 8 (diapause was observed in 17% of *Wolbachia*-infected individuals vs 23% of uninfected individuals; Fisher’s exact test; *P* =0.0001) but did not influence diapause at stage 10 (*P* = 0.31). Placing all of these variables into a single model revealed that temperature and generation accounted for the majority of explainable variation in the diapause phenotype, though more than 50% of the variation remains unexplained (S2 Table).

### Diapause is heritable with a polygenic signal

We estimated a narrow-sense heritability of diapause using genome-wide genotypes in a restricted maximum likelihood analysis in GCTA [71, 72]. We estimated heritabilities of approximately 0.12 and 0.08 for stage 8 and 10 diapause, respectively, using all genotype data from both populations combined. These estimates far exceeded the heritability calculated for 1,000 permuted phenotype datasets (Fig 3A; B). To characterize the genetic architecture of natural variation in diapause, we performed a GWAS using reconstructed genotypes (S6 Fig), from 2,823 hybrid individuals for diapause scored at two cutoffs. We performed this analysis, using 100 different imputations of missing genotype data, in populations A and B separately, as well as the two populations combined (“both”). We also generated 1000 permutations of the combined dataset and 100 permutations of each individual population. We then calculated the genomic inflation factor, λ_GC_, for the mapping results of each phenotype in each permutation. For all three mapping populations, λ_GC_ was slightly greater than 1, whereas the permutations ranged from ∼0.8 to ∼1.1 (Fig 3C). The slight inflation of λ_GC_ is indicative of the polygenic basis of diapause and was variable across chromosomes (S7 Fig). Visual inspection of a representative Manhattan plot also revealed a broadly polygenic signal of diapause, with SNPs scattered throughout the genome (Fig 3D). We note relatively few alleles associated with diapause on the X chromosome, which appears to be specifically caused by a lack of associations in population B (S7 Fig). Surprisingly, X is the only chromosome that appears to be more diverse in population B relative to A (S2 Fig).

**Fig 3.**
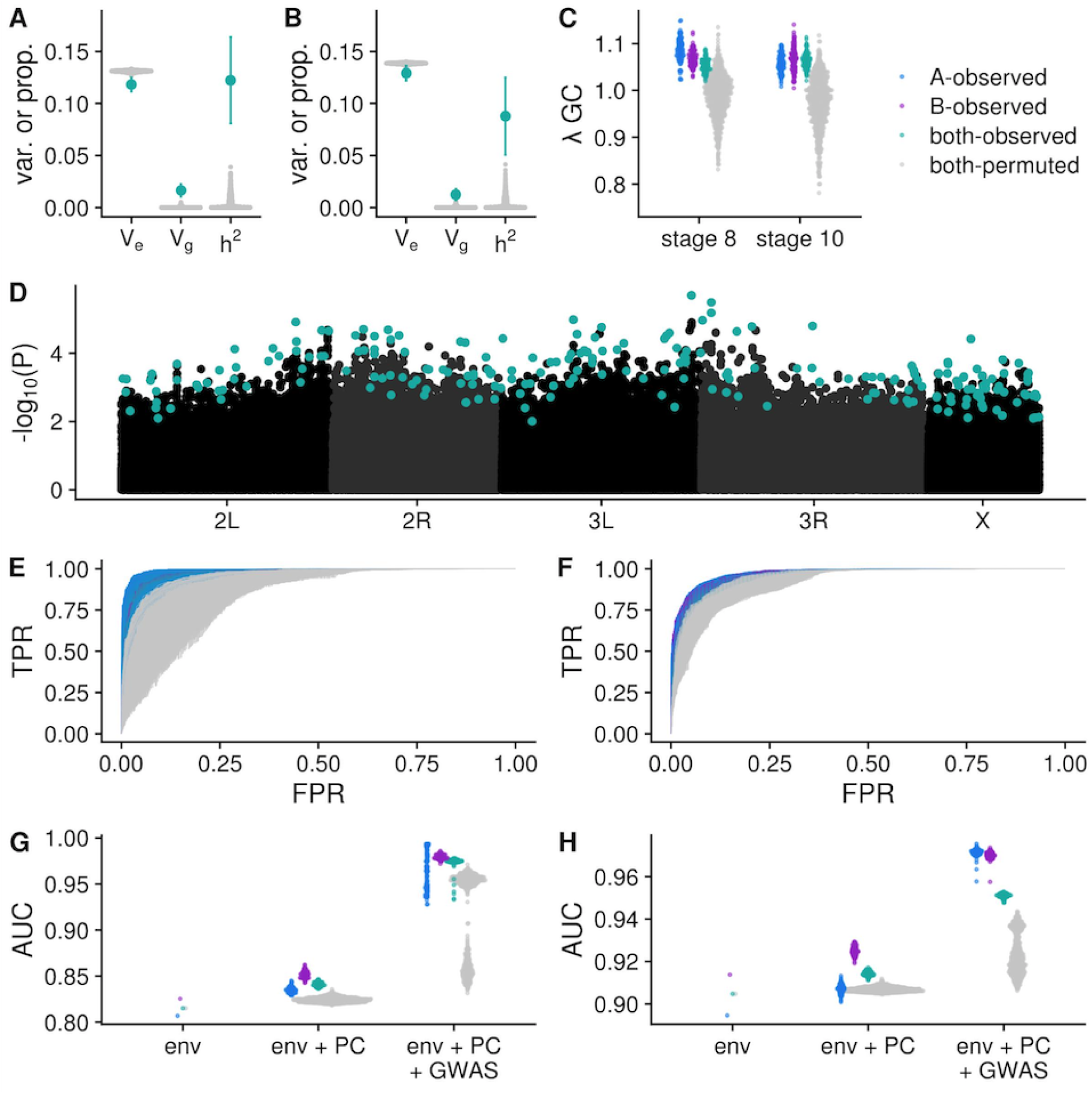
Diapause is a polygenic trait. (A-B) V_g_, V_e_ and heritabiilty estimates for stage 8 (A) and stage 10 (B) diapause phenotypes. Teal points indicate observed estimates +/- 95% confidence interval for heritability in the hybrid swarm (populations A and B combined). Grey points indicate heritability estimates for 1000 permutations. (C) Genomic inflation factor (also known as _GC_) for GWAS in 100 imputations of actual data (teal=A+B, blue=A, purple=B) and 1^λ^000 permutations of A+B (grey). (D) Manhattan plot for GENESIS *P*-values for stage 10 diapause in one imputation. Teal points indicate LASSO SNPs. (E-F) Receiver operating characteristic (ROC) curves for stage 8 (E) and stage 10 (F) diapause predictions made using LASSO SNPs. Phenotypes for each individual in the mapping population were predicted using the informative environmental variables, genetic principal components, and SNPs chosen by LASSO. At any given false positive rate (FPR), the observed data (blue, purple, and teal lines) have higher true positive rates (TPR) relative to permutated GWAS (gray lines). (G-H) Quantification of ROC analysis using area under the curve metric (AUC). “env” is a model containing only environmental data; “env + PC” includes environmental data and 32 principal components; “env + PC + GWAS” includes the former plus the genotypes of up to several hundred SNPs chosen by LASSO.

Following the GWAS, we used LASSO [73] to identify a subset of unlinked, informative SNPs (hereafter, “LASSO SNPs”) from the top 10,000 SNPs ranked by *P*-value (Fig 3D). LASSO takes a number of potential predictor variables and chooses those that are most informative yet independent [74]. Each LASSO model started with environmental covariates, the top 32 principal components derived from genome-wide SNPs, and the SNP genotypes; the resulting model usually retained several hundred SNPs. LASSO SNPs (S6 Table) showed low levels of LD (S8 Fig), suggesting the algorithm was successful in choosing unlinked informative markers. More LASSO SNPs were identified in the observed data relative to permutations (S9 Fig), demonstrating increased genetic signal in the true ordering of the data. However, we note that LASSO SNPs may be markers of important haplotypes and not causative loci.

To determine whether the SNPs identified by LASSO were in fact informative for predicting diapause phenotypes, we implemented a receiver operating characteristic curve (ROC) analysis [75]. The SNPs chosen by LASSO substantially improved the accuracy of the predicted phenotypes, with a higher true positive rate (TPR) and lower false positive rate (FPR) in the observed data relative to the permutations (Fig 3E; F, compare blue/green/purple lines to grey). This difference can be quantified using the area under the curve (AUC), which was higher for predictions in the observed data relative to the permutations for both phenotypes (Fig 3G; H). Additionally, we found that for both phenotypes, adding genetic principal components to the model improved the model relative to environment alone, and adding LASSO SNPs further improved the model over principal components plus environment (Fig 3G; H). This improvement was greater in the actual data relative to the permutations; combined, these observations suggest true genetic signal in the observed data.

We tested for shared genetic signal for diapause between the two mapping populations by counting the number of SNPs shared between populations A and B for each imputation and permutation of the data. We found that the number of shared LASSO SNPs or quantile-ranked SNPs was very small, and the overlap between SNPs identified from actual data did not exceed the overlap between SNPs identified in permutations (S10A Fig). However, we did observe an excess of shared SNPs associated with both of the two phenotypes. We found that for the quantile-ranked cutoffs, the number of SNPs shared in the observed data was generally greater than the number of SNPs shared between permutations (S10B Fig). This observation suggests that some loci affect diapause at both stages. In contrast, we observed little overlap of LASSO SNPs between the two phenotypes, again indicating that LASSO SNPs may be markers and not causative.

Intriguingly, we found no effect of two previously described variants (*cpo* SNP: *dm3* chr3R: 13,793,588; and *timeless* indel: chr2L:, 3,504,474) [52,54,60,76] on diapause (S11 Fig). We tested for association between diapause and karyotype at five cosmopolitan inversions [In(2L)Ns, In(2R)t, In(3R)Payne, In(3R)C, and In(3R)Mo] segregating in the hybrid swarm. We found that individuals heterozygous for In(3R)Payne had a modest decrease in diapause induction at both stages in population A and both populations combined (S3 Table); however, these effects were not significant after correcting for multiple testing of the various inversions. Therefore, the cosmopolitan inversions do not appear to play major roles in diapause induction. We also investigated the functional annotation categories of SNPs associated with diapause. The vast majority of diapause-associated SNPs were found in non-coding regions: upstream/downstream of annotated genes, or in introns (S4 Table). However, there was little signal of enrichment or de-enrichment for particular classes of diapause-associated SNPs relative to permutations (S5 Table).

### Linkage is not responsible for the polygenic basis of diapause

Collectively, these results suggest that many loci throughout the genome contribute to phenotypic variation in diapause. We speculated that the nature of our mapping population might increase linkage disequilibrium (LD), potentially causing large linkage blocks to be associated with diapause. To test this possibility, we examined the general signal of LD in our mapping population in contrast with a wild-derived population from North Carolina (the Drosophila Genetic Reference Panel, or DGRP [77]). We also analyzed the 68 parental lines that produced the hybrid swarm, and randomly down-sampled the DGRP to 68 lines for comparison. Four to five generations of mating reduced LD in the hybrids relative to the parental lines (Fig 4A-B; compare pink and teal). The LD of the DGRP is slightly lower than that of the hybrid swarm (compare orange and teal); however, this effect is partially mediated by number of lines, as the down-sampled DGRP (yellow) had higher LD than the hybrid swarm at larger distances. None of the mapping populations demonstrated substantial long-distance LD between arms of the same chromosome or across chromosomes (Fig 4C-E). Therefore, while LD between nearby SNPs may contribute to some signal in the GWAS, the polygenic signal is not solely due to linkage, since LD drops below 0.05 within ∼10 kb in the F4s and F5s of the hybrid swarm.

**Fig 4.**
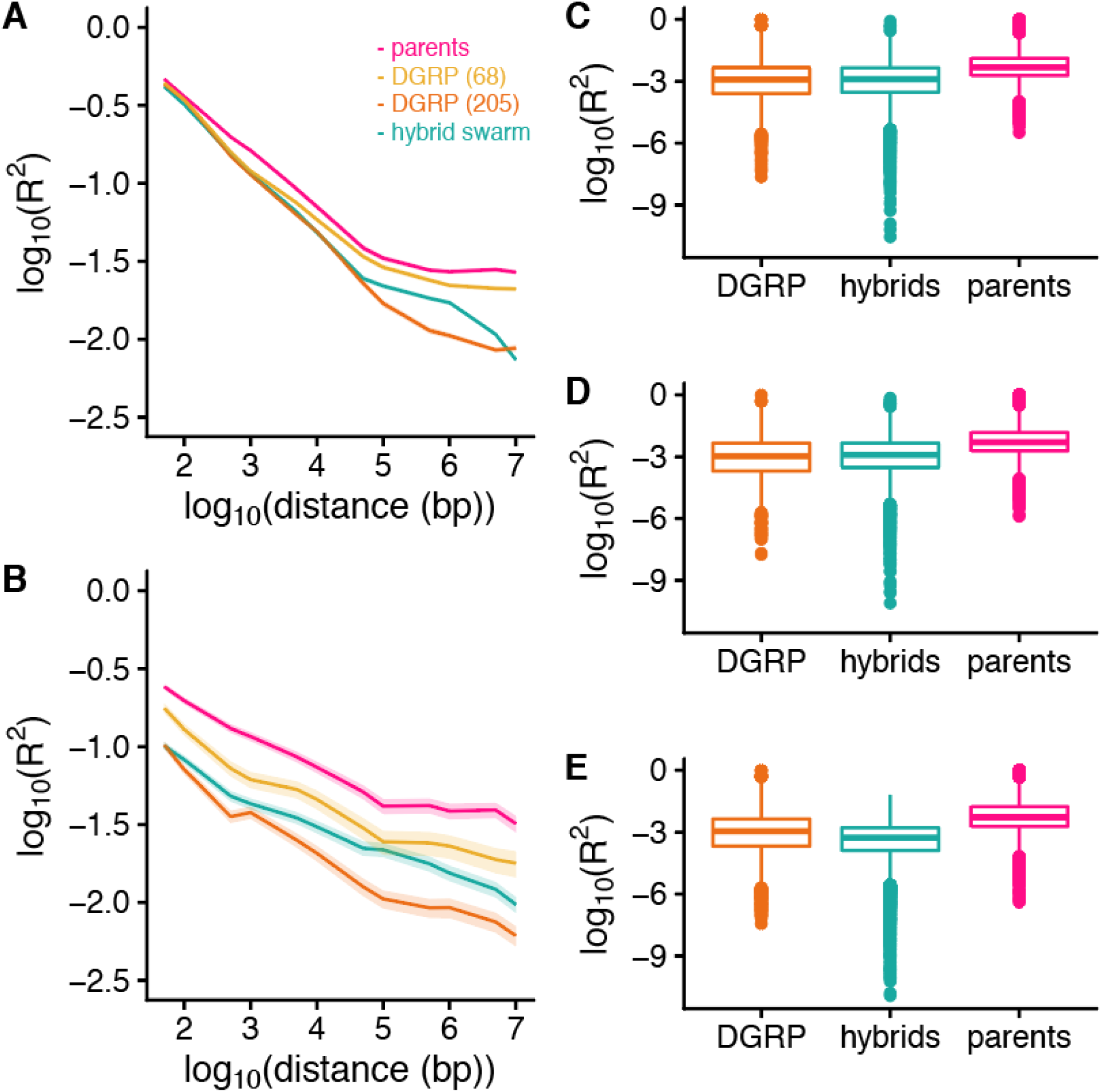
Analysis of linkage disequilibrium supports a polygenic basis of diapause. (A-B): Linkage disequilibrium (LD) decay in the hybrid swarm (teal) contrasted to the DGRP (orange), DGRP down-sampled to 68 lines (yellow), and hybrid swarm parents (pink) in common (minor allele frequency > 0.1, A) and rare (MAF < 0.05, B) SNPs. 10,000 SNPs were randomly sampled from common SNPs with minor allele frequency (MAF) > 0.1 or rare SNPs with MAF < 0.05. LD (R^2^) to nearby SNPs at fixed distances was measured. Lines represent median LD; ribbons represent the 95% confidence intervals. (C-E) Long distance LD between pairs of SNPs randomly sampled from 2L and 2R (C), 3L and 3R (D), or sampled from different chromosomes (E). All analysis was conducted on populations A and B combined.

### Diapause-associated SNPs are likely to be clinal, but not seasonal

We tested for signatures of local adaptation across space (a north to south cline in North America) and time (between fall and spring) by intersecting the GWAS results with existing datasets of spatiotemporal SNP variation in *D. melanogaster* [53, 78]. Based on *D. melanogaster’s* natural history [18, 19], we predicted that pro-diapause alleles (which increase diapause in the GWAS) would be at higher frequency in the north and in the spring. We used LASSO SNPs as well as genome-wide quantile thresholds to conduct this analysis. We calculated a polygenic score [79, 80] by multiplying the clinal or seasonal effect size (including sign) by the GWAS effect size and sign and summing this product across all SNPs, all LASSO SNPs, the top 0.01% of the GWAS, or the top 0.1% of the GWAS (see S6 Table and Materials and Methods for details). In this test, we predict that concordant signal in the GWAS and the clinal/seasonal datasets should result in positive numbers that exceed permutations.

We first compared our data to the results from Bergland *et al* (2014), which sampled allele frequencies across a latitudinal cline and also identified SNPs that repeatedly oscillate in allele frequency between spring and fall in a single Pennsylvania orchard over three years. We found that in population A and both populations combined, the average clinal polygenic score of the actual data was positive and exceeded the permutations for the stage 8 diapause phenotype for the top 0.01% and 0.1% of GWAS SNPs (Fig. 5A). Therefore, the combination of many small-effect alleles that vary clinally could produce the clinal variation of diapause observed in North America. This trend was not observed in LASSO SNPs, suggesting that LASSO SNPs may not be the subset of SNPs with the greatest ecological relevance, or that the clinal signal that we observe is partially driven by some linked sites among quantile-ranked SNPs. No strong trends were observed for the SNPs associated with stage 10 diapause, suggesting the possibility of different spatial selection pressures on SNPs underlying the two phenotypes. Furthermore, no strong seasonal trends were observed for either phenotype (Fig 5B), suggesting that pro-diapause alleles in our mapping population do not repeatedly increase in frequency in spring relative to fall in natural populations. We performed the same analysis on data collected by Machado *et al* (2019), which also sampled a cline and compared spring and fall allele frequencies in 20 populations sampled from North America and Europe. We observed a similar trend of clinal signal in diapause-associated alleles for the stage 8, but not stage 10, data, but limited parallelism of diapause effects and seasonal signal in this dataset (S12 Fig).

**Fig 5.**
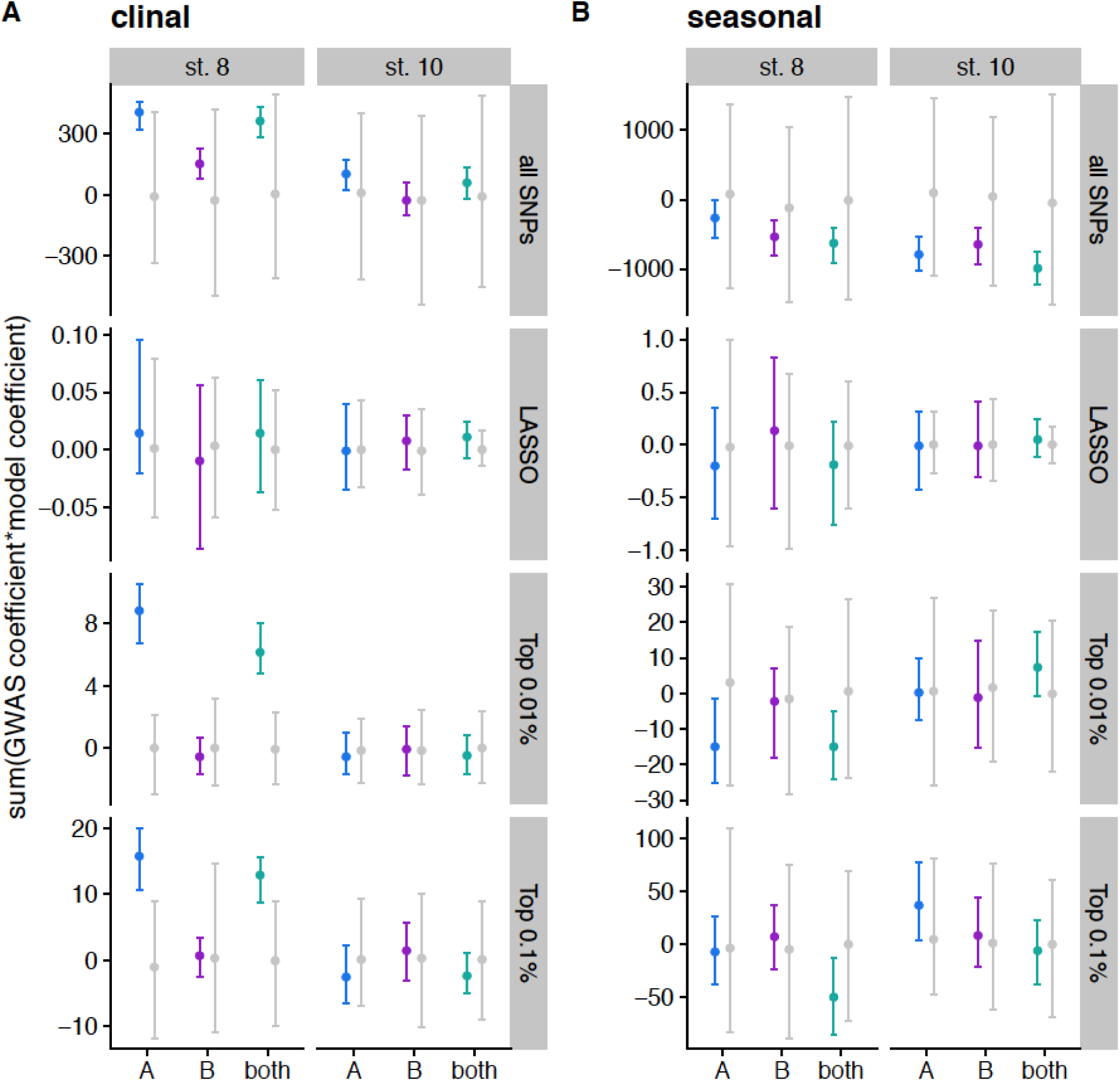
SNPs associated with diapause at stage 8 vary predictably across latitudinal clines. A) Polygenic scores calculated by multiplying clinal effect size reported in Bergland *et al* (2014) and GWAS effect size for each SNP and summing across all SNPs, LASSO SNPs, the top 0.01% of the GWAS, and the top 0.1% of the GWAS. Effect sizes are polarized such that positive numbers indicated pro-diapause alleles are more common in the north. Data are shown with a point for the mean and error bars extending to the 2.5% and 97.5% quantiles. Colored points indicate actual data for 100 imputations of each mapping population; grey points indicate the distribution for permutations. B) Polygenic scores calculated for seasonal data by multiplying seasonal effect size reported in Bergland *et al* (2014) and GWAS effect size, polarized so that pro-diapause and spring are positive.

The lack of predictable polygenic signal in the seasonal tests led to the intriguing possibility that diapause might evolve seasonally via changes in allele frequencies of different SNPs in different populations. We tested for concordant signal of diapause-associated SNPs among seasonally varying polymorphisms from 20 individual populations sampled in Machado *et al* 2019 [78]. Some, but not all, populations showed predictable seasonal allele frequency changes of diapause-associated SNPs (S13 Fig; S14 Fig), though the direction varied, with some populations showing an excess of pro-diapause alleles in the spring (positive scores), and others showing an excess of pro-diapause alleles in the fall (negative scores). Therefore, we suggest that seasonal evolution of diapause-associated SNPs can and does occur in individual populations, but in an unpredictable manner across years and populations.

### Hybrid swarms evolve an increased capacity for diapause in the winter in natural environments

To study the natural seasonal dynamics of diapause expression, we introduced our hybrid swarm populations to outdoor cages fed regularly with rotting apples and bananas to experience semi-natural “wild” conditions (S15 Fig). These cages were not density controlled and were composed of individuals of various ages because they were allowed to propagate freely, with overlapping generations. We sampled flies periodically from these cages over six months and assessed ovary status in the captured females. Surprisingly, we found that a substantial proportion of flies appeared to be in diapause, even during the favorable months of summer and early fall. We refer to these flies as “diapause-like” since they were not placed in the standard laboratory diapause assay as virgin females. All field-caught samples (G0) from fruit-fed cages (Fig 6A, green lines) had higher incidence of diapause-like ovaries than a reference sample of flies reared in lab cages at 25°C (Fig 6A, grey dashed lines). The single highest rate of diapause-like ovaries for stage 10 was observed from a sample in mid-December that was collected in sub-freezing conditions (Fig 6C), suggesting that cold temperatures may be one of several factors that reduce ovary maturation in natural conditions. However, the highest diapause incidence for stage 8 was observed in October, suggesting other factors besides cold temperature may contribute as well, and the two phenotypes may respond to somewhat different signals. Intriguingly, diapause incidence of field caught samples was quite low in November, even though conditions were cool relative to previous months. This observation further suggests that environmental factors beyond temperature mediate ovary development.

**Fig 6.**
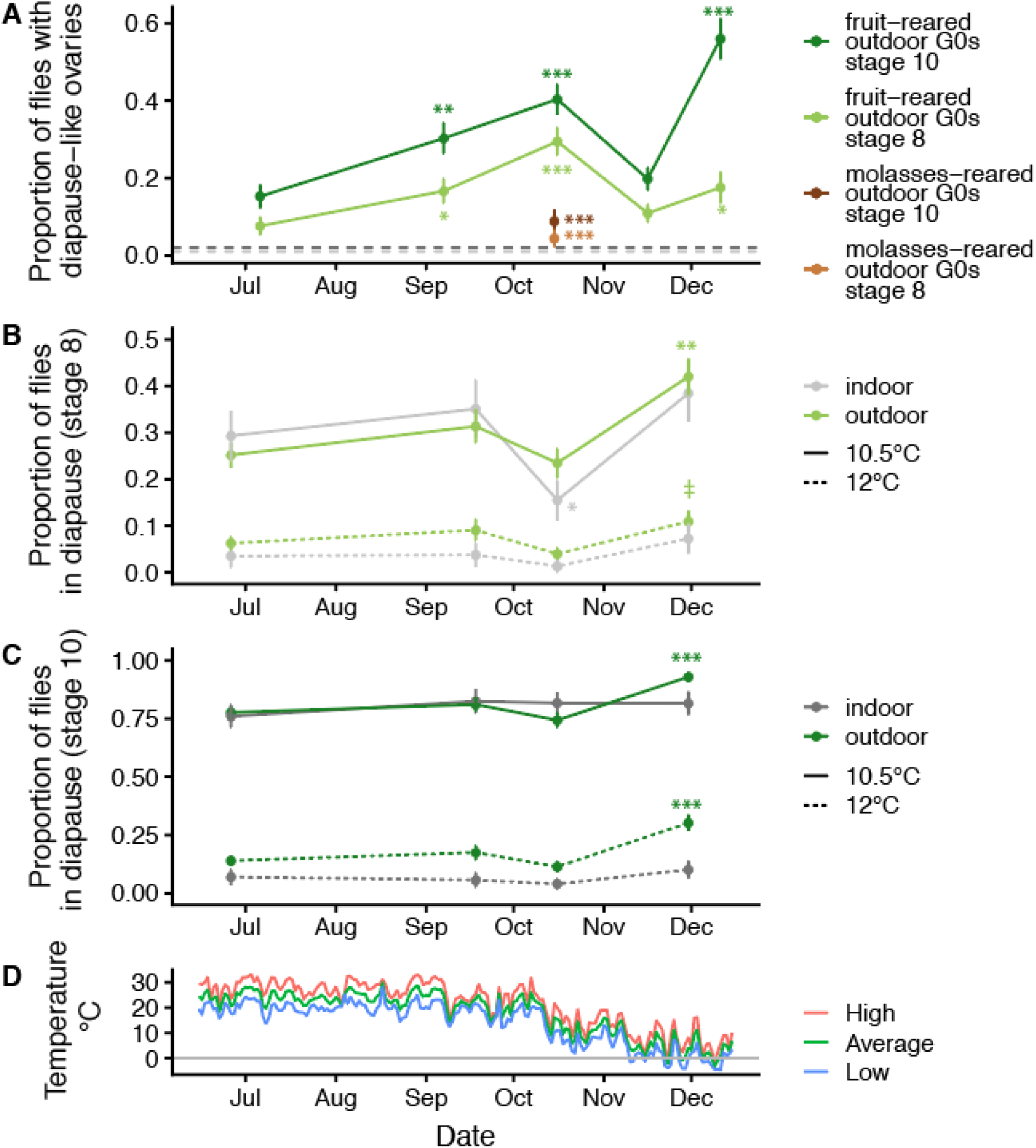
Plasticity and selection contribute to increased diapause in late fall in field-reared samples. (A) Ovaries were dissected from outdoor cage flies reared on fruit and diapause was assessed at stage 8 (light green) or 10 (dark green). A general linear model was used to determine whether diapause at later collection dates differed significantly the first collection on June 26^th^, 2018. (‡ P < 0.1, * *P* < 0.05, ** *P* < 0.01, *** *P* < 1x 10^-4^). Samples were also collected at a single time point from cages reared on cornmeal-molasses food (brown). This sample had significantly lower diapause incidence than fruit-reared samples collected one day later at both stages. The horizontal grey lines indicate diapause incidence for stage 8 (light grey) and stage 10 (dark grey) in a single sample of 96 mated, 5-8 day old flies reared in laboratory cages on cornmeal-molasses food. (B) Field cage and laboratory cage flies were collected at several timepoints and reared in the lab for two generations before assessing diapause at stage 8 in the standard assay at either 10.5 °C (solid lines) or 12 °C (dashed lines). Diapause was marginally increased in the December outdoor sample at 12 °C (*P* = 0.08) and significantly increased in December at 10.5 °C (*P* = 0.0002), whereas it remained relatively consistent across indoor flies. (C) Same as B, but phenotypes for stage 10 diapause. Diapause increased in the December sample of outdoor cage flies relative to the first sample from June (general linear model, *P* = 3.7 x 10^-5^ at 12 °C, *P* = 6.1 x 10^-5^ at 10.5 °C), while diapause was consistent across samples in the lab-reared flies (*P* > 0.05 for all pairwise comparisons). (D) Weather Underground temperature data during the field season for Carter Mountain, VA, approximately 2 km from our field site.

Diapause in the field was influenced by nutrition: flies sampled from a separate set of outdoor cages that were fed cornmeal-molasses food revealed a much lower proportion of flies in diapause when compared to fruit-reared flies captured just one day later (Fig 6A; compare brown and green points in October). Therefore, fruit consumption, or the microbiota associated with fruit in the wild [81], may cause reduced ovary maturation relative to standard lab medium. This finding agrees with our result that live yeast diminishes diapause (S3 Fig); the large quantity of deactivated yeast present in the cornmeal-molasses food, or other nutritional differences between the two substrates [82], may have enhanced ovary development in outdoor flies fed standard food. However, we cannot rule out additional differences between indoor and outdoor cages since density and age of sampled flies were not controlled in the outdoor cages. Nonetheless, the fruit-fed cages, which more closely resemble the environment of wild flies in an orchard or compost pile, may more accurately represent the natural state of investment in reproduction in the wild; rich laboratory fly medium may permit more oogenesis than normally occurs in nature [83–86].

To test the hypothesis that evolved genetic changes in the outdoor populations also contributed to increased diapause propensity in early winter, we captured seasonal samples from outdoor cages and reared them in the lab for two generations (“G2”). We exposed the G2 offspring to diapause-inducing conditions to perform a common-garden assay of diapause in flies that evolved under natural conditions. We concurrently sampled flies from our indoor hybrid swarm cages and bred them under identical conditions as negative controls. We found that the outdoor populations evolved a significantly increased propensity for diapause in late fall for both diapause phenotypes, whereas the indoor populations experienced few significant changes in diapause propensity (Fig 6B; C). This result was observed in flies held at two different diapause-inducing temperatures (10.5 °C and 12 °C), and the changes in diapause incidence were stronger for stage 10 diapause (Fig 6C) relative to stage 8 (Fig 6B). We note that both indoor and outdoor cages experienced parallel changes in diapause, likely due to experimental variation between cohorts. However, the magnitude of change from the first collection to the final collection was significant for outdoor cages (*P* < 0.05 for three of four tests, Fig6B; C) but not indoor cages (*P* > 0.05 for all tests). The evolved change in diapause corresponds to the onset in mid-November of sustained cold conditions at our field site (Fig 6D), suggesting that these conditions may have caused selection for individuals able to diapause. We note that the results of these field experiments may have been influenced by potential invasion of wild flies into our field cages, which would change the genetic composition of the *D. melanogaster* populations. Based on the ratio of *D. simulans* to *D. melanogaster* at nearby Carter Mountain Orchard in Charlottesville, VA, we estimate that at most 5% of *D. melanogaster* in the cages were invaders (see Methods). Collectively, our field results suggest that environmental conditions produce plastic changes in ovary development and also result in selection for increased propensity to diapause.

### Diapause-associated SNPs are present at high frequencies in Zambia

Diapause has been proposed to be a recent adaptation to cold in temperate populations of *D. melanogaster* [12,24,87], but see [35, 88]. To test whether the diapause-associated alleles mapped here are present in ancestral populations, we first examined the median allele frequency of pro-diapause alleles in a large sample of flies from Zambia [89, 90]. We predicted that pro-diapause alleles would have lower allele frequencies in Zambia than the pro-diapause alleles identified in permutations. We found that the actual median allele frequency of pro-diapause alleles generally falls within the expected range of allele frequencies based on permutations for both phenotypes when considering all SNPs and LASSO SNPs (Fig 8). However, when considering the top 0.01% or 0.1% of SNPs in the GWAS, we found that pro-diapause alleles for diapause at stage 8 are more common than expected by chance in Zambia. This trend was not observed for the stage 10 phenotype; the allele frequencies for these SNPs fell within the expected range based on permutation. The high allele frequencies of stage 8 diapause SNPs is not due to an excess of diapause-associated SNPs in tracts of European admixture in Zambian flies (S16 Fig). The abundance of pro-diapause alleles suggests they are longstanding polymorphisms that are not deleterious, and may even be favorable, in ancestral climates.

**Fig. 8.**
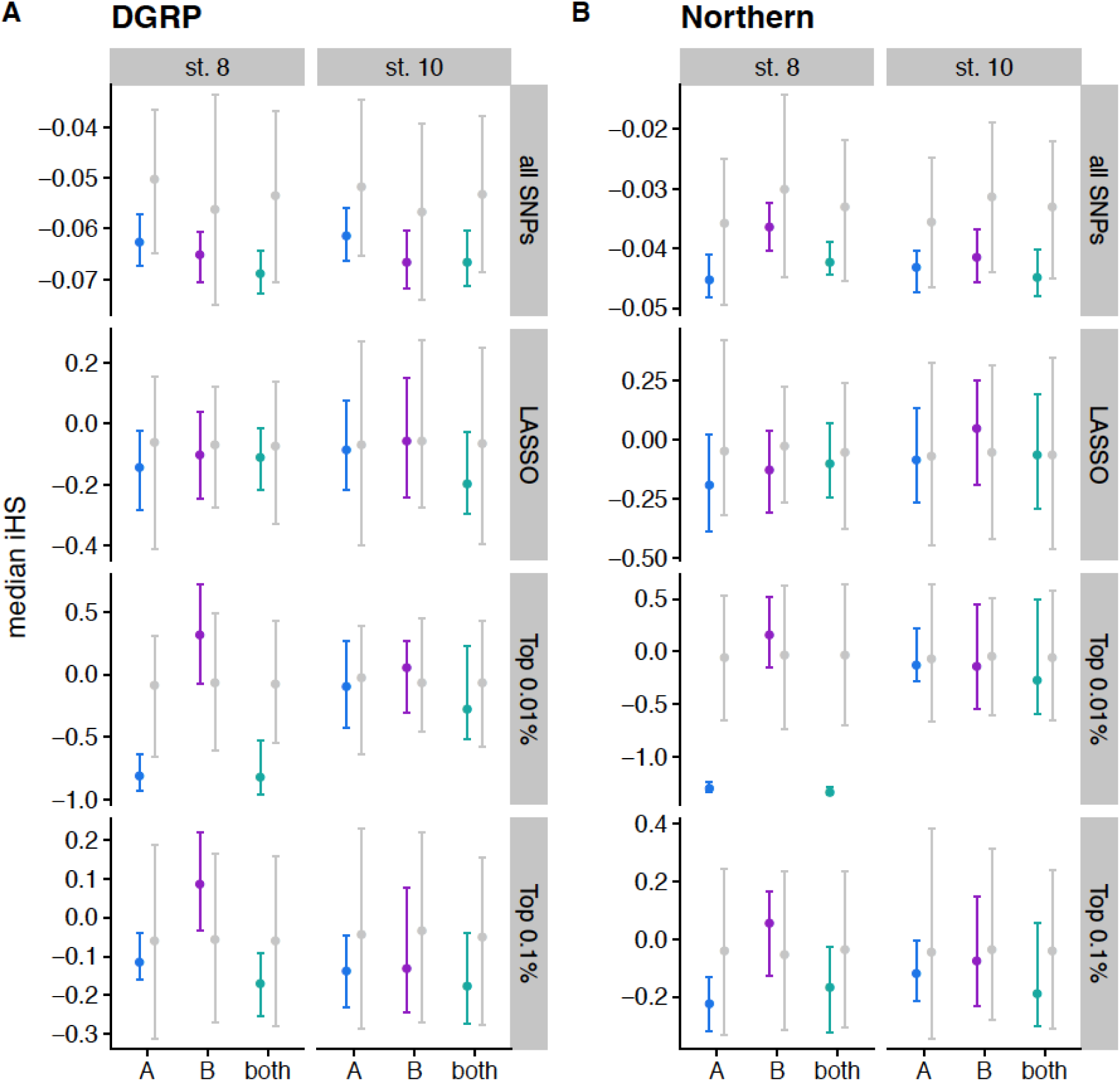
Integrated haplotype homozygosity score (iHS) of diapause-associated SNPs in two populations. (A) iHS was calculated for every SNP in the DGRP. For each GWAS, the median iHS was calculated for all SNPs, the LASSO SNPs, the top 0.01% of SNPs in the GWAS, and the top 0.1% of SNPs in the GWAS for each phenotype. Points represent the median; error bars extend to the 2.5% to 97.5% quantiles. Colored bars are 100 imputations of the original data. Grey bars are 100 permutations for populations A and B; 1000 permutations for both. (B) Same as panel A, but using a set of 205 lines collected from Pennsylvania and Maine (“Northern”).

### Evidence for partial sweeps of anti-diapause alleles in North America

We tested for signals of partial selective sweeps on pro-diapause alleles in North America. We used the DGRP, from Raleigh, North Carolina, [77] and a set of sequenced, inbred lines collected from Pennsylvania and Maine (“Northern” lines) to calculate the integrated haplotype homozygosity score (iHS) [91] for the pro-diapause alleles identified in the GWAS. In this test, an elevated iHS would suggest that the pro-diapause allele is found on a longer shared haplotype, suggesting a recent sweep, whereas depressed iHS would suggest a sweep of the anti-diapause allele. Median iHS of both stage 8 and stage 10 diapause-associated SNPs was generally weakly depressed relative to permutations (except for LASSO SNPs, Fig 8B), suggesting the possibility of partial selective sweeps that increased the frequency of anti-diapause alleles in both the DGRP and Northern populations. This result is consistent with the relatively high frequency of pro-diapause alleles observed in Zambia.

## Discussion

### The genetic basis of an ecologically relevant trait identified from mapping in a hybrid swarm

Herein, we estimated the genetic architecture of natural variation in ovarian diapause in *D. melanogaster* using a novel hybrid-swarm based mapping strategy. Our work changes the interpretation of the genetic basis of variation in diapause in *D. melanogaster*, which has previously been characterized as being under the control of a small number of loci with large effect using QTL mapping and candidate gene approaches [27,52,60,76]. We studied spatiotemporal variation of small-effect, diapause-associated alleles in global *D. melanogaster* populations, finding evidence that both pro- and anti-diapause alleles have experienced selection in different geographic regions and seasons.

The SNPs we identify, which are generally unremarkable in a genomic context (Fig 5; Fig 7; Fig 8), are evidence of “gold dust” for an ecologically relevant trait: alleles that are likely critical to evolutionary processes but nonetheless have neither large phenotypic effects nor dramatic genomic signatures of selection [92]. Our analysis of clinal variation in diapause-associated SNPs is evidence that the accumulation of subtle allele frequency differences in many SNPs with slight phenotypic effects may produce phenotypic differences across broad geographic scales. Whether spatial and temporal selection on these SNPs is due to selection for diapause or selection for correlated traits with a shared genetic basis remains unknown. However, collectively these unexceptional polymorphisms can contribute to ecologically important variation. The lack of strong genomic signals in diapause-associated SNPs relative to randomly identified SNPs suggests that perhaps a large fraction of polymorphisms in *D. melanogaster* could play roles in other complex traits [93] that may have experienced varying degrees of spatiotemporal selection, recent partial sweeps, and balancing selection. If many SNPs underlie ecologically relevant phenotypic variation, strong deviations from the genomic background of patterns of diversity will be the exception, not the norm. More studies of the evolutionary history of alleles underlying complex traits will be required to determine whether the pattern we observe for diapause is a general phenomenon.

**Fig 7.**
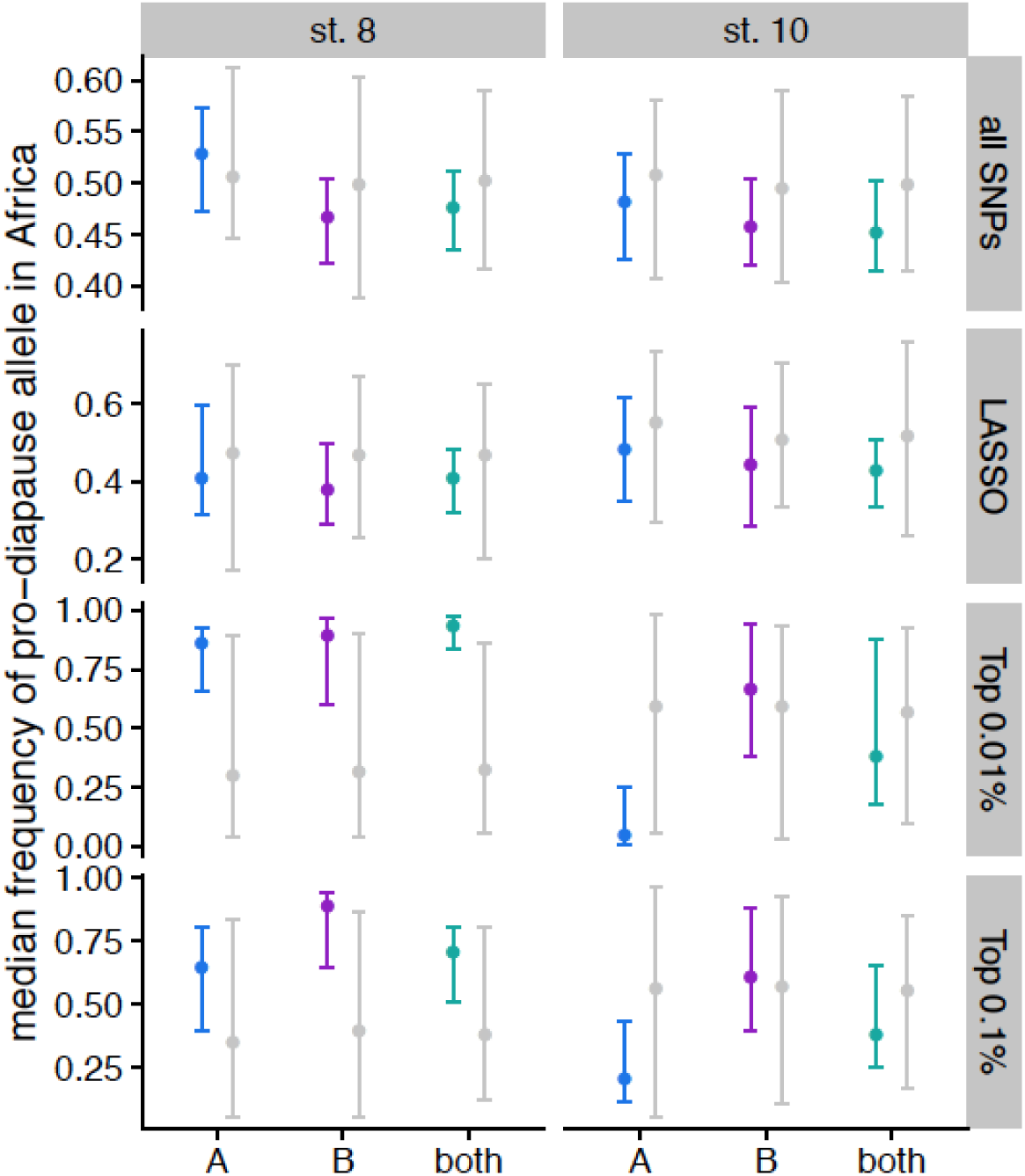
Diapause-associated alleles are common in Zambia. The median allele frequency of pro-diapause alleles for all SNPs in the GWAS, LASSO SNPs, the top 0.01% of the GWAS, and the top 0.1% of the GWAS for stage 8 (left) and stage 10 (right). Points represent median of the imputations/permutations; error bars show 2.5% to 97.5% quantiles. Grey points/bars are 100 permutations for populations A and B; 1000 permutations for both. Colored points/bars are 100 imputations of the original data.

### Modest heritability and strong environmental control of diapause

We find that the heritability of diapause is low but detectable (Fig 3), and we demonstrate that this heritable variation may be subject to selection under field conditions (Fig 6). We note that our estimates of individual effects of SNPs are likely underestimates due to the inclusion of a genome-wide genetic relatedness matrix in our model, which would partially correct away the effect of any given SNP. Like other studies [24,35,36,94], we identify temperature and nutritional exposure as two critical variables that influence diapause and extend this analysis to outbred lab and field-caught samples. Specifically, we find that diapause induction is exquisitely sensitive to extremely small differences in temperature (Fig 2), emphasizing the importance of precise thermal control and monitoring for experiments studying diapause in this species [95]. This fine-grained sensitivity suggests that even temperature variation across shelves of an incubator or at different distances from a light source could potentially influence experimental outcomes; these potential sources of thermal variation were controlled in our experiment by our use of multiple environmental chambers with nearly identical physical setup aside from temperature and photoperiod.

Our finding that diapause was not induced by short day length (but rather by long days in our experiment, Fig 2) is somewhat consistent with other studies that failed to find an effect of photoperiod on diapause in *D. melanogaster* [94, 95]. However, some recent studies do observe short-day photoperiodic responses in diapause [88, 96]. One study suggested that light conditions which mimic the natural daily variation of solar radiation can in fact generate a photoperiodic response [97]. In our study, short days under rectangular light cycles with standard white LED light did not increase diapause induction. The slightly increased diapause incidence we observe at longer photoperiods is surprising and warrants future study. Because we were able to account for temperature variation on a precise scale, we can rule out the potential of light-mediated temperature differences. Future experiments should work to disentangle the roles of temperature, wavelength and variable light intensity on photoperiodism in *D. melanogaster*.

### Different genetic signals underlie unique diapause phenotypes

Throughout our results, we observe different genetic signals underlying diapause scored at two different developmental stages. Diapause at stage 8 has been used as the phenotype for most studies of diapause [15,18,19,52]; however, others have argued that stage 10 is a more biologically relevant phenotype [35]. Stage 8 marks the onset of vitellogenesis [66], whereas stage 10 requires clearing a major checkpoint in the ovarian development program before proceeding to yolk-demanding stages [98]. We find an enrichment of SNPs shared between the two phenotypes, but the population genetic signals of diapause SNPs tend to be stronger for the stage 8 phenotype. The SNPs underlying stage 8 diapause vary more predictably clinally (Fig 5), are more likely to show seasonal variation (S13 Fig), are at higher frequency in Zambian populations (Fig 7), and show stronger potential selection on anti-diapause alleles (Fig 8). However, we observed more dramatic seasonal variation in the stage 10 phenotype in our field study. Combined, these results suggest that the two phenotypes may be influenced by both different genetic architectures and different environmental cues, and therefore may respond differently to temporally or geographically varying selection. We also note that the genetic makeup of the mapping population appears to be critically important to the identification of SNPs associated with a polygenic trait like diapause, consistent with quantitative genetic variation in diapause being influenced by rare variants.

### Weak clinal signal in alleles underlying a strongly clinal phenotype

Diapause shows strong clinality in both North America [96] and Australia [67], with flies collected at higher latitudes showing higher diapause incidence in common gardens. This trend has not been seen in Europe, perhaps due to the high frequency in southern Europe of a pro-diapause allele that recently arose in Italy [99]. Collective effect of many weakly clinal diapause-associated alleles is sufficient to produce signal that is consistent with clinal variation in phenotype (Fig 5). However, this generally weak trend suggests those SNPs with the strongest clinal signal may not always be the most ecologically relevant SNPs, at least for some phenotypes. Furthermore, this signal was not seen when examining unlinked LASSO SNPs, suggesting that some linkage among top diapause-associated SNPs could potentially contribute to the observed clinal signal.

Our evidence of multilocus clinal selection for diapause adds to a growing literature showing evidence of spatial differentiation of loci underlying complex traits in natural populations. For example, in *D. melanogaster*, clinal SNPs are also enriched for SNPs influencing cuticular hydrocarbon content [100]. In gypsy moths, SNPs associated with three ecologically relevant traits have consistent clinal signals [101], and in corals, SNPs associated with heat tolerance are more common in warmer reefs and in warmer microclimates within a single reef [102]. Drought-associated SNPs also show spatial variation in European populations of *Arabidopsis thaliana* [103], and models suggest that those populations with more drought-tolerance alleles may adapt to global warming more effectively. Therefore, in addition to elucidating genetic mechanisms of local adaptation, the ability to detect polygenic adaptation of ecologically relevant traits over space is important for predicting population-level changes in response to climate change.

### Seasonal evolution of diapause may be idiosyncratic

We identified several pieces of evidence for stochastic or unpredictable seasonal variation in diapause. We observed a genetic shift towards higher diapause in the late fall in our field experiment (Fig 6) but saw little evidence for repeatable seasonal variation in pro-diapause alleles across multiple years (Fig 5) or multiple populations (S12 Fig). The increase in diapause in late fall may not solely be due to selection on the diapause trait itself; flies that are able to diapause are also more tolerant of cold [15], which could potentially be the phenotype under selection. If the ability to diapause makes flies more tolerant of cold, or if diapause and cold tolerance are influenced by some of the same alleles, then selection for cold tolerance in late fall could explain the increase in diapause. We also found that some individual populations show signals of seasonal variation in diapause-associated SNPs (S12-S13 Fig), though the direction of this variation is often the opposite of what we would predict, with pro-diapause alleles more common in the fall.

We propose three possible explanations for the absence of repeatable seasonal signals in diapause-associated SNPs. First, we note that the seasonality of diapause has only been tested in mid-latitudes [19 and this study]; more southern or northern environments may not produce strong seasonal variation in diapause or other life history traits. In this case, we would not expect to see repeatable seasonal changes in diapause-associated SNPs across diverse populations. However, two of the populations in which we observed concordant patterns of allele frequency changes in diapause-associated alleles were Georgia and Massachusetts (S13-S14 Fig). This observation suggests that higher and lower latitude populations may also undergo adaptive seasonal changes in diapause. We also note that the design of our mapping experiment may have given us greater power to detect clinal signal relative to seasonal signal, due to the wide geographic spread of our starting lines and relatively few lines of known seasonal origin.

Second, the variants associated with diapause may also be favorable for stress resistance during seasons other than winter, resulting in a lack of strong spring/fall differentiation. This possibility is supported by the presence and even excess of pro-diapause alleles in central African populations, which experience seasonal dry periods and food limitation as well as cooler temperatures at high elevations [104]. To our knowledge, this study is the first to examine diapause status in field-caught flies across seasons. We observed surprisingly high incidence of diapause-like ovaries in outdoor cage-reared flies over the course of the summer and fall. While high diapause incidence was observed in December, we also observed increasing diapause throughout the summer and fall, followed by a dramatic drop in November. This finding suggests that diapause-like ovaries may be a response to general stress conditions. Indeed, starvation, heat stress, and crowding are known to cause arrest of oocyte maturation and degradation of pre-stage 10 egg chambers [83–85,105,106]. Stochastic seasonal phenotypic evolution has been documented in stick insects (*Timema cristinae*). In *Timema*, the evolution of the stripe pattern, which is under strong frequency-dependent selection, is highly predictable across seasons, whereas the evolution of color, which is under diverse selective pressures, varies unpredictably [107]. Diapause and its underlying SNPs may be subject to more complex patterns of selection due to pleiotropy and environmental plasticity, resulting in a general lack of predictably across populations and years. This possibility is further confounded by the limitations of endpoint-based sampling. While the populations were broadly sampled in “spring” and “fall”, the particular selective pressures experienced by flies in each sample may have varied dramatically between locations and years.

Lastly, seasonal evolution of diapause may require only a subset of the variants we identified, resulting in a lack of broad-scale signal across multiple sampling populations or years. This possibility could occur because only a fraction of the sites are sufficient to induce adaptive changes, or could possibly be due to false-positives and limited statistical power in both the GWAS reported here and in the seasonal surveys [53]. The seasonal model employed by [78] prioritizes SNPs that change in the same direction across many populations. The pro-diapause variants selected in different populations may differ from year to year, resulting in a lack of signal when comparing our large collection of diapause-associated variants to those SNPs with repeatable seasonal changes in allele frequency. We find evidence for this possibility in our analysis of individual populations, which revealed consistent changes in the frequency of diapause associated alleles in some, but not all, populations (S13-14 Fig). Likewise, polygenic adaptation to thermal regimes in *Drosophila* can use different combinations of genetic loci, with replicate populations evolving frequency changes in only subsets of the same alleles when subject to the same selective conditions [108]. However, we did observe selection for diapausing genotypes in our field data, with flies sampled in late November showing an increased genetic propensity for diapause. Given the observed phenotypic shift, we hypothesize that sequencing the late November samples would show an increase in frequency of at least some pro-diapause alleles relative to samples collected earlier in the season.

Taken together, our results confirm that North American populations carry heritable and selectable genetic variation for diapause, but the genetic basis of seasonal selection on this trait may not be repeatable from year to year or population to population. While strong evidence exists for seasonal variation in the genetic composition of local *D. melanogaster* populations based on pooled sequencing [53, 78], the loci underlying a single quantitative trait likely do not capture the complexity of the selective forces driving these allele frequency changes. Interactions between variable environments, polygenic traits, and pleiotropic alleles may result in a lack of broad-scale seasonal signal in diapause-associated SNPs.

### Diapause may be an ancestral adaptation coopted for cold

Our results on the evolutionary history of pro-diapause alleles are consistent with recent work suggesting that diapause is in fact an ancient, polymorphic adaptation in *Drosophila* [88]. While two studies [12, 87] observed negligible diapause incidence in African lines of *Drosophila*, these studies classified entire isofemale lines as diapausing or non-diapausing based on at least 50% of individuals having ovarioles developed to stage 8 [24]. Therefore, it is possible that some individuals were in diapause even if the entire line was classified as non-diapausing. More recent studies have documented quantitative variation in diapause induction among various African lines of *D. melanogaste*r [35, 88], as well as other related species [88], using a stage 10 cutoff, but these studies did not rule out potential introgression of European haplotypes. We find that alleles promoting diapause are as common, or more common, than expected by chance in Zambia. Taken together, our genetic data and previously published phenotypic results suggest that diapause at either stage may in fact be an ancestral trait that is also polymorphic in related species.

Diapause in African flies may be related to cold temperatures at mid to high elevations [109, 110], or to seasonal stressors other than cold, including wet-dry seasons that influence food availability. Ancestral populations of *D. melanogaster* may have subsisted on marula, which only fruits for part of the year [104]. As a result, dry conditions or seasonal lack of food availability may have required intermittent diapause or estivation. Our data on diapause-associated SNPs in central Africa support this hypothesis. Interestingly, our data are also consistent with the observation by Bergland *et al* (2014) that summer-favored alleles of seasonally varying SNPs are more likely to be rare in Africa. Further, the Zambian data also consistent with the evidence of partial selective sweeps of anti-diapause alleles in North Carolina and the Northeast. Collectively, these data suggest diapause may in fact be favored in Zambia, and the ability to *not* diapause may be favored, at times, in temperate latitudes of North America. Our findings support a model in which *D. melanogaster* is highly opportunistic, taking advantage of favorable conditions at any point in the year but also restricting investment in reproduction when conditions are unfavorable for offspring survival, whether due to cold, desiccation, starvation, or other stressors.

## Conclusions

Understanding the genetic basis of local adaptation has been a major goal for evolutionary biology in the genomic era [111, 112]. Studying this question in *D. melanogaster*, which exhibits dramatic phenotypic variation across space and time and offers extensive population genomic resources, allows for a thorough exploration of the evolutionary history of alleles underlying adaptive phenotypes. Diapause is a particularly valuable trait to dissect at a genome-wide level because it underlies demonstrable life history tradeoffs and is relevant to predicting insect responses to changing climate conditions [113]. The genetic basis of variation in diapause induction has been investigated in numerous other species and ranges from a single underlying locus in *D. littoralis,* flesh flies and linden bugs [48,114,115], to a few loci of a small effect in the face fly and European corn borer [116–118], to many loci in mosquitoes and the speckled wood butterfly [119, 120]. With the exception of the wood butterflies, most of these studies estimated the number of loci using traditional quantitative genetics approaches; our study is unique in its estimation of the genome-wide genetic architecture of diapause via association mapping. The association mapping performed here indicates that ancestral variation in this highly polygenic trait contributes in unique ways to spatial and temporal variation in common selection pressures associated with life in temperate environments. Whether this result generalizes to other life-history, behavioral, and physiological traits that underlie local adaptation across time and space remains an open question.

Compared to many insects with hard-programmed diapauses that are triggered by photoperiod and last for months or even years [9], ovarian dormancy in *D. melanogaster* is short-lived, readily reversible, and highly dependent on present environmental conditions. These characteristics of diapause in *D. melanogaster* may be a result of the genetic architecture of the trait, with hundreds or even thousands of SNPs slightly modulating an individual’s propensity to diapause under a variety of unfavorable conditions. These loci vary somewhat predictably across space but not across time; nonetheless populations do evolve an increased propensity for diapause in unfavorable conditions, perhaps via selection of unique allelic combinations in different populations, or selection for linked traits. On a broad geographic scale, clinal differentiation of thousands of alleles of small effect results in cumulative genetic effects that produce latitudinal phenotypic variation. Despite similar patterns of diapause variation over latitudes and seasons in North America, we conclude that local adaptation relies on different patterns of selection in space and time to optimize performance in variable conditions.

## Materials and Methods

### Construction of custom photoperiod chambers

We built environmental chambers (S17A-B Fig) from custom-cut opaque black plastic (Quality Machine Service, Waynesboro, VA). Design files for the machining of each wall are available upon request. Each chamber was controlled by a Raspberry Pi Model 3 computer with a static IP address and its own GitHub account running custom Python scripts (see https://github.com/bergland-rpi/rpi-02 for an example of scripts). To improve air circulation and prevent temperature changes associated with lights, four 92 mm computer fans (OutletPC.com) were mounted behind light-proof circular vents (4” diameter darkroom vents, midgetlouver.com). The two fans on the sides blew air in from the outside, and fans on the back and top pulled air out to create constant airflow and reduce the warming produced by lights. A custom circuit board (S17C Fig) was designed with Fritzing software (fritzing.org). The following electronic components were directly soldered to each circuit board: TSL2561 Digital Luminosity detector, SHT-31D Temperature and Humidity Sensor, MCP9808 Temperature sensor, and 74AHCT125 Quad Level Shifter. A 24-LED RGBW Natural White NeoPixel ring was connected to the circuit board with 22 AWG wire. After the initial mapping experiments, the lights were replaced with a 64-LED NeoPixel grid. All electronic components were purchased from www.adafruit.com. A custom Python script turned lights on and off for fixed photoperiods, while recording temperature, light intensity, and humidity every 60 seconds. After the initial mapping experiments, the LED lights were replaced with 64-LED RGBW Natural white NeoPixel grids for future experiments. The TSL2561 sensor was used to ensure constant light intensity across boxes for all experiments following the initial mapping.

The boxes were housed in a cold room held at 10°C (S17A Fig). However, we noticed that there was spatial variation in the actual room temperature, and we exploited this variation to expose flies to a broad range of temperatures (S3 Fig). To further modulate temperature, some boxes were outfitted with ZooMed ReptiTherm habitat heaters in one of three sizes (6×8”, 8×12”, or 8×18”). We found that these heaters increased the air temperature in the chambers by approximately 0.5, 1, and 1.5 °C respectively. We placed all fly vials horizontally on wire racks elevated 3 cm above the surface of the heaters to prevent directly warming the flies or their vials. We calibrated the temperature readings in each box by manually recording the temperature at the position of the fly vials using a high accuracy thermometer (Model EL-WIFI-DTP+; www.dataq.com) and offsetting the recorded temperatures based on this standardized reading. Quality control checks of our environmental data revealed that all chambers had consistent temperatures throughout the course of the experiment (S3A Fig) and that diurnal temperature fluctuations were limited to approximately 0.25°C (S3B-C Fig).

### Hybrid swarm construction and collection of individuals for phenotyping

Seventy inbred or isofemale lines spanning seven geographic/seasonal collections (S1 Table) were chosen to initiate the hybrid swarm: Rocky Ridge Orchard, Bowdoin, Maine [NCBI BioProject #PRJNA383555]; Ithaca, New York [121]; June and October collections from Linvilla Orchard, Media, Pennsylvania [122]; the Drosophila Genetic Reference Panel from Raleigh, North Carolina [77]; the Southeastern US [123], and the Bahamas [123]. Ten inbred/isofemale lines were chosen at random from each collection, and five lines were randomly assigned to each of two hybrid swarms. We reassigned approximately five lines in an attempt to balance cosmopolitan inversion frequencies across the two hybrid swarms (see “Inversion genotyping” below). Two lines (24,2 and 12LN6-24) produced an insufficient number of offspring to generate F1 crosses, so they were eliminated, and each population was initiated with 34 lines. See S1 Table for complete information about founding lines.

For each population, a total of 4 sets of 34 round-robin crosses were initiated, so each line appeared in 8 out of 136 total crosses (4 crosses used males from each line, 4 additional crosses used the females). The order of crosses was randomly generated. Fifteen virgin females and 10 males were placed in a yeasted bottle of cornmeal-molasses medium and allowed 72 hours to lay eggs. One day prior to the eclosion of the first adults, the open bottles were placed in a 2m x 2m x 2m cage (Bioquip product # 1406C). The flies were given ∼5 days to eclose, and then 20 trays of fresh yeasted cornmeal-molasses food (∼800 mL media in a 23 cm x 23 cm tray) were provided on day 14 to each cage to collect eggs for 48 hours. The trays were incubated at 25°C in a 12:12 light: dark light cycle, 50% relative humidity. After eclosion, 8 of 20 trays were reintroduced to the cage. For the F3, F4, and F5 generations, 20 trays were provided for 24 hours for egg laying. After egg collection, the food was removed and covered and incubated at 25 C in a 12:12 light cycle. At the F6 generation, the population size in the cages was reduced by adding only 10 food trays for 16-24 hours, and the cages were maintained in this manner for all future generations.

To collect individuals for the GWAS, yeasted bottles containing 35 mL of cornmeal-molasses media were placed in each cage for 3-6 hours to collect F4 and F5 generations. Offspring were incubated for 9-10 days at 25°C on a 12:12 light cycle. Female flies were collected under light CO_2_ anesthesia within 2 hours of eclosion (as indicated by the presence of folded wings, enlarged/pale abdomen and/or meconium). 20 flies were placed in a vial containing 10 mL of cornmeal-molasses media, and flies were placed into temperature-controlled chambers within 1 hour of collection. Flies were distributed to one of 47 chambers assigned to a photoperiod of 9, 11, 13, or 15 hours of light and a temperature varying between ∼10 and 15 °C (S3 Fig). After 13-15 days, the flies were transferred to fresh food. The flies were snap frozen in liquid nitrogen after exactly 28 days and stored at −80 °C.

### Yeast supplementation experiment

We collected virgin female flies from the F20 generation of each cage as described above and placed vials in 9L:15D light cycles at 5 different temperatures. Half of the vials were supplemented with a sprinkle of live baker’s yeast, and the other half received no yeast. After four weeks, flies were snap frozen.

### Phenotyping

Flies were thawed in 70% ethanol, then transferred to a ∼100 µL droplet of phosphate buffered saline for ovary dissection. Two aspects of ovary development were scored. First, the most advanced, non-egg stage of ovariole observed was recorded. We recorded stages from 6 (or less) to 11; stages 12-13 were combined into stage 11 [66]. Second, the number of fully formed eggs was counted (counted from 0-15 or scored as 15+ if more than 15 eggs were present). These phenotypes were later reduced to three binary phenotypes: diapause at stage 8 (no ovarioles observed past stage 7), diapause stage 10 (no ovarioles observed past stage 9), or diapause at stage 14 (no mature, stage 14 eggs present).

We analyzed the influence of environmental variables (temperature, photoperiod, generation, population and *Wolbachia* status (see below) using binomial generalized linear models. We used the R package *car* [124] to estimate the sum of squares explained by each variable with function *Anova()*, and then divided by the total sum of squares to estimate percent variation explained (PVE).

### DNA extraction and library preparation

After dissection, fly carcasses were placed in DNA lysis buffer (Agencourt) in a 96 well deep well plate. Forceps were cleaned with ethanol between each individual dissection to prevent DNA contamination. After completing a 96 well plate, the carcasses were lysed in a Qiagen TissueLyser using four 2-millimeter stainless steel beads. The lysates were spun down, transferred to a fresh 96 well plate, and frozen at - 80 °C until further processing (see DNA extraction below). Two randomly chosen blank wells were left on each 96 well plate to verify plate identity and orientation.

DNA was extracted from 20-30 pooled flies (parental strains) or individual flies (hybrid swarm offspring) using the DNAdvance kit (Agencourt) in 96 well plates according to manufacturer’s instructions. In total, we genotyped 2,823 individual flies. An RNase treatment was added between wash 1 and wash 2, and the DNA was bound with fresh beads following the RNase treatment. Sequencing libraries were prepared using a modified Nextera protocol that permits low volumes and DNA concentrations [125]. Briefly, DNA was quantified with a Picogreen assay and normalized to 1 ng/µL with a liquid handling robot. One ng of DNA was used for library preparation with unique barcode combinations for every sample (barcodes available in supplemental file on DataDryad). For the parental lines, individual libraries were pooled to obtain equal concentrations of each line. They were sequenced in a single lane of HiSeqX with paired-end, 150 bp reads at the Hudson Alpha Institute for Biotechnology. Ten 96-well libraries of hybrid swarm individuals were pooled for each lane of sequencing so that ∼940 flies and 20 blanks were combined per lane. Library sizes and quality were verified by Bioanalyzer. Hybrid swarm lanes were sequenced with Illumina HiSeq3000 paired-end, 150 bp reads at the Oklahoma Medical Research Foundation sequencing center.

### SNP calling in parental genomes

We resequenced the genomes of all 68 parental lines to confirm their identities and compared the sequences to published sequences (see S1 Table for SRA accessions of founding line sequences). Following read merging with *PEAR* [126] and read mapping to the *Drosophila* genome version R5/dm3 with the *mem* algorithm in *bwa* [127], we called preliminary SNPs with *GATK*’s *Unified Genotyper* [128]. We found that one line (12LN6_41_B47) did not match previous sequencing, and two other lines (20,17 and 20,28) appeared to be partially contaminated (some chromosomes were accurate, others had discrepancies from previous sequencing). For these lines, we used our resequencing data as the only parental genome sequence; for the other lines, we combined our new sequence data with existing data for higher depth coverage. Individual gVCF files were generated with *HaplotypeCaller* in GATK [128] and then combined into a single parental gVCF with *CombineGVCFs*. We used randomly sampled known SNPs from the DGRP to calibrate the SNP calls with *VariantRecalibrator* and *ApplyRecalibration*. We filtered down to two sets of SNPs; one conservative set of ∼1 million SNPs with the highest quality (ts_filter_level=99) and a second more extensive set of ∼3.1 million SNPs with a broader range of SNP quality (ts_filter_level=99.9). The former set of more high-confidence SNPs was used for genome reconstruction; the latter set was further filtered as described below and used for the GWAS to allow us to test more SNPs for trait association. For reconstruction purposes, heterozygous sites in the parental lines were treated as missing data.

### *Wolbachia* status

To determine whether inbred lines and hybrids were infected with *Wolbachia*, we counted the number of sequencing reads mapping to the *Wolbachia* genome (which was part of our reference genome) and divided that number by the total number of reads mapping to chromosomes 2L, 2R, 3L, 3R, and X. This ratio produced a clearly bimodal distribution, with individuals falling into two groups: those with a ratio of substantially less than 1:1000 (0.001) and those with a ratio greater than 1:1000. We classified all individuals in the former group as *Wolbachia*-negative, and the latter individuals as *Wolbachia*-positive.

### Hybrid swarm genome reconstruction

Paired end reads were merged and mapped as described above. Bam files were then processed through our in-house genome reconstruction pipeline [63] Briefly, polymorphic reads were counted with *ASEReadCounter* in *GATK* [128] with a minimum mapping quality score of 10. *HARP* [129] was used to preliminarily call parental haplotypes. The top 14 possible founders were then used for precise genome reconstruction in *RABBIT* [130]. The output of *RABBIT* was translated to phased diploid genotypes using a custom script [63]. All lines included in the founding populations were recovered following reconstruction of the F4 and F5 generations of the hybrid swarm (S18 Fig). The median proportion of the genome derived from a line was 2.7% (range = 0.5% - 9%; expected = 1/34 = 2.9%). Of over 800,000 private SNPs in the founding lines, only 109 SNPs were lost in the hybrid swarm, suggesting that virtually all founding haplotypes were recovered at least once in the sequenced hybrid swarm.

### Recombination simulation and accuracy calculations

We generated hybrid swarms *in silico* using the genotypes of our founding lines and the pipeline described in Weller and Bergland 2019 [63]. Genome sequences for each parental line were generated with *FastaAlternateReferenceMaker* in *GATK* [128] with flag *use_IUPAC_sample* to generate ambiguous bases at heterozygous sites. We simulated F4 and F5 generations for populations A and B based on recombination rates in [131] (S19 Fig). For each population and generation, we constructed 100 replicate populations of 10,000 individuals and randomly chose 5 individuals from each population. Simulated reads at 0.5X coverage for recombinant individuals were generated with *wgsim* (https://github.com/lh3/wgsim). These reads were mapped to dm3, and the simulated hybrid genomes were reconstructed as described above. To determine the accuracy of reconstruction, the known genotypes of the simulated individual were compared to the reconstructed genotypes, and the proportion of sites with identical genotypes was calculated. Sites that were missing or heterozygous in the actual founder genotypes were excluded from the accuracy calculation. We found that for both sets of founders, the majority of individuals were > 99.9% accurate (S20 Fig). We therefore predict similar levels of accuracy in our actual sequencing data.

### Masking regions with poor reconstruction quality

Based on the simulations described above, we found that small genomic segments (<1 Mb) from parental haplotypes were over-represented in our reconstructed data (see S19 Fig). Haplotypes less than 1 Mb made up ∼4% of all simulated genome haplotypes but made up ∼20% of all reconstructed haplotypes. Therefore, any reconstructed segment less than 1Mb was masked from our genotype calls as missing data. To count the number of recombination events, any consecutive short (< 1Mb) paths were grouped together as an “unknown” parental haplotype, and that single unknown path was counted as a parental segment for recombination purposes. If a short path was surrounded by the same parental haplotype on either side, we “bridged” over it and did not count it as an additional recombination event. Following this clean up step, visual inspection revealed that the resulting genome reconstructions from both simulated and actual sequencing data more closely matched the recombination rate and haplotype size predicted by our recombination simulator (S19 Fig); however, the cleaned-up reconstructions still significantly differed from the simulated distributions of haplotype size and recombination count (Kolmogorov-Smirnov test, *P* < 8 x 10^-6^ for all tests). This discrepancy occurred in both the reconstruction of simulated data as well as the reconstruction of actual data.

### Imputation of missing data

Because the *GENESIS* software package used for the GWAS requires a complete dataset and our individuals were drawn from two populations with unique genetic composition, we performed a custom imputation of missing data within each population separately. The imputation required two steps. First, for any site at which the parental line was heterozygous, we randomly chose one allele for each offspring genotype. After this step was performed, the remaining missing data (due to missed calls in the parental genotyping or masked short segments in the reconstruction) were imputed by taking the most likely genotype based on Hardy-Weinberg allele frequencies. This step was calculated within each hybrid population to accurately capture allele frequency differences between the two populations. We repeated this random-choice followed by Hardy-Weinberg imputation algorithm 100 times to create 100 uniquely imputed genotype sets. All analyses were carried out on all imputed data sets, and the range of values for imputations are presented for transparency of the variation that this imputation introduces. The Hardy-Weinberg imputation procedure was also used to impute allele frequencies for the population genetic analysis (see below).

### SNP filtering prior to association mapping

We constructed two SNP filters for mapping in our hybrid swarm population. The initial results of variant calling (see above) contained ∼3.1 million SNPs. We filtered this set of SNPs with two additional filters prior to association mapping. The first filter was a strict quality control filter to identify SNPs to be used for construction of the genetic relatedness matrix (GRM). The second filter was slightly less stringent to produce a larger set of SNPs for mapping. For the strict quality control filter, we excluded SNPs that had > 10% of genotypes replaced by our imputation algorithm, an F_ST_ > 0.2 between the two populations (as calculated by *snpgdsHWE()* in *SNPRelate*), SNPs that were fixed in one population (but not the other), and SNPs with a Hardy Weinberg Equilibrium *P*-value of <10^-20^ in either population alone or the two populations combined. This filtering resulted in ∼1.2 million SNPs used for GRM construction. The less stringent filter was the same as above, but allowed SNPs that were fixed in one population but not the other to remain for the GWAS. This filter resulted in ∼2.2 million SNPs. After mapping, SNPs were further filtered for minor allele frequency > 0.05 in downstream analyses.

### Analysis of genetic structure of hybrid populations and parents

Principal components were calculated with the function *snpgdsPCA()* in the R package *SNPRelate* [132]. Identity by state calculations were performed with *snpgdsIBS().* Karyotypes for each chromosome arm were inferred based on diagnostic SNPs identified in [133]. Principal component analysis of reconstructed genotypes revealed strong differentiation between populations A and B (S2 Fig), which was also evident in the identity by state genetic relatedness matrix of all individuals (S21 Fig). Overall, individuals were more related to individuals from the same swarm than from the alternate swarm. Therefore, the two hybrid swarm populations represented the full genetic diversity of their founding lines and were genetically distinct populations.

We calculated LD decay in the hybrid swarm, the hybrid swarm parents, and the DGRP using the *snpgdsLDMat()* function in *SNPRelate*. We randomly sampled 10,000 SNPs and then identified SNPs that were approximately 0.1, 0.5, 1, 5, 10, 50, 100, 500, and 1,000 kb away from the focal SNP (+/- 5%). When multiple SNPs were identified at the appropriate distance, a single SNP was randomly chosen. We performed this calculation for rare (minor allele frequency < 0.05), and common (minor allele frequency > 0.1) SNPs.

To calculate long-distance LD, we sampled SNPs in two ways. First, we sampled 10,000 random pairs of SNPs on opposite arms of chromosomes 2 and 3 (pairs on chromosome 2L and 2R, or 3L and 3R). Second, we sampled 30,000 random pairs of SNPs on independent chromosomes (e.g., one SNP on chromosome 2 and one SNP on chromosome 3.

### Inversion genotyping

The most likely genotypes at major cosmopolitan inversions in the parental strains and hybrid offspring were identified using diagnostic SNPs described in [133]. A binomial general linear model accounting for temperature, photoperiod, swarm, *Wolbachia*, and generation was used to test for the effect of each cosmopolitan inversion on diapause.

### Heritability estimation

We estimated narrow-sense heritability using *GCTA* [71] using the restricted maximum likelihood analysis (REML) method and the Fisher-scoring algorithm [72]. We generated a genetic relatedness matrix with the --make-grm flag in *GCTA*. We used the same covariates used for association mapping and flags --reml, --reml-alg 1, and --reml-bendV to calculate heritability for the original data as well as 1000 permutations of the sample ids. Confidence intervals were calculated as 1.96 * standard error.

### Mapping in *GENESIS*

We constructed GRMs using the *snpgdsGRM(method= “Eigenstrat”)* function in the *SNPRelate* package [132] using all SNPs that passed the mapping filter (see above) for both populations combined, as well as populations A and B separately. The appropriate GRM was then passed to *GENESIS* [134, 135] with the models:

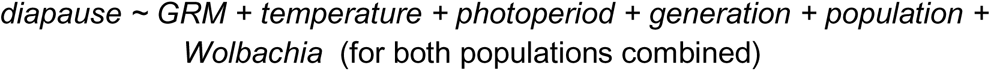

or

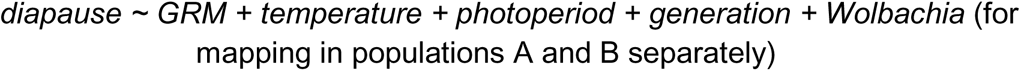

to calculate an effect at each SNP. A binomial model was used with diapause (at either stage 10 or stage 8) scored as 1 and non-diapause scored as 0. A separate GRM and mapping result was calculated for each of the 100 imputed datasets.

### Permuted GWAS

We performed 1000 permutations of the GWAS using both populations, and 100 permutations of each single-population GWAS. Phenotypes and environmental variables were permuted together so that the environmental effects would remain constant across permutations; our permutations shuffled the sample IDs *within* each hybrid swarm population to dissociate genotype and phenotype. For each permutation, one of the 100 imputed data sets was randomly chosen for genotypes. Identical permutations were used for the stage 8 and stage 10 data and for populations A, B, and both.

### LASSO SNPs

We used LASSO to identify a subset of informative SNPs out of the top 10,000 SNPs (ranked by *P*-value) calculated in each GWAS using the *cv.biglasso()* function the R package *biglasso* [136]. After filtering for a minor allele frequency of > 0.05 in the mapping population, the genotypes for the top 10,000 GWAS SNPs were added to a LASSO model that included temperature, photoperiod, generation, *Wolbachia*, and 32 principal components. We ran three models: 1) environment only, 2) environment + principal components and 3) environment + principal components + 10,000 genotypes. SNPs retained in model 3 are referred to as “LASSO SNPs”. We also used *snpgdsLDMat()* from *SNPRelate* to calculate LD among LASSO SNPs.

### Receiving operator characteristic (ROC) analysis

We used the R packages *ROCR* [75] to analyze the performance of environmental data, principal components, and LASSO SNPs in predicting individual phenotypes. The predictor variables chosen in each LASSO model were used to predict the stage 8 or stage 10 diapause phenotype with the *predict()* function. We then assessed these predictions using the *performance()* function with parameters “tpr”, “fpr”, and “auc”.

### SNP annotation

We annotated the predicted effects of all SNPs identified in the hybrid swarm using SNPEff [137] with reference genome BDGP5.75 using default parameters. We grouped these annotations into 6 categories: UTR, upstream/downstream (within 5 kb), intronic, intergenic, synonymous, and non-synonymous. For each imputation and permutation of the GWAS, we calculated the percentage of LASSO SNPs and top SNPs falling into each annotation category. We then compared the percentage of SNPs in each category in the permutations to the percentage of SNPs in that category in the observed data. To assign a *P*-value, we took the median percentage of the 100 imputations of the observed data and identified its quantile rank in the permutations. In this test, *P*-values of below 0.05 are indicative of a potential de-enrichment relative to permutations, whereas *P*-values above 0.95 are indicative of an enrichment.

### Clinal and seasonal polygenic score analysis

We examined spatiotemporal variation of diapause-associated SNPs using two previously generated datasets. Bergland *et al* (2014) sampled a cline from Florida to Maine and identified SNPs with repeatable seasonal changes in allele frequency in a single Pennsylvania orchard over the course of three years. Additionally, we used the data from Machado *et al* (2019), which also sampled a cline and calculated genome-wide seasonal changes in allele frequency across 20 populations sampled from North America and Europe. We re-calculated clinal *P*-values for all SNPs in the Machado *et al* (2019) dataset using a wider latitudinal spread than originally calculated. For each location near the East Coast (Homestead, FL; Athens, GA; Hahia, GA; Eutawville, SC; Charlottesville, VA; Media, PA; State College, PA; Lancaster, MA; Ithaca, NY; Bowdoin, ME), the average allele frequency across all sampling times was calculated for each SNP. These allele frequencies were corrected for the number of individuals sequenced and read depth as in [78], then regressed to latitude in a binomial general linear model. The effect sizes (beta) from the general linear model were extracted to determine the significance and direction of the cline.

We calculated polygenic scores [121,138–140] using the effect sizes and signs (the Score statistic from *GENESIS* for ranked SNPs or the coefficient from the LASSO model for LASSO SNPs) from the GWAS and the clinal and seasonal effect sizes described above. We then multiplied the GWAS effect by the clinal or seasonal effect for each SNP and summed these products across all tested SNPs. These tests were polarized such that SNPs with concordant signals (i.e., those where the pro-diapause allele is more common in the north or in the spring) will result in positive values for this score, whereas SNPs with discordant signals will result in negative values. We repeated this calculation for the permutations and compared the actual average polygenic scores to those calculated in the permutations.

We used a slightly different procedure to calculate polygenic scores for individual populations from [78]. We logit-transformed the fall and spring allele frequencies for each population and then subtracted the transformed fall frequency from the transformed spring frequency. This difference in logit allele frequencies was then multiplied by the GWAS effect size and summed as described above.

### Field experiment

To test whether our hybrid swarm populations carried seasonally selectable genetic variation for diapause, we placed hybrid swarm individuals (generation 30) into six outdoor field cages near Charlottesville, VA (37°57’33.5”N 78°28’18.5”W) in early June 2018. Three cages were initiated with each hybrid swarm population. Each cage was constructed around a dwarf peach tree and fed with approximately 3 kg of Red Delicious apples and 3 kg of bananas every week until November, when feeding was reduced to biweekly. Feeding was stopped in mid-December. Each cage was founded with approximately 3,000 flies that were then allowed to reproduce in overlapping generations. We sampled these populations, as well as the indoor hybrid swarm cages multiple times during the summer and fall (“G0” samples). For some collections, we dissected the field caught females to determine whether the ovaries appeared to be in diapause.

Because other Drosophilid species, including *D. simulans,* infiltrated our cages, it is possible that some of the G0 females dissected were in fact *D. simulans*; however, we never observed more than approximately 10% *simulans* males in any collection. It is also possible that some wild *D. melanogaster* infiltrated the cages. At Carter Mountain Orchard, which is located approximately 2 km from our field site, we observed that *D. simulans* made up 60-100% of all *simulans/melanogaster* individuals over the course of our field experiment. Therefore, if at most 10% of flies in the cages were *D. simulans*, we conservatively estimate that at most 5% of *D. melanogaster* in the cages were wild flies that unintentionally entered the cages.

To determine genetic changes in the propensity to diapause, we reared the sampled flies for two generations in the lab. For the first generation, isofemale lines were established in vials and offspring males were screened to determine *D. melanogaster* or *D. simulans* identity. All *D. melanogaster* G1s were combined and a subset were used to establish density-controlled bottles for G2s. The G2s were then assayed for diapause using our standard diapause assay (28 days, 9L:15D, either 10.5 or 12°C). We used a binomial generalized linear model to compare diapause incidence at each collection date to the initial timepoint (June 26^th^). Weather data was downloaded from wunderground.com; station ID: KVACHARL73 (Carter Mountain, VA).

### Population genetic analysis

We calculated allele frequencies from flies collected in Zambia during phase 3 of the Drosophila Genome Nexus [89, 90]. We used the full sequence FASTA files to generate allele frequencies at every SNP via custom scripts (https://github.com/alanbergland/DEST/). Using the GWAS results, we calculated the median allele frequency of the pro-diapause alleles in Zambia and compared these values to those generated using the permuted GWAS. To look for overlap with tracts of European admixture in Zambian flies, we downloaded the admixture tracts for this dataset (https://www.johnpool.net/genomes.html) and counted the number of times each SNP of interest overlapped with one of these tracts.

We used two North American fly samples to test for selective sweeps. First, we used publicly available data from the Drosophila Genetic Resource Panel (DGRP) [77]. Second, we used a set of 205 inbred lines collected from Pennsylvania and Maine (hereafter “Northern lines”). These lines were collected from Media, Pennsylvania in June and October of 2012, as well as from Bowdoinham, Maine in October 2012. The mapped reads for this dataset are available on the SRA (project number PRJNA383555). SNPs were called with *HaplotypeCaller* in *GATK* to produce gVCF files, combined with *CombineGenotypes*, and filtered for quality based on known SNPs from the DGRP. This VCF file was then used in parallel with the DGRP for population genetic analysis described below.

We calculated the integrated haplotype homozygosity score (iHS) [91] for every SNP in the DGRP or Northern lines using the R package *rehh* [141, 142]. Because iHS calculation in this package requires complete genotype information, we imputed missing genotypes in each dataset based on the most likely genotype given Hardy-Weinberg equilibrium as described for the hybrid swarm above. We calculated iHS by assigning the pro-diapause allele as the “derived” allele at each SNP based on the Score statistic from GENESIS. For each set of SNPs of interest, we calculated the median iHS and compared these values to those generated using the permuted GWAS.

### Statistical analysis and plotting

All analysis was performed using R versions 3.3 to 3.5 [143]. In addition to the aforementioned packages, the following packages were used for general analysis and plotting: *ggplot2* [144], *cowplot* [145], *data.table* [146], *foreach* [147], *doMC* [148], *ggbeeswarm* [149], *lubridate* [150], *maps* [64], and *viridis* [151].

### Data accessibility

All scripts and code used for data analysis and plotting are available at https://github.com/ericksonp/diapause-scripts-clean. All raw data and processed data used to generate figures are deposited in Data Dryad (available for review at: https://datadryad.org/review?doi=doi:10.5061/dryad.rd465d6; accession # doi:10.5061/dryad.rd465d6). All sequencing reads have been submitted to the Sequence Read Archive (BioProject # PRJNA522357).

## Acknowledgements

We thank Helen Stone, Liam Miller, Austin Edwards, and Sasha Bilal for laboratory and field assistance. We thank Erin Voss and Robert Porter for technical assistance. We thank AnhThu Nguyen from the University of Virginia Genomics Core for assistance with library preparation and Graham Wiley from the Oklahoma Medical Research Foundation for assistance with Illumina sequencing. We thank Mickey Pawlick and employees of Quality Machine Service for their assistance in designing and constructing the environmental chambers. We are grateful to David Glover and Richard Davis for assistance with the environmental chambers as well as research facility maintenance. We thank Andy Wyland at Morven Farms for assistance in establishing and maintaining our experimental orchard. We thank Sam Brunjes for helping establish the Raspberry Pi network used in the environmental chambers.

## Author contributions

PAE, PS, and AOB conceived of the study. PAE and AOB designed the study. PAE and DYS conducted the experiments. PAE and AOB analyzed the data using code contributed by CAW. ASB generated the Northern population VCF file. PAE and AOB wrote the manuscript. PAE, CAW, PS, and AOB edited the manuscript. PAE and AOB obtained funding for the experiments. All authors read and agreed to the manuscript prior to submission.

## Supporting Information

**S1 Fig.**
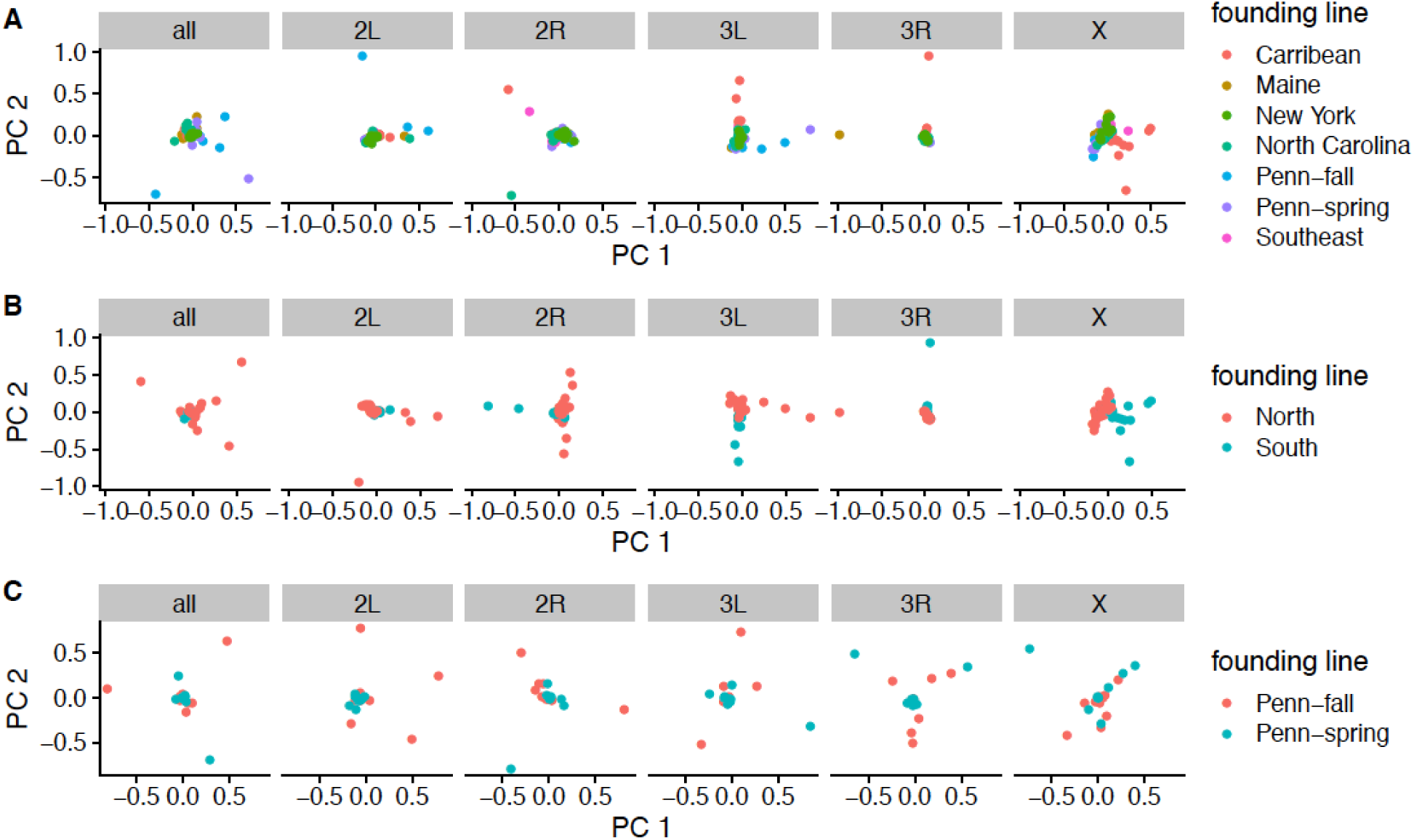
Principal component analysis of genetic diversity of parental lines. Principal components (PC) were calculated for (A) all parental lines, (B) northern and southern lines (excluding the DGRP from North Carolina), and (C) lines with known spring and fall collection dates. PCs were calculated genome-wide (all) or for each chromosome arm separately.

**S2 Fig.**
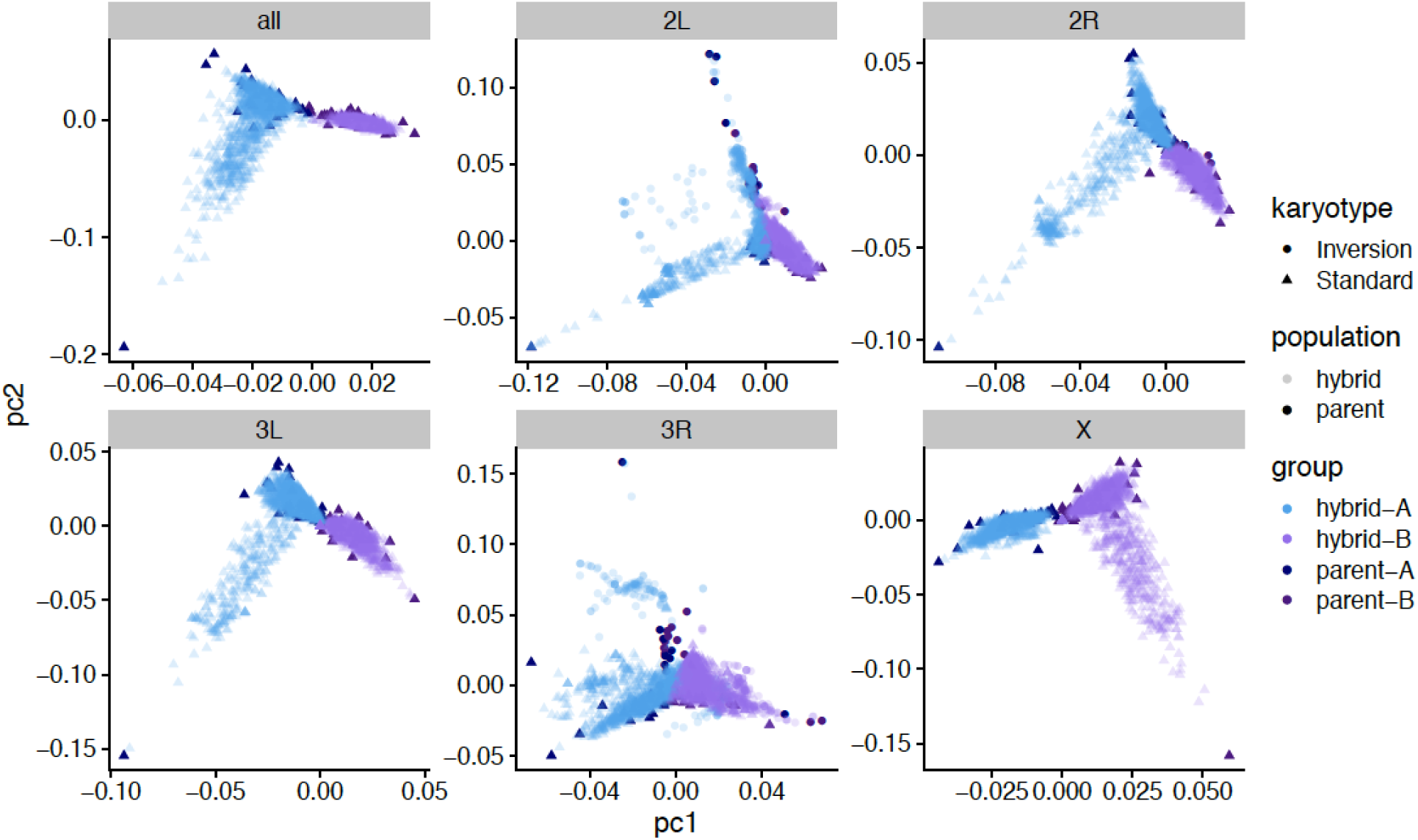
Principal components analysis of hybrid swarm populations, including parental lines. PC1 and PC2 are plotted for all chromosomes combined (all), as well as each individual chromosome arm, and are color coded by population. Dark shapes indicate parental lines, while transparent shapes indicate individual hybrids. Circles indicate an individual that is either heterozygous or homozygous for one of the cosmopolitan inversions on that chromosome.

**S3 Fig.**
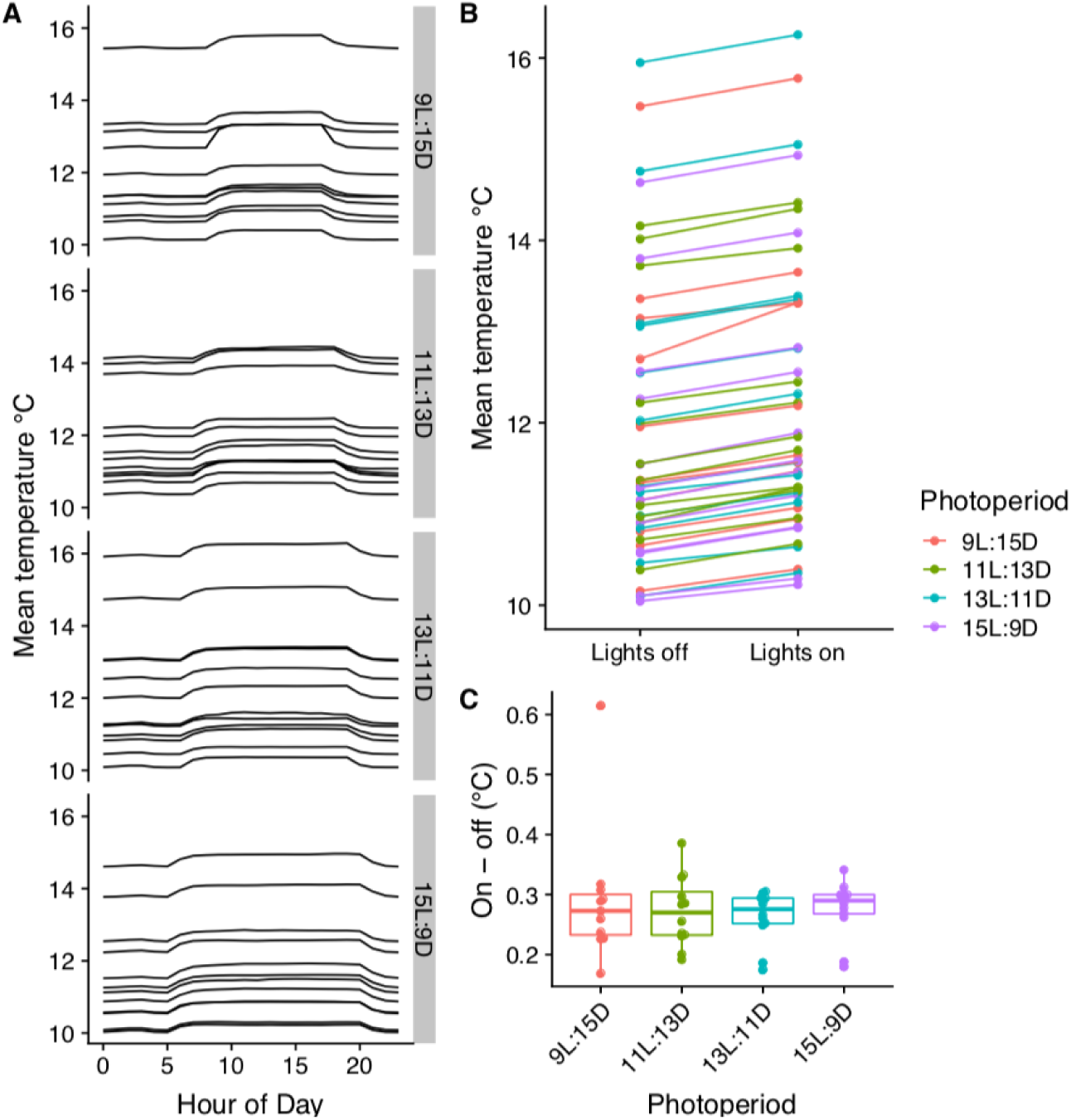
Environmental control chambers maintain constant temperatures with little temperature change from lights. (A) Average temperature of each box across a 24 hour day, separated by photoperiod into vertical facets. Each line represents the mean temperature of a single box recorded every 60 s for ∼6 weeks. (B). Average temperature of each box when lights are off and lights are on, color coded by photoperiod. (C). Difference in average temperature when lights are on and lights are off. Each point represents one box.

**S4 Fig.**
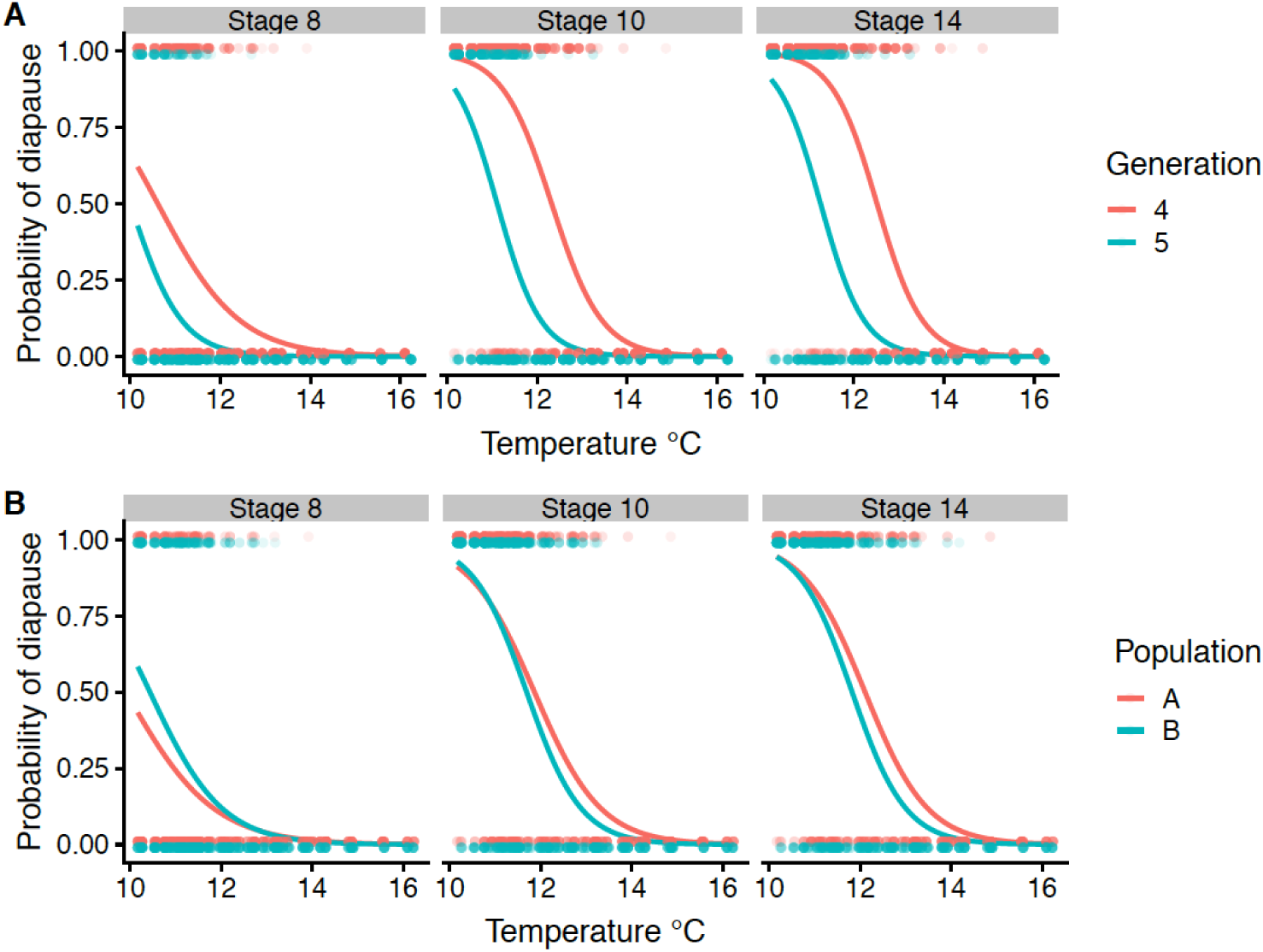
Diapause phenotypes differ between generations and cages. Grey points indicate individual phenotypes (1= diapause, 0 = non-diapause). Lines represent binomial models for each group. (A) F4s had uniformly higher diapause incidence than F5s, regardless of phenotype assessed (general linear model, *P* < 2 x 10^-16^ for all). (B) The two hybrid swarms were also significantly different for all three diapause phenotypes. For stage 8, population B had generally higher diapause incidence, while cage A had higher incidence for stages 10 and 14 (general linear model, *P* = 0.0006, *P* = 0.01, *P* = 1.75 x 10^-5^, respectively).

**S5 Fig.**
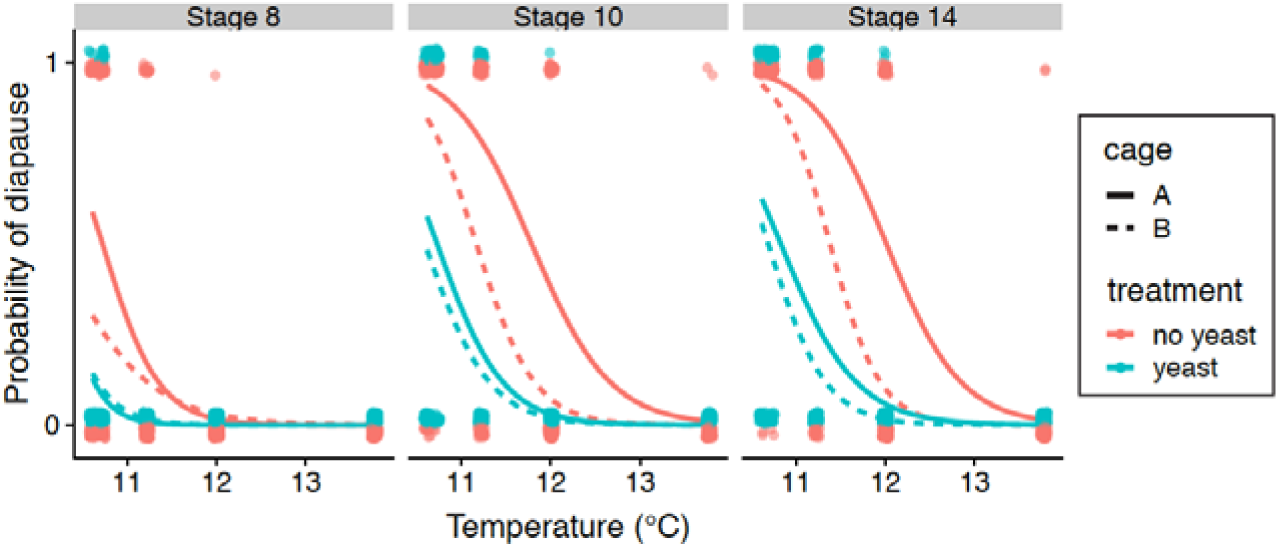
Live yeast supplementation decreases diapause incidence across temperatures. Advanced generation hybrid swarm individuals were exposed to diapause inducing conditions (9L:15D, 10-14°C) with or without a sprinkling of live baker’s yeast on the surface of the food. Points represent individuals with jitter added for visual clarity (1= diapause, 0 = nondiapuse); lines represent binomial models for each population and treatment. The presence of yeast decreased diapause at all thresholds scored (binomial general linear model, *P* < 2 x 10^-16^ for all stages). There was a significant effect of population for all three phenotypes, with cage A showing higher diapause incidence than cage B, regardless of temperature or yeast treatment (*P* = 0.03, *P* = 3.3 x 10^-6^, *P* = 7.1 x 10^-7^).

**S6 Fig.**
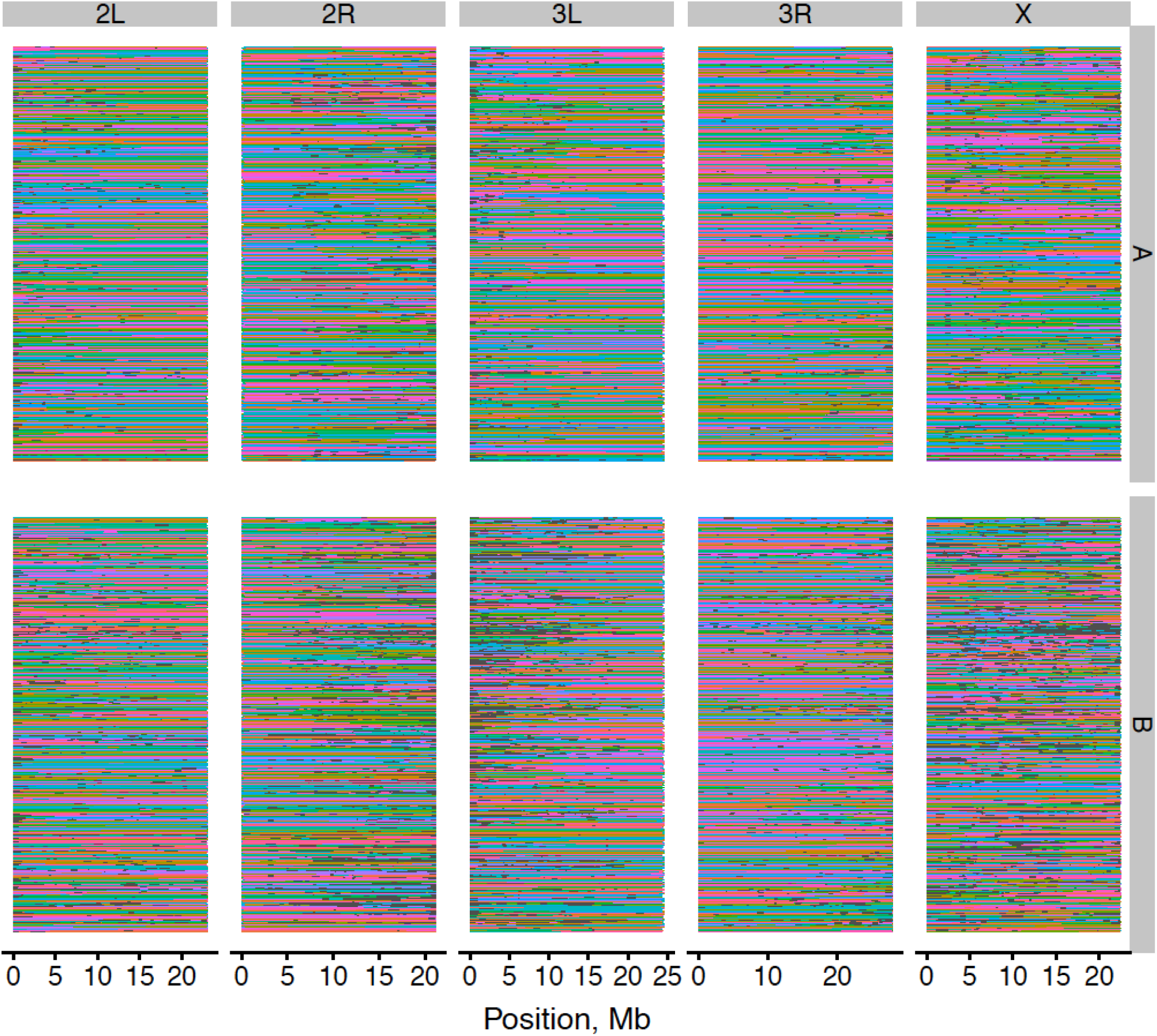
Genome reconstructions for individual hybrids from populations A (top) and B (bottom). Each horizontal line represents one haploid chromosome (n=2823 diploid individuals total). Data are separated by chromosome arm (horizontal) and population (vertical). Grey indicates regions that were masked due to short inferred parental haplotypes (roughly 1.2% of sequences).

**S7 Fig.**
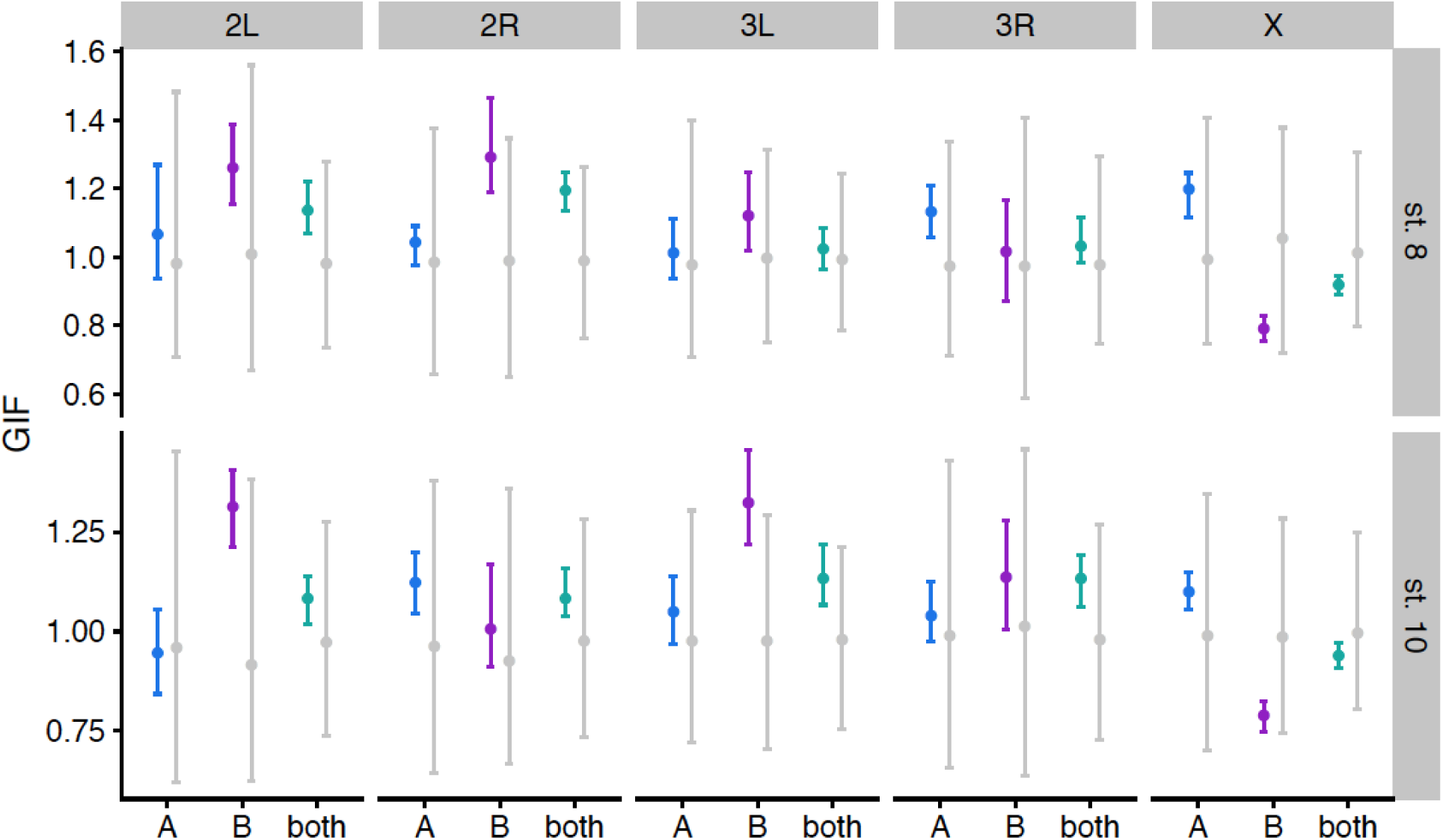
λ_GC_ is slightly inflated relative to permutations for autosomes, but not X. The genomic inflation factor (GIF) was calculated for each chromosome separately in al three mapping populations. Colors illustrate 100 imputations of the observed data; grey indicates permutations (100 permutations for A and B, 1000 permutations for the combined data). Points indicate the median, bars illustrate the 2.5%-97.5% quantiles.

**S8 Fig.**
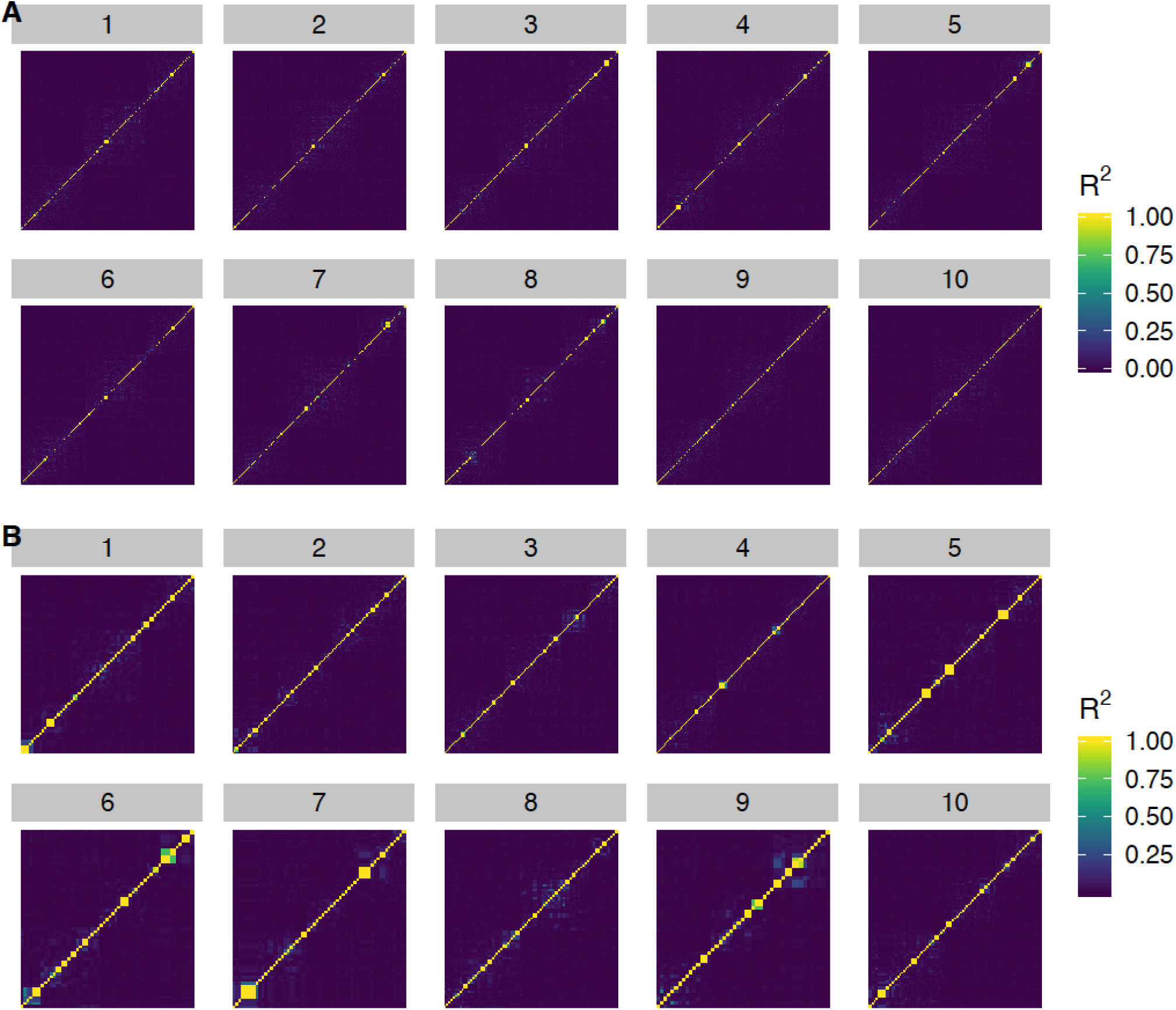
LASSO SNPs are generally unlinked. LD heatmaps showing R^2^ for LASSO SNPs from 10 imputations of the original ordering of the data (A) and 10 random permutations (B). LASSO SNPs for stage 10 diapause from the combined A+B mapping population were used to generate this figure.

**S9 Fig.**
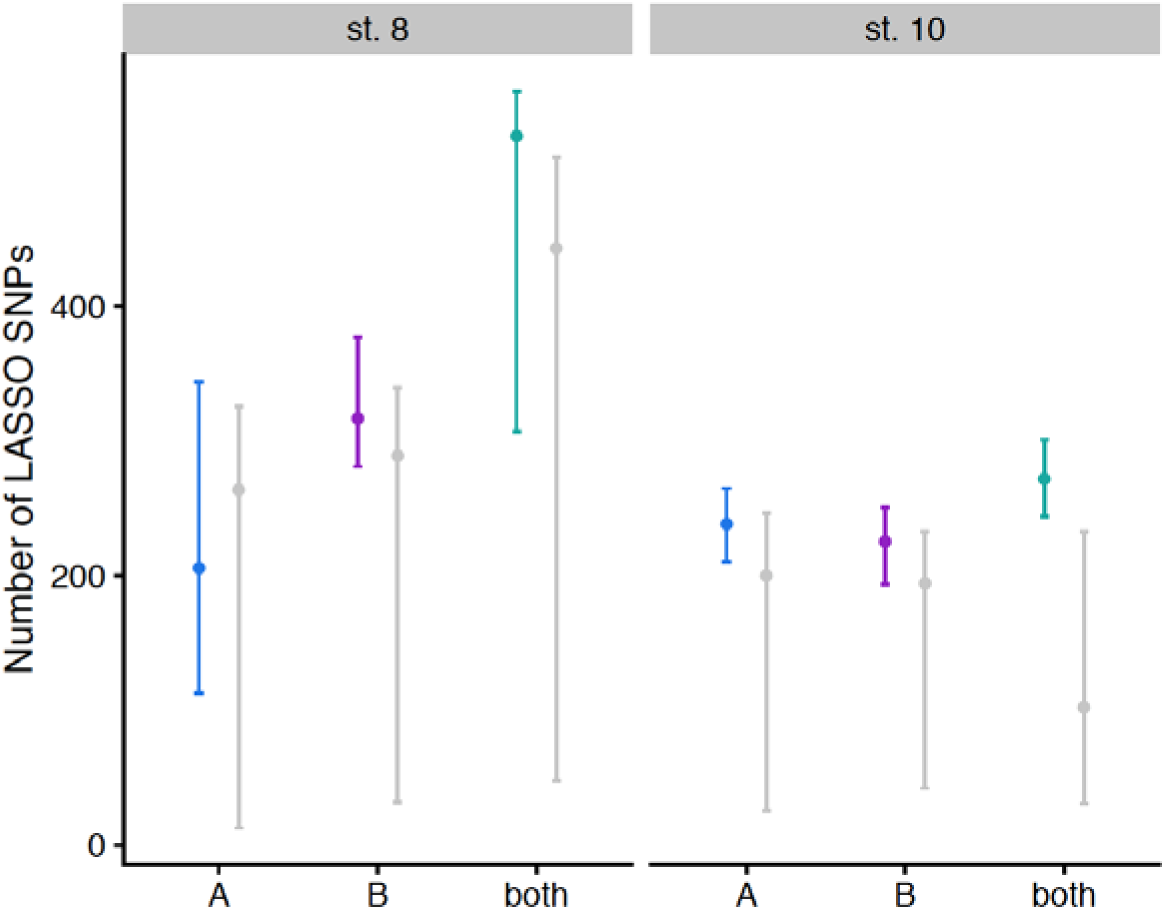
Number of SNPs chosen by LASSO is higher for actual data relative to permutations. The number of LASSO SNPs was counted for each imputation or permutation of each phenotype in each mapping population. Points represent the median; bars extend to the 2.5% and 97.5% quantiles. Colors represent 100 imputations of the observed data; grey bars represent permutations (100 permutations for A and B, 1000 permutations for both).

**S10 Fig.**
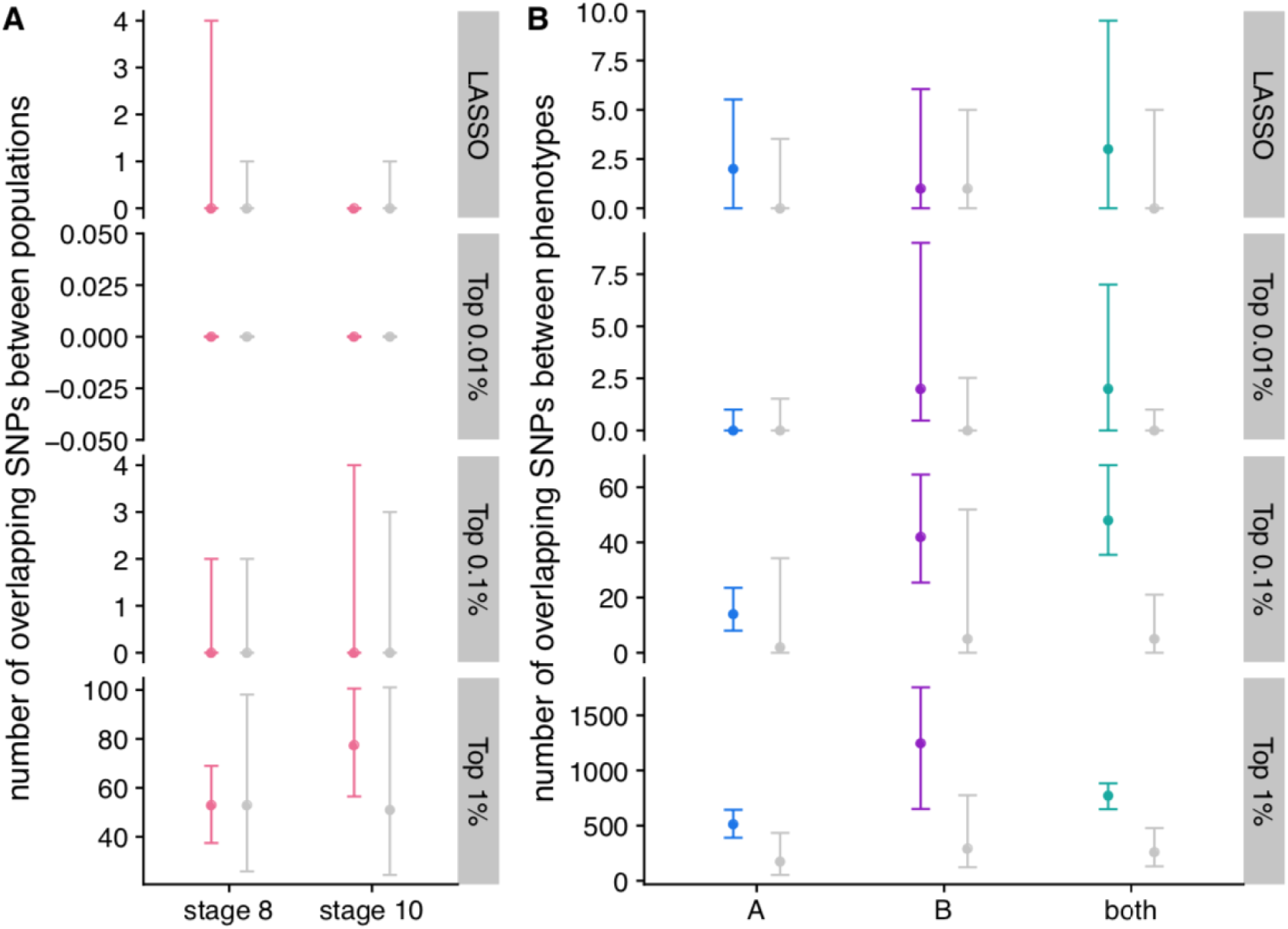
Number of diapause-associated SNPs shared between mapping populations and phenotypes. (A). For each imputation and permutation, the number of SNPs shared between populations A and B was counted for various sets of GWAS SNPs. Pink indicates 100 imputations of the actual data, grey indicates 100 permutations. Points represent the median and error bars extend to the 2.5% and 97.5% quantiles. (B) For each imputation and permutation, the number of SNPs shared between stage 8 and stage 10 diapause was counted. Grey points represent 100 permutations of populations A and B; 1000 permutations of both populations combined. Colored points represent 100 imputations of the actual data.

**S11 Fig.**
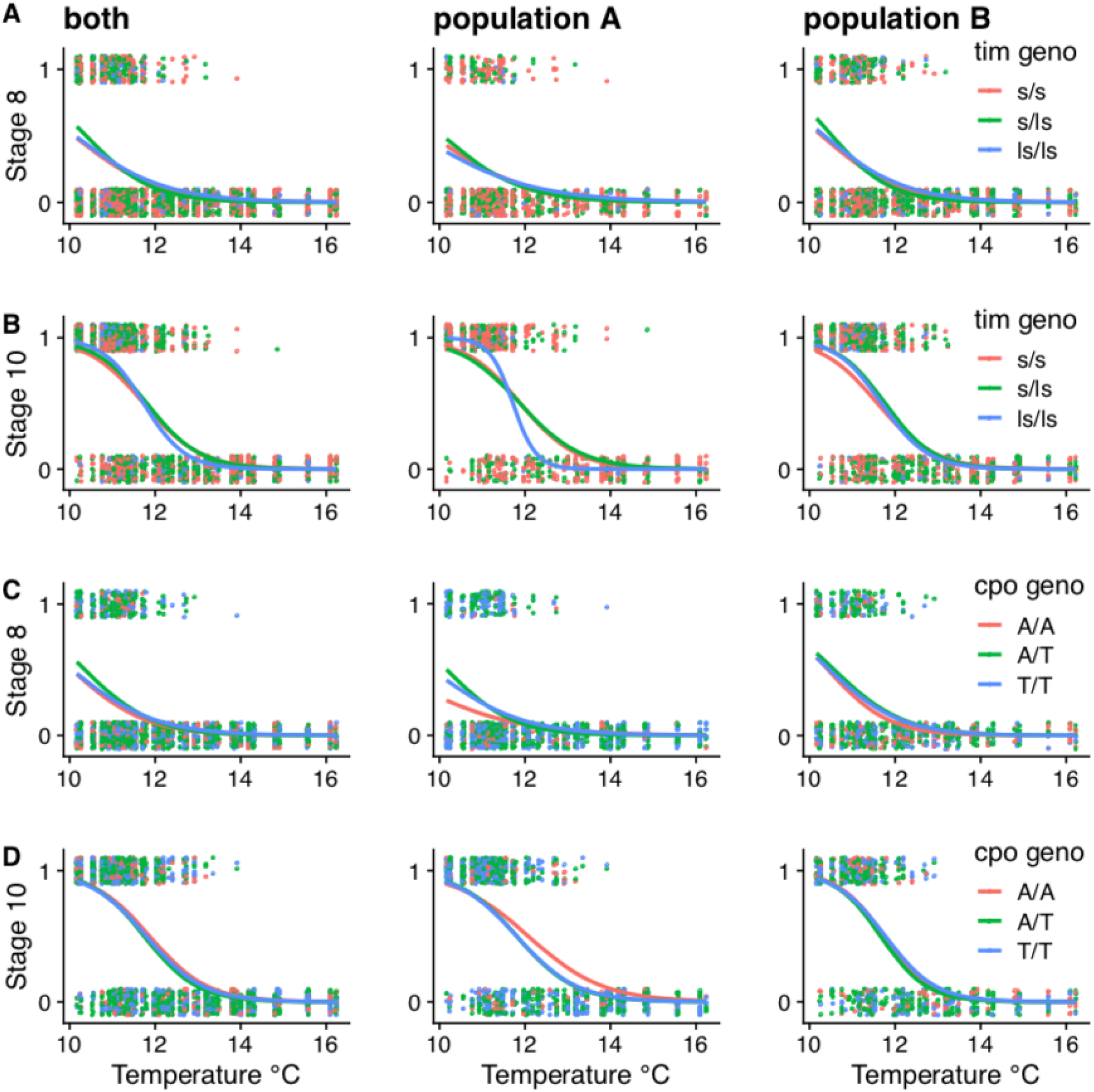
Previously identified diapause-associated variants have no effect in this study. (A-B). A 1 bp indel that creates an alternate translation start site in *timeless* (*tim*) does not affect diapause in the full dataset or either individual population after correcting for temperature for diapause at stage 8 (A) or stage 10 (B). (C-D) An intronic SNP in *couch potato (cpo)* also has no effect in the full dataset or either individual population after correcting for temperature for diapause at stage 8 (C) or stage 10 (D). General linear model *P*-values for genotype are greater than 0.05 for all models, except for *tim* in population B, stage 8 (*P* = 0.0189).

**S12 Fig.**
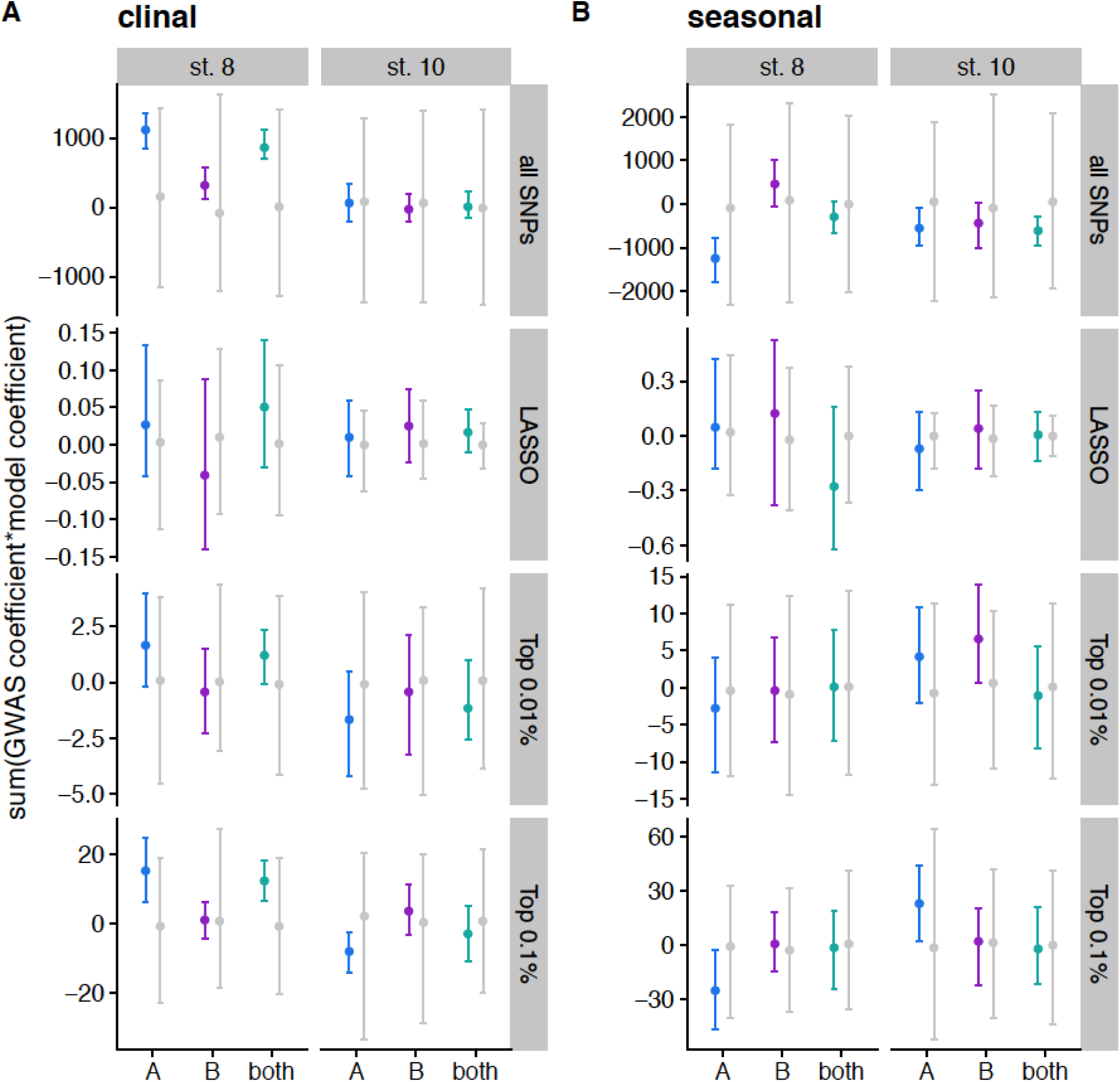
Polygenic score test for data from Machado *et al*, 2019. A) Polygenic scores calculated by multiplying clinal effect size and GWAS effect sizes for each SNP and summing across all SNPs, LASSO SNPs, the top 0.01% of the GWAS, and the top 0.1% of the GWAS. Effect sizes are polarized such that positive numbers indicated pro- diapause alleles are more common in the north. Data are shown with a point for the mean and error bars extending to the 2.5% and 97.5% quantiles. Colored points indicate actual data for 100 imputations of each mapping population; grey points indicate the distribution for permutations. B) Polygenic scores calculated for seasonal data by multiplying seasonal betas and GWAS effect sizes, polarized so that pro-diapause and spring are positive.

**S13 Fig.**
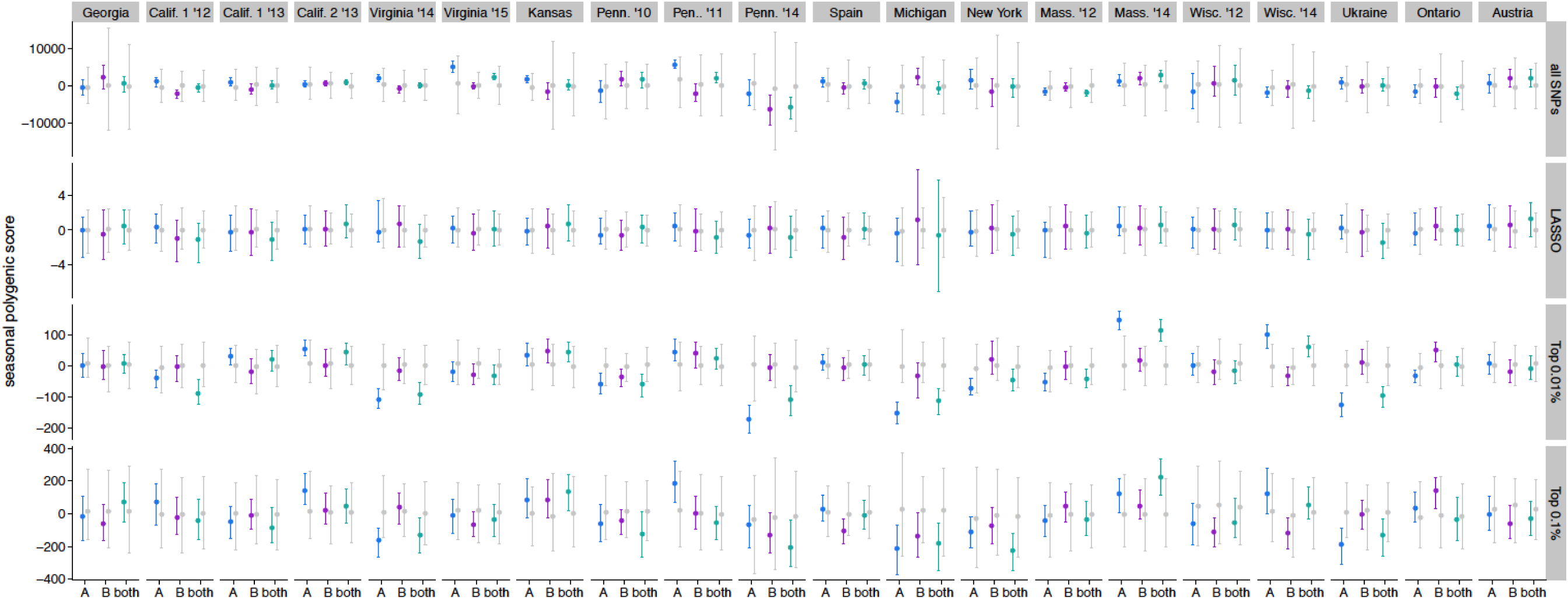
Individual population polygenic score test for stage 8 diapause in populations collected by Machado *et al* (2019). The GWAS or LASSO effect size was multiplied by the logit-transformed change in allele frequency from spring to fall in 20 populations. These products were then summed across all SNPs of interest for each mapping population. Populations are ordered by increasing latitude and year. Points represent median, error bars represent 2.5% and 97.5% quantiles. Grey points/bars are permutations; colors represent 100 imputations of the observed data.

**S14 Fig.**
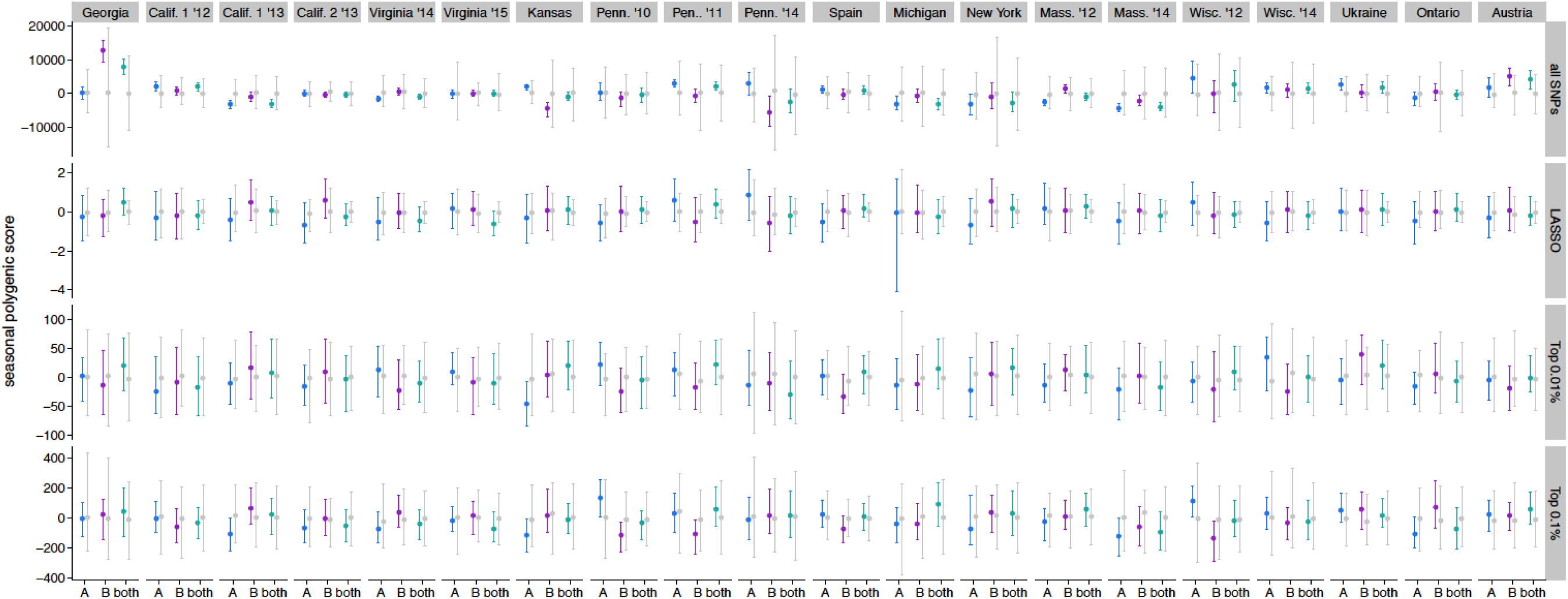
Individual population polygenic score test for stage 10 diapause in populations collected by Machado *et al* (2019). The GWAS or LASSO effect size was multiplied by the logit-transformed change in allele frequency from spring to fall in 20 populations. These products were then summed across all SNPs of interest for each mapping population. Populations are ordered by increasing latitude and year. Points represent median, error bars represent 2.5% and 97.5% quantiles. Grey points/bars are permutations; colors represent 100 imputations of the observed data.

**S15 Fig.**
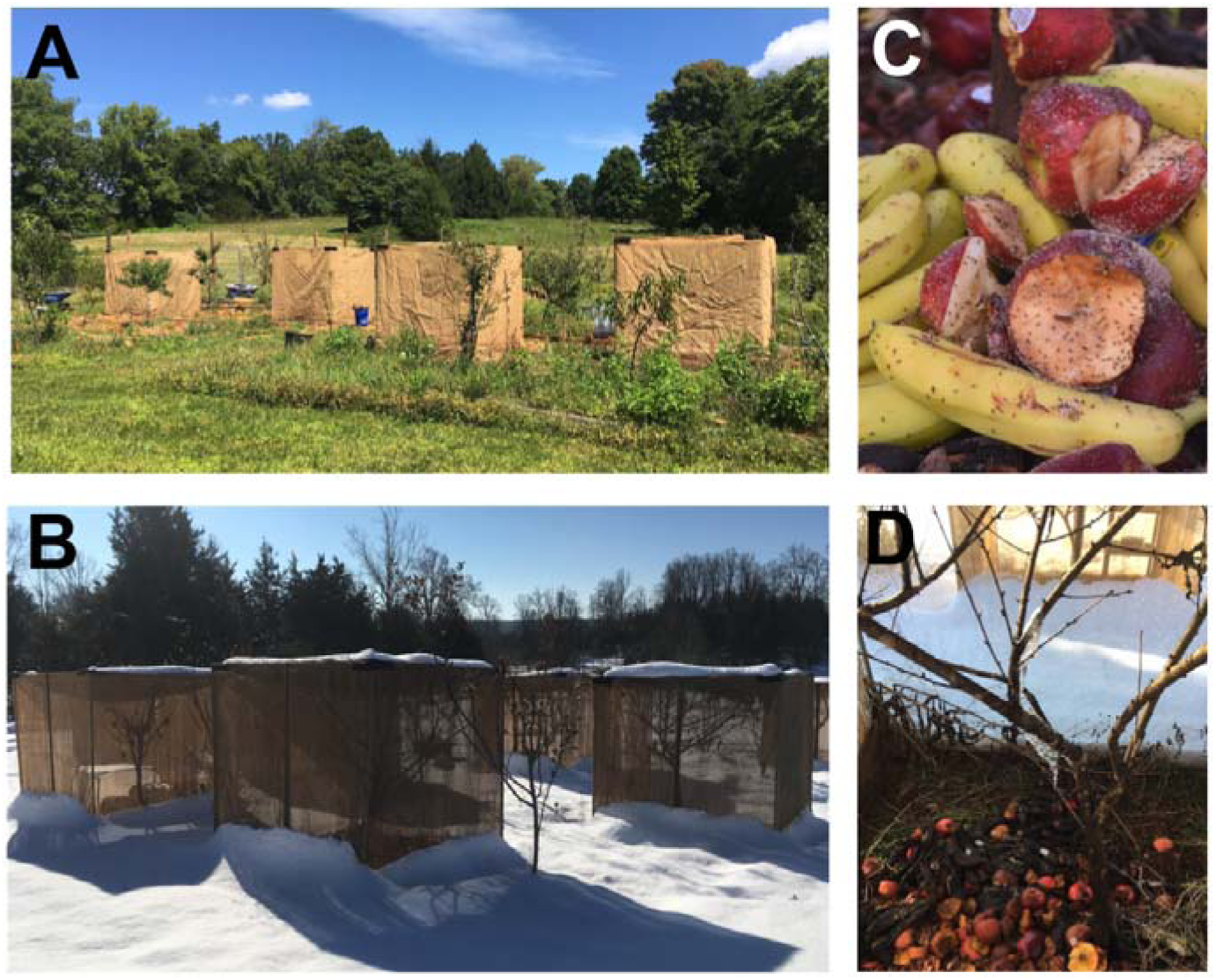
Field cages used to study diapause evolution under natural conditions. Summer (A) and winter (B) views of the experimental orchard with peach trees enclosed in mesh cages. (C) Flies feeding on yeasted fruit. (D) Winter view of compost pile in more advanced stage of decomposition. Photos in B and D were taken on December 10^th^, 2018, the final collection point in Fig 6. Surviving *D. melanogaster* were recovered from the cages on this day, despite heavy snow and several days of sub-freezing temperatures.

**S16 Fig.**
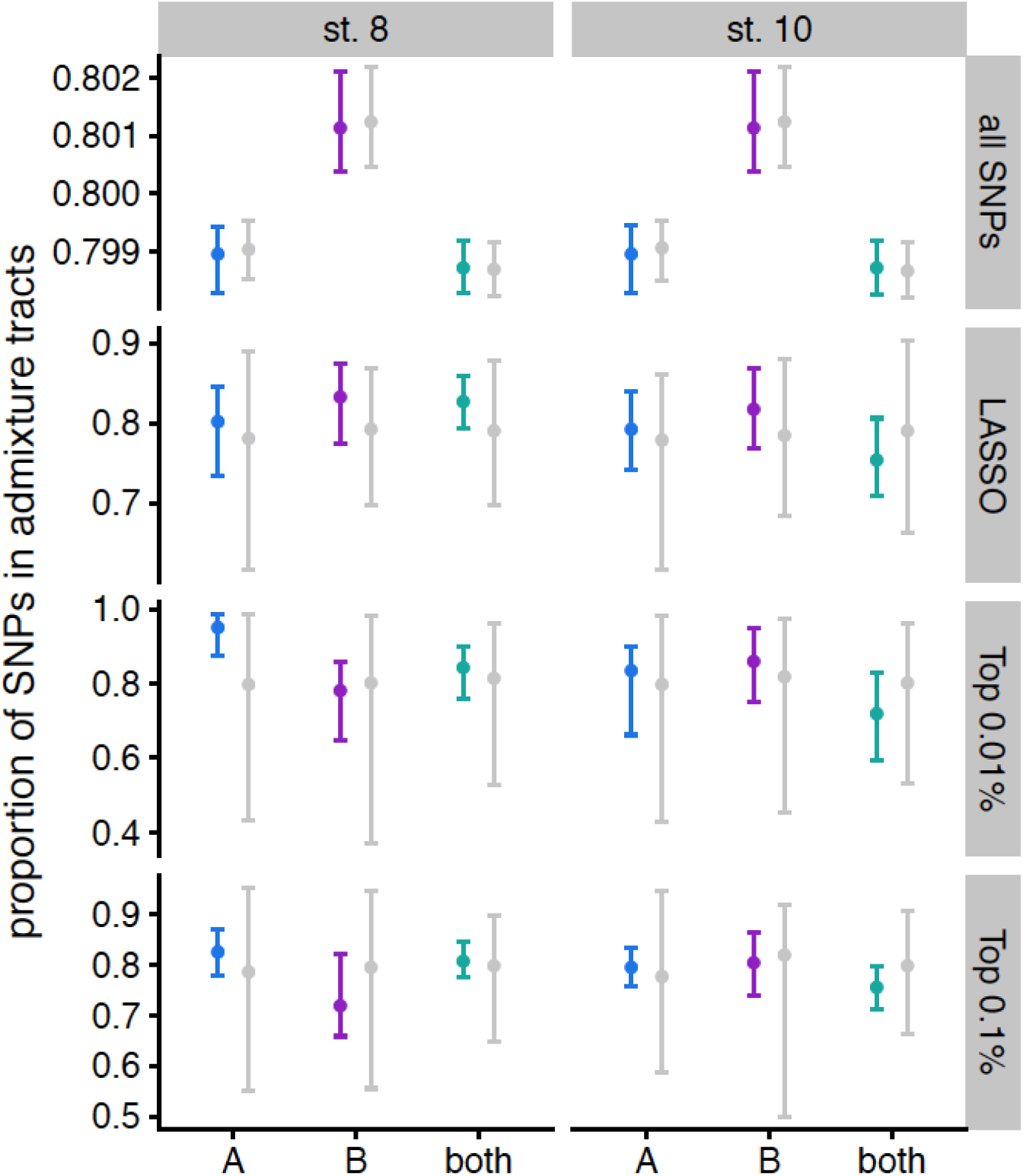
GWAS SNPs are not enriched in tracts of admixture between European and Zambian flies. Each set of SNPs was intersected with the admixture tracts of European/Zambian admixture, and the proportion of SNPs found within at least one admixture tract was calculated for each mapping population. Points represent median, error bars represent 2.5% and 97.5% quantiles. Grey points/bars are permutations; colors represent 100 imputations of the observed data.

**S17 Fig.**
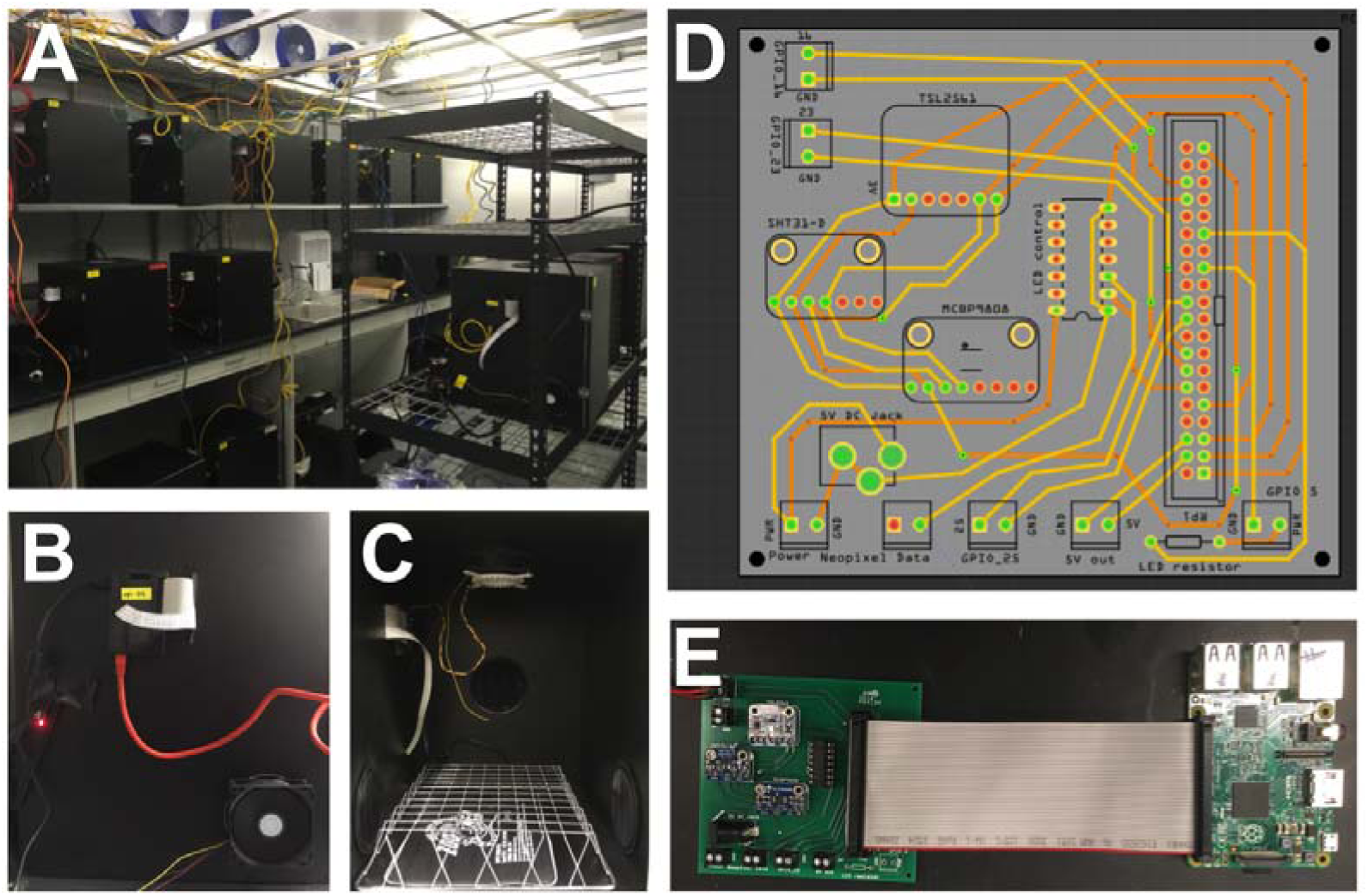
Photoperiod chambers. A) Array of chambers in cold room. B) Side view illustrating fan and externally mounted Raspberry Pi with ethernet connection. C) Interior view illustrating heating element (bottom), light-proof vents (sides), LED lights (top) and circuit board (top left). D) Fritzing layout for custom printed circuit board (PCB). File available upon request. E) Circuit board connected to Raspberry Pi computer via 40 pin ribbon cable.

**S18 Fig.**
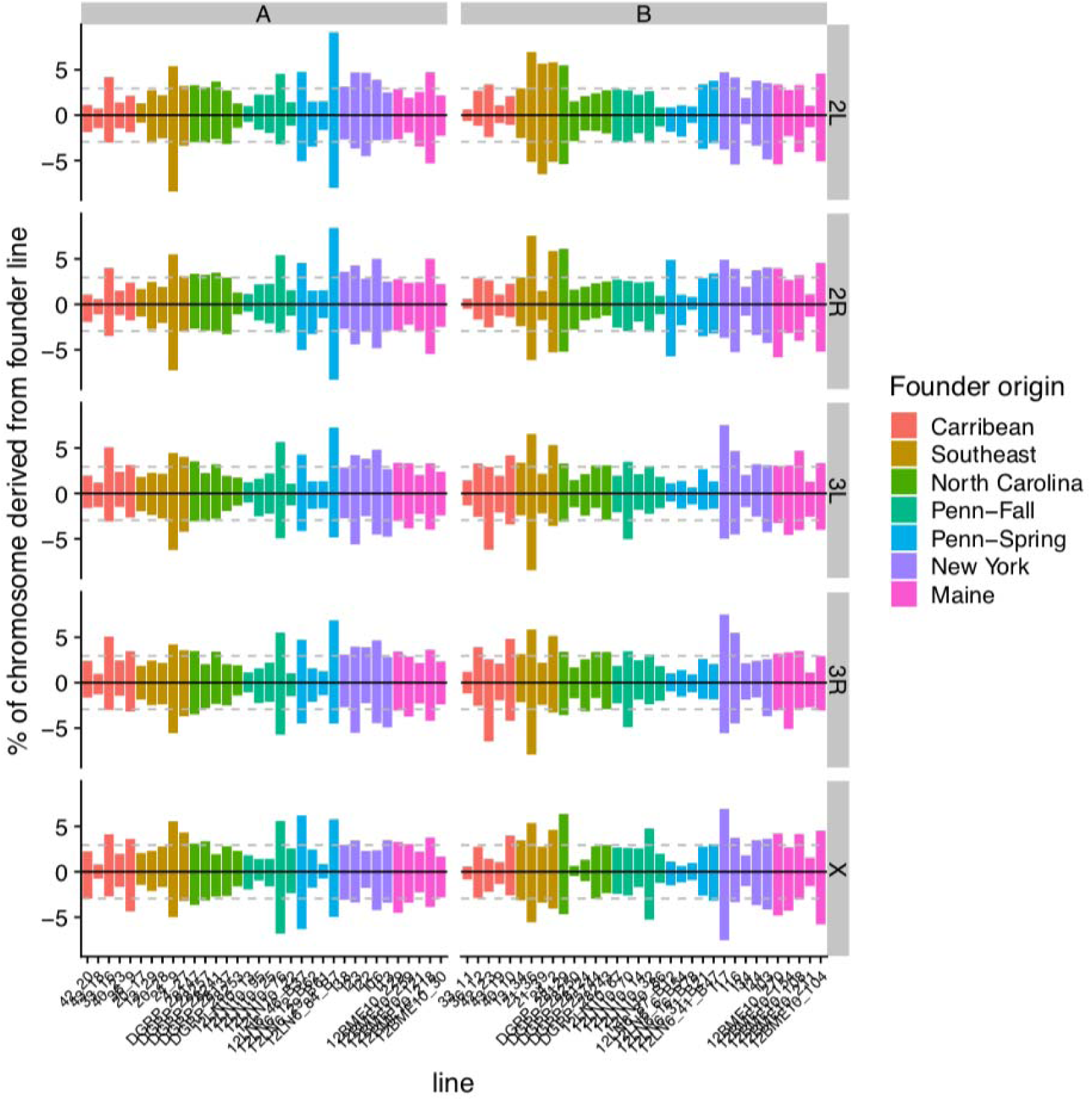
All founding lines are represented in F4 and F5 hybrid swarms. The percent contribution of each founding line in each chromosome arm was calculated for swarm A (left) and B (right). F4s are shown with positive values, and F5s are mirrored with negative values below. Color coding corresponds to geographical origin of the lines. Dashed grey lines indicate the expected contribution of each line (1/34 = 2.9%) under perfectly even admixture.

**S19 Fig.**
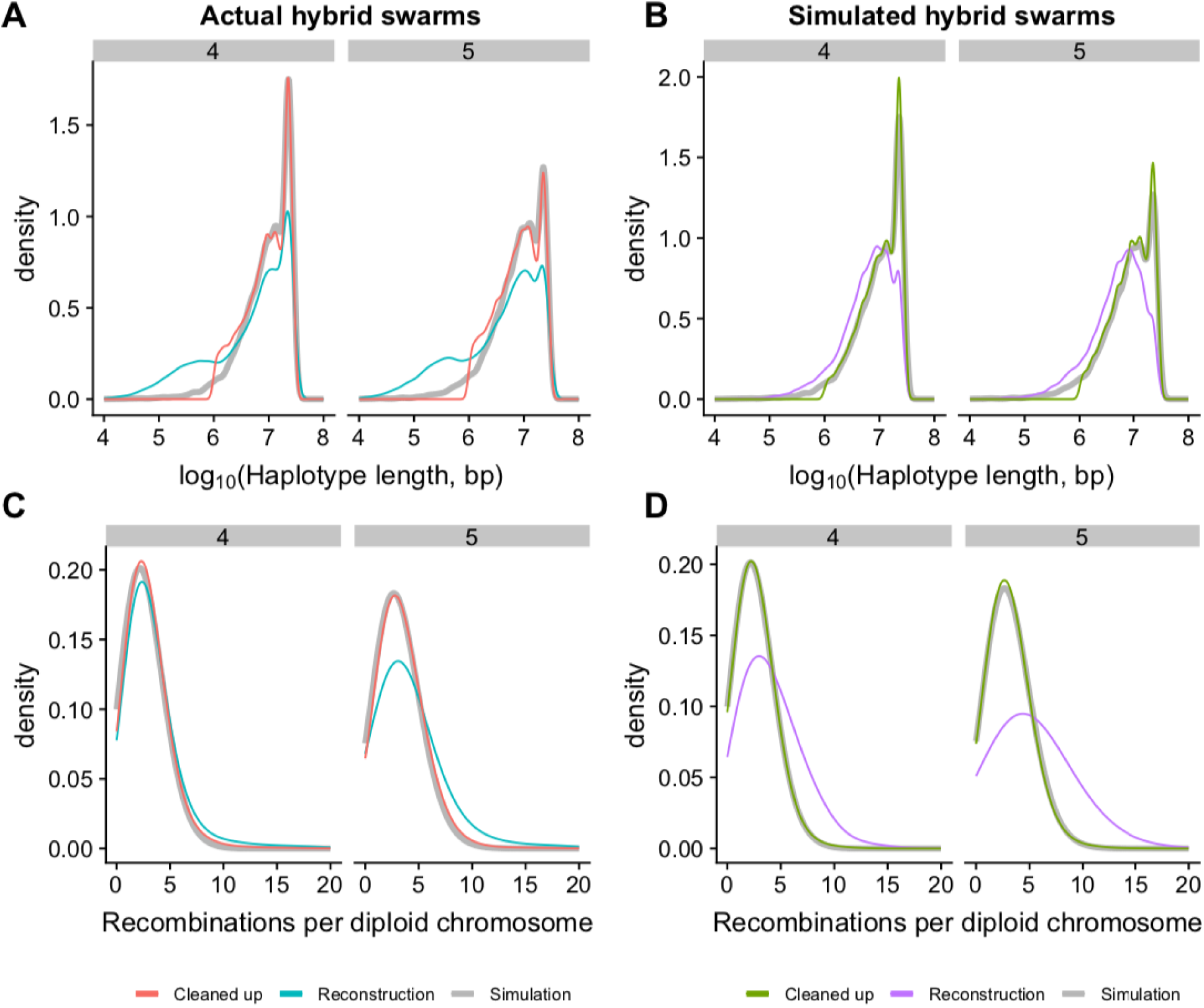
Haplotype size and recombination distributions for reconstructed F4 and F5 genomes. (A-B) The initial genome reconstruction of sequencing data (blue, A) shows an excess of short (∼10,000-1,000,000 bp) haplotypes relative to simulated F4 and F5 populations (grey bold line). This excess also occurs in reconstructions of simulated hybrid swarm reads (B, purple). Cleaning up the reconstruction data by combining adjacent short (< 1 Mb) haplotypes into unknown haplotypes and dropping singleton short haplotypes results in a distribution of haplotype sizes that more closely match the simulations for both empirical data (red, A) and simulated data (green, B). (C-D) Raw reconstructions have an excess of recombination events (blue and purple) relative to simulated data (grey). The cleanup procedure results in recombination numbers on par with the simulated data (red and green).

**S20 Fig.**
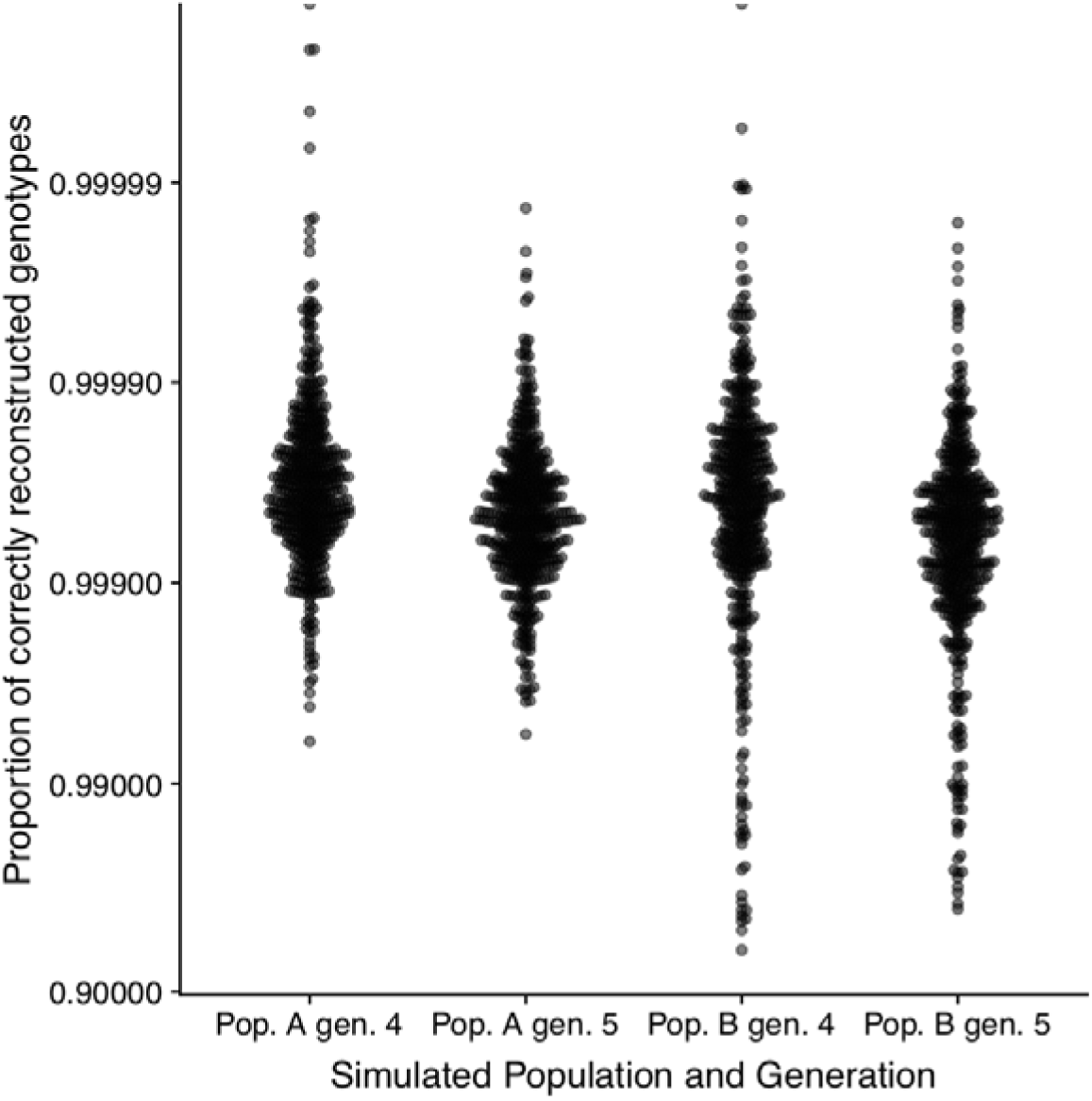
Accuracy of simulated genome reconstructions. Hybrid F4 and F5 individuals were simulated from the founding lines for populations A and B using a custom script, and 0.5X coverage sequencing reads were generated with *wgsim*. These simulated reads were passed through the genome reconstruction pipeline, and the reconstructed genotypes were compared to the original simulated individual. Accuracy was determined as the proportion of all sites with an exact match between the reconstructed genotype and actual genotype. The vast majority of individuals have an accuracy of >99%, though accuracy is higher in population A than population B. Note logarithmic scale of y-axis.

**S21 Fig.**
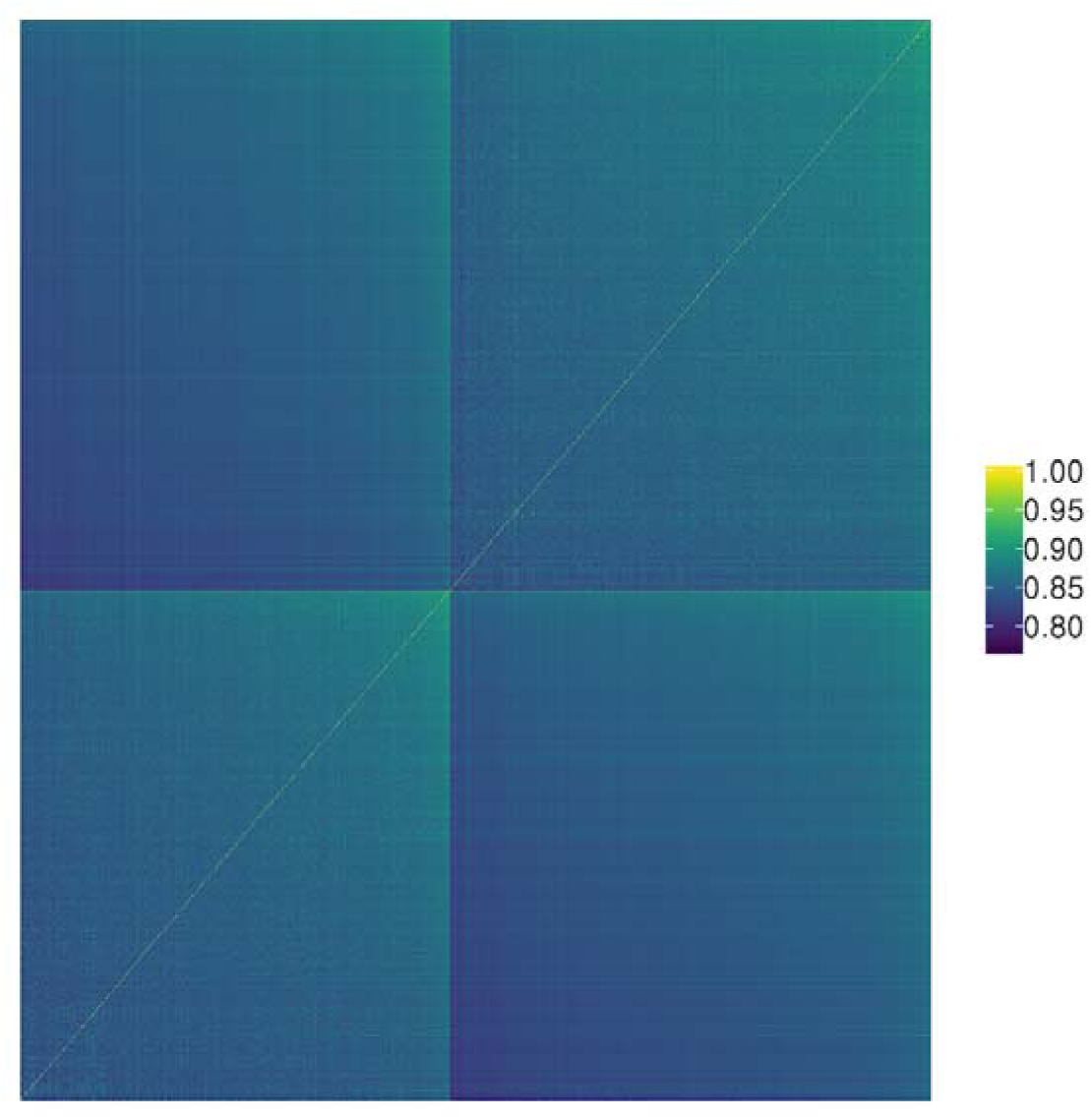
Identity-by-state genetic relatedness matrix for all hybrid swarm individuals. Individuals are ordered by population (A on left, B on right) and sorted by identity by state. IBS was calculated using an LD-pruned set of ∼63,000 SNPs with allele frequencies > 0.05. Individuals are generally more closely related to other individuals in the same population.

Table. S1 SRA and collection information for previously sequenced parental lines. ***(Excel file)***

**S2 Table.** Variance decomposition for variables influencing diapause.

|  | Stage 8 |  |  |  | Stage 10 |  |  |  |
| --- | --- | --- | --- | --- | --- | --- | --- | --- |
|  | Sum Sq | F | P | PVE | Sum Sq | F | P | PVE |
| temperature | 504.4 | 651.1 | 2.37E-129 | **17.67** | 1558.9 | 1676.2 | 8.55E-288 | **33.84** |
| photoperiod | 1.8 | 2.3 | 0.127 | **0.06** | 16.0 | 17.2 | 3.50E-05 | **0.35** |
| generation | 141.8 | 183.1 | 1.86E-40 | **4.97** | 416.0 | 447.3 | 3.04E-92 | **9.03** |
| population | 17.8 | 23.0 | 1.75E-06 | **0.62** | 0.8 | 0.9 | 0.34 | **0.02** |
| Wolbachia | 8.2 | 10.5 | 0.0011 | **0.29** | 0.4 | 0.5 | 0.49 | **0.01** |
| temperature*photoperiod | 3.6 | 4.7 | 0.03 | **0.13** | 0.7 | 0.7 | 0.39 | **0.01** |
| residuals | 2177.8 | - | - | **76.27** | 2613.3 | - | - | **56.74** |

**S3 Table.**
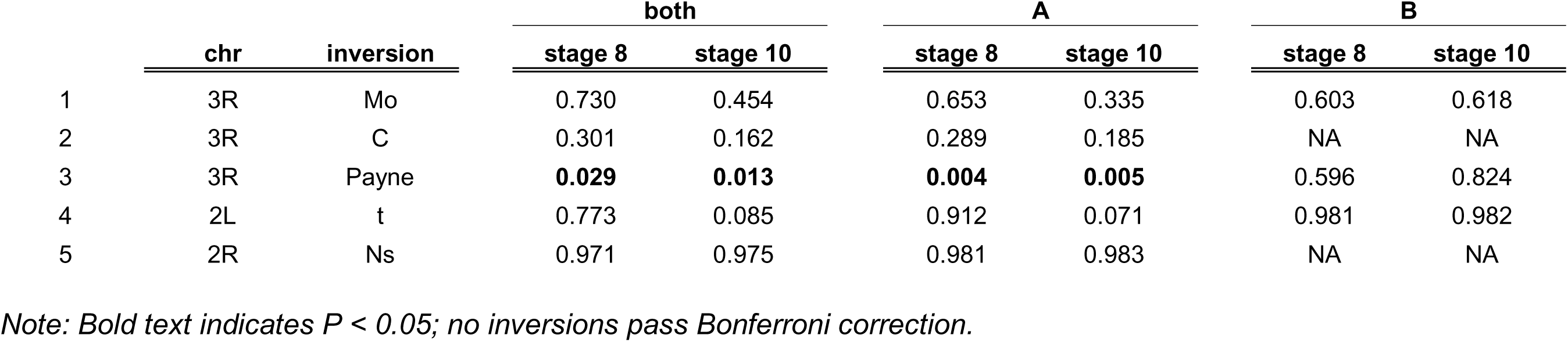
*P*-values from general linear model of the effects of cosmopolitan inversions on diapause in each population

|  | chr | inversion | both |  | A |  | B |  |
| --- | --- | --- | --- | --- | --- | --- | --- | --- |
|  |  |  | stage 8 | stage 10 | stage 8 | stage 10 | stage 8 | stage 10 |
| 1 | 3R | Mo | 0.730 | 0.454 | 0.653 | 0.335 | 0.603 | 0.618 |
| 2 | 3R | C | 0.301 | 0.162 | 0.289 | 0.185 | NA | NA |
| 3 | 3R | Payne | **0.029** | **0.013** | **0.004** | **0.005** | 0.596 | 0.824 |
| 4 | 2L | t | 0.773 | 0.085 | 0.912 | 0.071 | 0.981 | 0.982 |
| 5 | 2R | Ns | 0.971 | 0.975 | 0.981 | 0.983 | NA | NA |

*Note: Bold text indicates $P < 0.05$ ; no inversions pass Bonferroni correction.*

**S4 Table:**
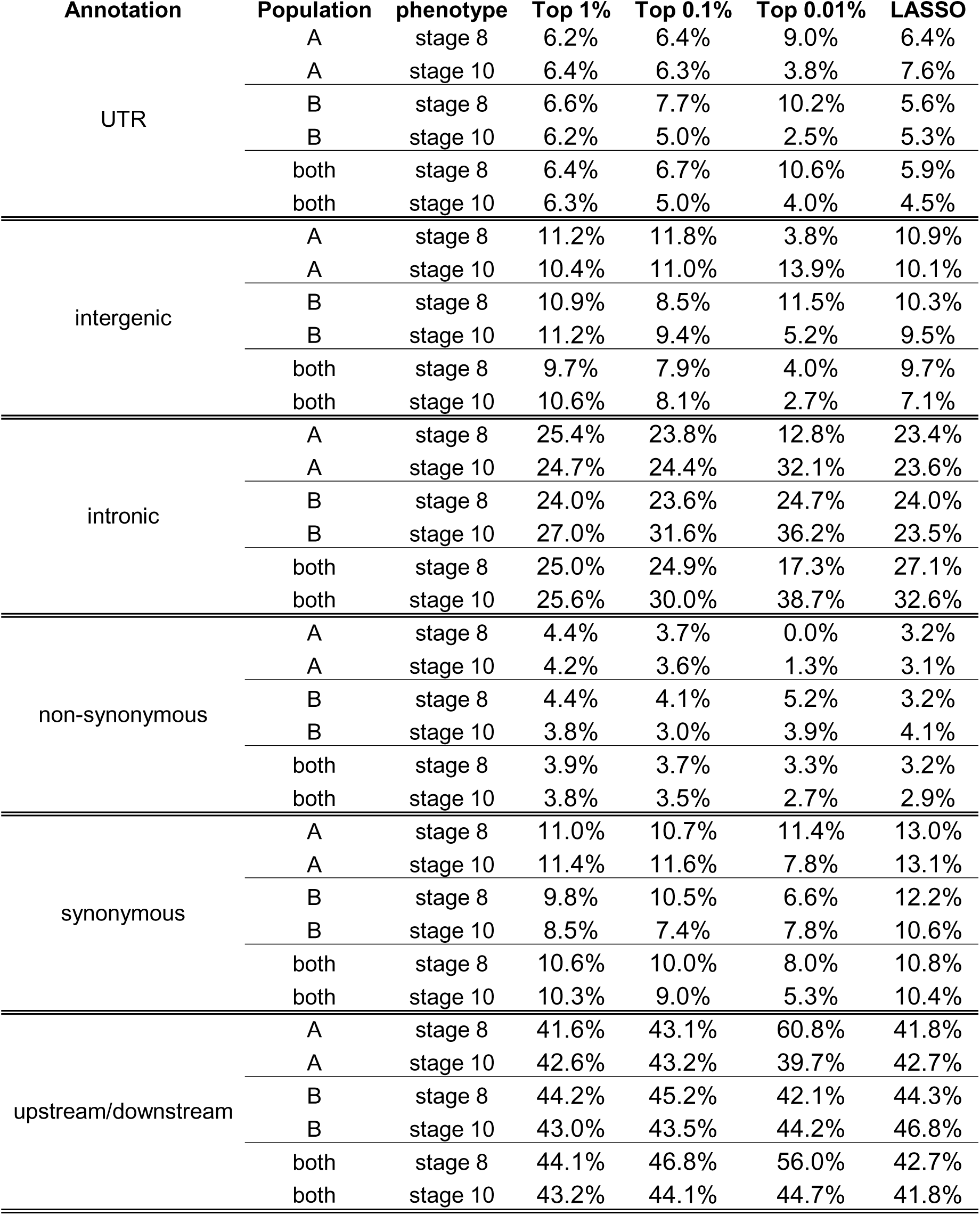
**Functional annotations in diapause-associated SNPs.** For each class of variant, the percentage of diapause-associated SNPs assigned to that variant type was quantified. The percentage reported is the median of 100 imputations.

| Annotation | Population | phenotype | Top 1% | Top 0.1% | Top 0.01% | LASSO |
| --- | --- | --- | --- | --- | --- | --- |
| UTR | A | stage 8 | 6.2% | 6.4% | 9.0% | 6.4% |
|  | A | stage 10 | 6.4% | 6.3% | 3.8% | 7.6% |
|  | B | stage 8 | 6.6% | 7.7% | 10.2% | 5.6% |
|  | B | stage 10 | 6.2% | 5.0% | 2.5% | 5.3% |
|  | both | stage 8 | 6.4% | 6.7% | 10.6% | 5.9% |
|  | both | stage 10 | 6.3% | 5.0% | 4.0% | 4.5% |
| intergenic | A | stage 8 | 11.2% | 11.8% | 3.8% | 10.9% |
|  | A | stage 10 | 10.4% | 11.0% | 13.9% | 10.1% |
|  | B | stage 8 | 10.9% | 8.5% | 11.5% | 10.3% |
|  | B | stage 10 | 11.2% | 9.4% | 5.2% | 9.5% |
|  | both | stage 8 | 9.7% | 7.9% | 4.0% | 9.7% |
|  | both | stage 10 | 10.6% | 8.1% | 2.7% | 7.1% |
| intronic | A | stage 8 | 25.4% | 23.8% | 12.8% | 23.4% |
|  | A | stage 10 | 24.7% | 24.4% | 32.1% | 23.6% |
|  | B | stage 8 | 24.0% | 23.6% | 24.7% | 24.0% |
|  | B | stage 10 | 27.0% | 31.6% | 36.2% | 23.5% |
|  | both | stage 8 | 25.0% | 24.9% | 17.3% | 27.1% |
|  | both | stage 10 | 25.6% | 30.0% | 38.7% | 32.6% |
| non-synonymous | A | stage 8 | 4.4% | 3.7% | 0.0% | 3.2% |
|  | A | stage 10 | 4.2% | 3.6% | 1.3% | 3.1% |
|  | B | stage 8 | 4.4% | 4.1% | 5.2% | 3.2% |
|  | B | stage 10 | 3.8% | 3.0% | 3.9% | 4.1% |
|  | both | stage 8 | 3.9% | 3.7% | 3.3% | 3.2% |
|  | both | stage 10 | 3.8% | 3.5% | 2.7% | 2.9% |
| synonymous | A | stage 8 | 11.0% | 10.7% | 11.4% | 13.0% |
|  | A | stage 10 | 11.4% | 11.6% | 7.8% | 13.1% |
|  | B | stage 8 | 9.8% | 10.5% | 6.6% | 12.2% |
|  | B | stage 10 | 8.5% | 7.4% | 7.8% | 10.6% |
|  | both | stage 8 | 10.6% | 10.0% | 8.0% | 10.8% |
|  | both | stage 10 | 10.3% | 9.0% | 5.3% | 10.4% |
| upstream/downstream | A | stage 8 | 41.6% | 43.1% | 60.8% | 41.8% |
|  | A | stage 10 | 42.6% | 43.2% | 39.7% | 42.7% |
|  | B | stage 8 | 44.2% | 45.2% | 42.1% | 44.3% |
|  | B | stage 10 | 43.0% | 43.5% | 44.2% | 46.8% |
|  | both | stage 8 | 44.1% | 46.8% | 56.0% | 42.7% |
|  | both | stage 10 | 43.2% | 44.1% | 44.7% | 41.8% |

**S5 Table:** **Enrichment of functional annotations in diapause-associated SNPs.** For each class of variant, the proportion of diapause-associated SNPs assigned to that variant type was quantified for each imputation and permutation. The numerical values are the quantile rank of the median of the 100 observed imputations relative to the distribution of the permutations. Quantile ranks below 5% and above 95% (i.e. those values that are de-enriched or enriched relative to permutations) are highlighted in bold.

| Annotation | Population | phenotype | Top 1% | Top 0.1% | Top 0.01% | LASSO |
| --- | --- | --- | --- | --- | --- | --- |
| UTR | A | stage 8 | 0.43 | 0.58 | 0.82 | 0.67 |
|  | A | stage 10 | 0.77 | 0.61 | 0.27 | 0.82 |
|  | B | stage 8 | 0.86 | 0.9 | 0.86 | 0.51 |
|  | B | stage 10 | 0.56 | 0.2 | 0.09 | 0.44 |
|  | both | stage 8 | 0.736 | 0.677 | 0.875 | 0.487 |
|  | both | stage 10 | 0.676 | 0.179 | 0.362 | 0.301 |
| intergenic | A | stage 8 | 0.64 | 0.71 | 0.12 | 0.55 |
|  | A | stage 10 | 0.43 | 0.62 | 0.79 | 0.43 |
|  | B | stage 8 | 0.58 | 0.11 | 0.53 | 0.45 |
|  | B | stage 10 | 0.61 | 0.27 | 0.19 | 0.35 |
|  | both | stage 8 | 0.124 | 0.109 | 0.129 | 0.321 |
|  | both | stage 10 | 0.448 | 0.118 | 0.073 | 0.144 |
| intronic | A | stage 8 | 0.76 | 0.46 | 0.08 | 0.41 |
|  | A | stage 10 | 0.44 | 0.41 | 0.84 | 0.35 |
|  | B | stage 8 | 0.31 | 0.32 | 0.59 | 0.32 |
|  | B | stage 10 | **0.97** | 0.94 | 0.95 | 0.37 |
|  | both | stage 8 | 0.653 | 0.552 | 0.214 | 0.742 |
|  | both | stage 10 | 0.816 | **0.963** | 0.95 | 0.917 |
| non-synonymous | A | stage 8 | 0.8 | 0.42 | 0.1 | 0.44 |
|  | A | stage 10 | 0.62 | 0.3 | 0.18 | 0.44 |
|  | B | stage 8 | 0.8 | 0.6 | 0.78 | 0.37 |
|  | B | stage 10 | 0.27 | 0.2 | 0.63 | 0.57 |
|  | both | stage 8 | 0.371 | 0.412 | 0.443 | 0.356 |
|  | both | stage 10 | 0.301 | 0.312 | 0.456 | 0.371 |
| synonymous | A | stage 8 | 0.32 | 0.28 | 0.42 | 0.54 |
|  | A | stage 10 | 0.55 | 0.56 | 0.32 | 0.61 |
|  | B | stage 8 | **0.01** | 0.35 | 0.15 | 0.67 |
|  | B | stage 10 | **0.01** | **0.03** | 0.27 | 0.39 |
|  | both | stage 8 | 0.085 | 0.176 | 0.272 | 0.36 |
|  | both | stage 10 | **0.031** | 0.067 | 0.107 | 0.365 |
| upstream/downstream | A | stage 8 | 0.14 | 0.54 | **0.97** | 0.48 |
|  | A | stage 10 | 0.51 | 0.57 | 0.35 | 0.61 |
|  | B | stage 8 | 0.89 | 0.83 | 0.46 | 0.61 |
|  | B | stage 10 | 0.52 | 0.64 | 0.55 | 0.84 |
|  | both | stage 8 | 0.872 | 0.914 | **0.952** | 0.547 |
|  | both | stage 10 | 0.661 | 0.671 | 0.609 | 0.482 |

Table. S6 List of LASSO SNPs, top 0.01% SNPs, and top 0.1% SNPs along with annotations. ***(Excel file)***

## Notes

### Competing Interest Statement

The authors have declared no competing interest.

## References

1. Kawecki TJ, Ebert D. Conceptual issues in local adaptation. Ecol Lett. 2004;7: 1225–1241. doi:10.1111/j.1461-0248.2004.00684.x

2. Paul MJ, Zucker Irving, Schwartz William J. Tracking the seasons: the internal calendars of vertebrates. Philos Trans R Soc B Biol Sci. 2008;363: 341–361. doi:10.1098/rstb.2007.2143

3. Andrés F, Coupland G. The genetic basis of flowering responses to seasonal cues. Nat Rev Genet. 2012;13: 627–639. doi:10.1038/nrg3291

4. Denlinger DL, Hahn DA, Merlin C, Holzapfel CM, Bradshaw WE. Keeping time without a spine: what can the insect clock teach us about seasonal adaptation? Phil Trans R Soc B. 2017;372: 20160257. doi:10.1098/rstb.2016.0257

5. Moran NA. The Evolution of Aphid Life Cycles. Annu Rev Entomol. 1992;37: 321– 348. doi:10.1146/annurev.en.37.010192.001541

6. Nijhout HF. Development and evolution of adaptive polyphenisms. Evol Dev. 2003;5: 9–18. doi:10.1046/j.1525-142X.2003.03003.x

7. Canard M. Seasonal adaptations of green lacewings (Neuroptera: Chrysopidae). Eur J Entomol Ceske Budejovice. 2005;102: 317.

8. Urquhart FA, Urquhart NR. Autumnal migration routes of the eastern population of the monarch butterfly (Danaus p. plexippus L.; Danaidae; Lepidoptera) in North America to the overwintering site in the Neovolcanic Plateau of Mexico. Can J Zool. 1978;56: 1759–1764. doi:10.1139/z78-240

9. Tauber MJ, Tauber CA, Masaki S. Seasonal Adaptations of Insects. Oxford University Press; 1986.

10. Denlinger DL. Regulation of Diapause. Annu Rev Entomol. 2002;47: 93–122. doi:10.1146/annurev.ento.47.091201.145137

11. Koštál V. Eco-physiological phases of insect diapause. J Insect Physiol. 2006;52: 113–127. doi:10.1016/j.jinsphys.2005.09.008

12. Schmidt P. Evolution and mechanisms of insect reproductive diapause: a plastic and pleiotropic life history syndrome. Mechanisms of Life History Evolution: The Genetics and Physiology of Life History Traits and Trade-Offs. 2011. pp. 221–229.

13. Tougeron K. Diapause research in insects: historical review and recent work perspectives. Entomol Exp Appl. 2019;167: 27–36. doi:10.1111/eea.12753

14. Bradshaw WE, Armbruster PA, Holzapfel CM. Fitness Consequences of Hibernal Diapause in the Pitcher-Plant Mosquito, Wyeomyia Smithii. Ecology. 1998;79: 1458–1462. doi:10.1890/0012-9658(1998)079[1458:FCOHDI]2.0.CO;2

15. Schmidt PS, Paaby AB, Heschel MS. Genetic variance for diapause expression and associated life histories in Drosophila melanogaster. Evolution. 2005;59: 2616–2625.

16. Chen C, Xia Q-W, Xiao H-J, Xiao L, Xue F-S. A comparison of the life-history traits between diapause and direct development individuals in the cotton bollworm, Helicoverpa armigera. J Insect Sci. 2014;14. doi:10.1093/jis/14.1.19

17. Bradshaw WE, Lounibos LP. Evolution of Dormancy and Its Photoperiodic Control in Pitcher-Plant Mosquitoes. Evolution. 1977;31: 546–567. doi:10.1111/j.1558-5646.1977.tb01044.x

18. Schmidt PS, Matzkin L, Ippolito M, Eanes WF. Geographic Variation in Diapause Incidence, Life-History Traits, and Climatic Adaptation in Drosophila Melanogaster. Evolution. 2005;59: 1721–1732. doi:10.1111/j.0014-3820.2005.tb01821.x

19. Schmidt PS, Conde DR. Environmental Heterogeneity and the Maintenance of Genetic Variation for Reproductive Diapause in Drosophila Melanogaster. Evolution. 2006;60: 1602–1611. doi:10.1111/j.0014-3820.2006.tb00505.x

20. Paolucci S, Zande L van de, Beukeboom LW. Adaptive latitudinal cline of photoperiodic diapause induction in the parasitoid Nasonia vitripennis in Europe. J Evol Biol. 2013;26: 705–718. doi:10.1111/jeb.12113

21. Posledovich D, Toftegaard T, Wiklund C, Ehrlén J, Gotthard K. Latitudinal variation in diapause duration and post-winter development in two pierid butterflies in relation to phenological specialization. Oecologia. 2015;177: 181– 190. doi:10.1007/s00442-014-3125-1

22. Lehmann P, Lyytinen A, Piiroinen S, Lindström L. Latitudinal differences in diapause related photoperiodic responses of European Colorado potato beetles (Leptinotarsa decemlineata). Evol Ecol. 2015;29: 269–282. doi:10.1007/s10682-015-9755-x

23. Klepsatel P, Gáliková M, Maio ND, Ricci S, Schlötterer C, Flatt T. Reproductive and post-reproductive life history of wild-caught Drosophila melanogaster under laboratory conditions. J Evol Biol. 2013;26: 1508–1520. doi:10.1111/jeb.12155

24. Saunders DS, Henrich VC, Gilbert LI. Induction of diapause in Drosophila melanogaster: photoperiodic regulation and the impact of arrhythmic clock mutations on time measurement. Proc Natl Acad Sci U S A. 1989;86: 3748–3752.

25. Saunders DS, Richard DS, Applebaum SW, Ma M, Gilbert LI. Photoperiodic diapause in Drosophila melanogaster involves a block to the juvenile hormone regulation of ovarian maturation. Gen Comp Endocrinol. 1990;79: 174–184. doi:10.1016/0016-6480(90)90102-R

26. Saunders DS. Insect Clocks, Third Edition. Elsevier; 2002.

27. Williams KD, Busto M, Suster ML, So AK-C, Ben-Shahar Y, Leevers SJ, et al. Natural variation in Drosophila melanogaster diapause due to the insulin-regulated PI3-kinase. Proc Natl Acad Sci. 2006;103: 15911–15915. doi:10.1073/pnas.0604592103

28. Liu Y, Liao S, Veenstra JA, Nässel DR. *Drosophila* insulin-like peptide 1 (DILP1) is transiently expressed during non-feeding stages and reproductive dormancy. Sci Rep. 2016;6: 26620. doi:10.1038/srep26620

29. Schiesari L, Andreatta G, Kyriacou CP, O’Connor MB, Costa R. The Insulin-Like Proteins dILPs-2/5 Determine Diapause Inducibility in Drosophila. PLOS ONE. 2016;11: e0163680. doi:10.1371/journal.pone.0163680

30. Richard DS, Jones JM, Barbarito MR, Stacy Cerula, Detweiler JP, Fisher SJ, et al. Vitellogenesis in diapausing and mutant Drosophila melanogaster: further evidence for the relative roles of ecdysteroids and juvenile hormones. J Insect Physiol. 2001;47: 905–913. doi:10.1016/S0022-1910(01)00063-4

31. Richard DS, Rybczynski R, Wilson TG, Wang Y, Wayne ML, Zhou Y, et al. Insulin signaling is necessary for vitellogenesis in Drosophila melanogaster independent of the roles of juvenile hormone and ecdysteroids: female sterility of the chico1 insulin signaling mutation is autonomous to the ovary. J Insect Physiol. 2005;51: 455–464. doi:10.1016/j.jinsphys.2004.12.013

32. Gilbert LI, Serafin RB, Watkins NL, Richard DS. Ecdysteroids regulate yolk protein uptake by Drosophila melanogaster oocytes. J Insect Physiol. 1998;44: 637–644. doi:10.1016/s0022-1910(98)00020-1

33. Andreatta G, Kyriacou CP, Flatt T, Costa R. Aminergic Signaling Controls Ovarian Dormancy in Drosophila. Sci Rep. 2018;8: 2030. doi:10.1038/s41598-018-20407-z

34. Kubrak OI, Ku erová L, Theopold U, Nässel DR. The Sleeping Beauty: How Reproductive D^č^ iapause Affects Hormone Signaling, Metabolism, Immune Response and Somatic Maintenance in Drosophila melanogaster. PLOS ONE. 2014;9: e113051. doi:10.1371/journal.pone.0113051

35. Lirakis M, Dolezal M, Schlötterer C. Redefining reproductive dormancy in Drosophila as a general stress response to cold temperatures. J Insect Physiol. 2018;107: 175–185. doi:10.1016/j.jinsphys.2018.04.006

36. Ojima N, Hara Y, Ito H, Yamamoto D. Genetic dissection of stress-induced reproductive arrest in Drosophila melanogaster females. PLOS Genet. 2018;14: e1007434. doi:10.1371/journal.pgen.1007434

37. Zhao X, Bergland AO, Behrman EL, Gregory BD, Petrov DA, Schmidt PS. Global transcriptional profiling of diapause and climatic adaptation in Drosophila melanogaster. Mol Biol Evol. 2015; msv263. doi:10.1093/molbev/msv263

38. Ku erová L, Kubrak OI, Bengtsson JM, Strnad H, Nylin S, Theopold U, et al. Slo^č^wed aging during reproductive dormancy is reflected in genome-wide transcriptome changes in Drosophila melanogaster. BMC Genomics. 2016;17: 50. doi:10.1186/s12864-016-2383-1

39. Ragland GJ, Keep E. Comparative transcriptomics support evolutionary convergence of diapause responses across Insecta. Physiol Entomol. 2017; n/a- n/a. doi:10.1111/phen.12193

40. Parker DJ, Ritchie MG, Kankare M. Preparing for Winter: The Transcriptomic Response Associated with Different Day Lengths in Drosophila montana. G3 Genes Genomes Genet. 2016;6: 1373–1381. doi:10.1534/g3.116.027870

41. Kankare M, Parker DJ, Merisalo M, Salminen TS, Hoikkala A. Transcriptional Differences between Diapausing and Non-Diapausing D. montana Females Reared under the Same Photoperiod and Temperature. PLOS ONE. 2016;11: e0161852. doi:10.1371/journal.pone.0161852

42. Zhai Y, Dong X, Gao H, Chen H, Yang P, Li P, et al. Quantitative Proteomic and Transcriptomic Analyses of Metabolic Regulation of Adult Reproductive Diapause in Drosophila suzukii (Diptera: Drosophilidae) Females. Front Physiol. 2019;10. doi:10.3389/fphys.2019.00344

43. Kubrak OI, Ku erová L, Theopold U, Nylin S, Nässel DR. Characterization of Reproductive D^č^ ormancy in Male Drosophila melanogaster. Front Physiol. 2016;7. doi:10.3389/fphys.2016.00572

44. Izquierdo JI. How does Drosophila melanogaster overwinter? Entomol Exp Appl. 1991;59: 51–58. doi:10.1111/j.1570-7458.1991.tb01485.x

45. Machado HE, Bergland AO, O’Brien KR, Behrman EL, Schmidt PS, Petrov DA. Comparative population genomics of latitudinal variation in Drosophila simulans and Drosophila melanogaster. Mol Ecol. 2016;25: 723–740. doi:10.1111/mec.13446

46. Ohtsu T, Kimura MT, Hori SH. Energy storage during reproductive diapause in the Drosophila melanogaster species group. J Comp Physiol B. 1992;162: 203–208. doi:10.1007/BF00357524

47. Higuchi C, Kimura MT. Influence of photoperiod on low temperature acclimation for cold-hardiness in Drosophila auraria. Physiol Entomol. 1985;10: 303–308. doi:10.1111/j.1365-3032.1985.tb00051.x

48. Lumme J, Oikarinen A. The genetic basis of the geographically variable photoperiodic diapause in Drosophila littoralis. Hereditas. 1977;86: 129–141. doi:10.1111/j.1601-5223.1977.tb01221.x

49. Lumme J. Phenology and Photoperiodic Diapause in Northern Populations of Drosophila. In: Dingle H, editor. Evolution of Insect Migration and Diapause. New York, NY: Springer US; 1978. pp. 145–170. doi:10.1007/978-1-4615-6941-1_7

50. Reis M, Valer FB, Vieira CP, Vieira J. Drosophila americana Diapausing Females Show Features Typical of Young Flies. PLOS ONE. 2015;10: e0138758. doi:10.1371/journal.pone.0138758

51. Tyukmaeva VI, Salminen TS, Kankare M, Knott KE, Hoikkala A. Adaptation to a seasonally varying environment: a strong latitudinal cline in reproductive diapause combined with high gene flow in Drosophila montana. Ecol Evol. 2011;1: 160– 168. doi:10.1002/ece3.14

52. Schmidt PS, Zhu C-T, Das J, Batavia M, Yang L, Eanes WF. An amino acid polymorphism in the couch potato gene forms the basis for climatic adaptation in Drosophila melanogaster. Proc Natl Acad Sci U S A. 2008;105: 16207–16211. doi:10.1073/pnas.0805485105

53. Bergland AO, Behrman EL, O’Brien KR, Schmidt PS, Petrov DA. Genomic Evidence of Rapid and Stable Adaptive Oscillations over Seasonal Time Scales in Drosophila. PLoS Genet. 2014;10: e1004775. doi:10.1371/journal.pgen.1004775

54. Cogni R, Kuczynski C, Koury S, Lavington E, Behrman EL, O’Brien KR, et al. The intensity of selection acting on the couch potato gene--spatial-temporal variation in a diapause cline. Evol Int J Org Evol. 2014;68: 538–548. doi:10.1111/evo.12291

55. Pool JE. The Mosaic Ancestry of the Drosophila Genetic Reference Panel and the D. melanogaster Reference Genome Reveals a Network of Epistatic Fitness Interactions. Mol Biol Evol. 2015;32: 3236–3251. doi:10.1093/molbev/msv194

56. Hsu S-K, Jakši AM, Nolte V, Lirakis M, Kofler R, Barghi N, et al. Rapid sex-specific adapta^ć^tion to high temperature in Drosophila. Tautz D, Ebert D, Reisser C, editors. eLife. 2020;9: e53237. doi:10.7554/eLife.53237

57. Lankinen P, Tyukmaeva VI, Hoikkala A. Northern Drosophila montana flies show variation both within and between cline populations in the critical day length evoking reproductive diapause. J Insect Physiol. 2013;59: 745–751. doi:10.1016/j.jinsphys.2013.05.006

58. Kimura MT. Quantitative response to photoperiod during reproductive diapause in the Drosophila auraria species-complex. J Insect Physiol. 1990;36: 147–152. doi:10.1016/0022-1910(90)90115-V

59. Tatar M, Chien SA, Priest NK. Negligible Senescence during Reproductive Dormancy in Drosophila melanogaster. Am Nat. 2001;158: 248–258. doi:10.1086/321320

60. Tauber E, Zordan M, Sandrelli F, Pegoraro M, Osterwalder N, Breda C, et al. Natural Selection Favors a Newly Derived timeless Allele in Drosophila melanogaster. Science. 2007;316: 1895–1898. doi:10.1126/science.1138412

61. Levins R. Evolution in Changing Environments: Some Theoretical Explorations. Princeton University Press; 1968.

62. Ragland GJ, Armbruster PA, Meuti ME. Evolutionary and functional genetics of insect diapause: a call for greater integration. Curr Opin Insect Sci. 2019;36: 74– 81. doi:10.1016/j.cois.2019.08.003

63. Weller CA, Bergland AO. Accurate, ultra-low coverage genome reconstruction and association studies in Hybrid Swarm mapping populations. bioRxiv. 2019; 671925. doi:10.1101/671925

64. Becker R, Wilks A, Brownrigg R, Minka T, Deckmyn A. maps: Draw Geographical Maps. 2018. Available: https://CRAN.R-project.org/package=maps

65. Middleton CA, Nongthomba U, Parry K, Sweeney ST, Sparrow JC, Elliott CJ. Neuromuscular organization and aminergic modulation of contractions in the Drosophila ovary. BMC Biol. 2006;4: 17. doi:10.1186/1741-7007-4-17

66. King RC. Ovarian development in Drosophila melanogaster. Academic Press; 1970.

67. Lee SF, Sgrò CM, Shirriffs J, Wee CW, Rako L, van Heerwaarden B, et al. Polymorphism in the couch potato gene clines in eastern Australia but is not associated with ovarian dormancy in Drosophila melanogaster. Mol Ecol. 2011;20: 2973–2984. doi:10.1111/j.1365-294X.2011.05155.x

68. Soller M, Bownes M, Kubli E. Mating and sex peptide stimulate the accumulation of yolk in oocytes of Drosophila melanogaster. Eur J Biochem. 1997;243: 732– 738.

69. Soller M, Bownes M, Kubli E. Control of oocyte maturation in sexually mature Drosophila females. Dev Biol. 1999;208: 337–351. doi:10.1006/dbio.1999.9210

70. Mirth CK, Nogueira Alves A, Piper MD. Turning food into eggs: insights from nutritional biology and developmental physiology of Drosophila. Curr Opin Insect Sci. 2019;31: 49–57. doi:10.1016/j.cois.2018.08.006

71. Yang J, Lee SH, Goddard ME, Visscher PM. GCTA: a tool for genome-wide complex trait analysis. Am J Hum Genet. 2011;88: 76–82. doi:10.1016/j.ajhg.2010.11.011

72. Yang J, Benyamin B, McEvoy BP, Gordon S, Henders AK, Nyholt DR, et al. Common SNPs explain a large proportion of the heritability for human height. Nat Genet. 2010;42: 565–569. doi:10.1038/ng.608

73. Tibshirani R. Regression Shrinkage and Selection via the Lasso. J R Stat Soc Ser B Methodol. 1996;58: 267–288.

74. Wu TT, Chen YF, Hastie T, Sobel E, Lange K. Genome-wide association analysis by lasso penalized logistic regression. Bioinformatics. 2009;25: 714–721. doi:10.1093/bioinformatics/btp041

75. Sing T, Sander O, Beerenwinkel N, Lengauer T. ROCR: visualizing classifier performance in R. Bioinformatics. 2005;21: 3940–3941. doi:10.1093/bioinformatics/bti623

76. Sandrelli F, Tauber E, Pegoraro M, Mazzotta G, Cisotto P, Landskron J, et al. A Molecular Basis for Natural Selection at the timeless Locus in Drosophila melanogaster. Science. 2007;316: 1898–1900. doi:10.1126/science.1138426

77. Mackay TFC, Richards S, Stone EA, Barbadilla A, Ayroles JF, Zhu D, et al. The Drosophila melanogaster Genetic Reference Panel. Nature. 2012;482: 173–178. doi:10.1038/nature10811

78. Machado HE, Bergland AO, Taylor R, Tilk S, Behrman E, Dyer K, et al. Broad geographic sampling reveals predictable and pervasive seasonal adaptation in Drosophila. bioRxiv. 2019; 337543. doi:10.1101/337543

79. Berg JJ, Coop G. A Population Genetic Signal of Polygenic Adaptation. PLOS Genet. 2014;10: e1004412. doi:10.1371/journal.pgen.1004412

80. Beissinger T, Kruppa J, Cavero D, Ha N-T, Erbe M, Simianer H. A Simple Test Identifies Selection on Complex Traits. Genetics. 2018;209: 321–333. doi:10.1534/genetics.118.300857

81. Pais IS, Valente RS, Sporniak M, Teixeira L. Drosophila melanogaster establishes a species-specific mutualistic interaction with stable gut-colonizing bacteria. PLOS Biol. 2018;16: e2005710. doi:10.1371/journal.pbio.2005710

82. Stone HM, Erickson PA, Bergland AO. Phenotypic plasticity, but not adaptive tracking, underlies seasonal variation in post-cold hardening freeze tolerance of Drosophila melanogaster. Ecol Evol. 2020;10: 217–231. doi:10.1002/ece3.5887

83. Drummond-Barbosa D, Spradling AC. Stem Cells and Their Progeny Respond to Nutritional Changes during Drosophila Oogenesis. Dev Biol. 2001;231: 265–278. doi:10.1006/dbio.2000.0135

84. Terashima J, Bownes M. Translating Available Food Into the Number of Eggs Laid by Drosophila melanogaster. Genetics. 2004;167: 1711–1719. doi:10.1534/genetics.103.024323

85. Terashima J, Takaki K, Sakurai S, Bownes M. Nutritional status affects 20- hydroxyecdysone concentration and progression of oogenesis in Drosophila melanogaster. J Endocrinol. 2005;187: 69–79. doi:10.1677/joe.1.06220

86. Lee KP, Simpson SJ, Clissold FJ, Brooks R, Ballard JWO, Taylor PW, et al. Lifespan and reproduction in Drosophila: New insights from nutritional geometry. Proc Natl Acad Sci. 2008;105: 2498–2503. doi:10.1073/pnas.0710787105

87. Fabian DK, Lack JB, Mathur V, Schlötterer C, Schmidt PS, Pool JE, et al. Spatially varying selection shapes life history clines among populations of Drosophila melanogaster from sub-Saharan Africa. J Evol Biol. 2015;28: 826–840. doi:10.1111/jeb.12607

88. Zonato V, Collins L, Pegoraro M, Tauber E, Kyriacou CP. Is diapause an ancient adaptation in Drosophila? J Insect Physiol. 2017;98: 267–274. doi:10.1016/j.jinsphys.2017.01.017

89. Lack JB, Cardeno CM, Crepeau MW, Taylor W, Corbett-Detig RB, Stevens KA, et al. The Drosophila Genome Nexus: A Population Genomic Resource of 623 Drosophila melanogaster Genomes, Including 197 from a Single Ancestral Range Population. Genetics. 2015;199: 1229–1241. doi:10.1534/genetics.115.174664

90. Lack JB, Lange JD, Tang AD, Corbett-Detig RB, Pool JE. A Thousand Fly Genomes: An Expanded Drosophila Genome Nexus. Mol Biol Evol. 2016; msw195. doi:10.1093/molbev/msw195

91. Voight BF, Kudaravalli S, Wen X, Pritchard JK. A Map of Recent Positive Selection in the Human Genome. PLOS Biol. 2006;4: e72. doi:10.1371/journal.pbio.0040072

92. Rockman MV. The QTN Program and the Alleles That Matter for Evolution: All That’s Gold Does Not Glitter. Evol Int J Org Evol. 2012;66: 1–17. doi:10.1111/j.1558-5646.2011.01486.x

93. Boyle EA, Li YI, Pritchard JK. An Expanded View of Complex Traits: From Polygenic to Omnigenic. Cell. 2017;169: 1177–1186. doi:10.1016/j.cell.2017.05.038

94. Emerson KJ, Uyemura AM, McDaniel KL, Schmidt PS, Bradshaw WE, Holzapfel CM. Environmental control of ovarian dormancy in natural populations of Drosophila melanogaster. J Comp Physiol A. 2009;195: 825–829. doi:10.1007/s00359-009-0460-5

95. Anduaga AM, Nagy D, Costa R, Kyriacou CP. Diapause in Drosophila melanogaster – Photoperiodicity, cold tolerance and metabolites. J Insect Physiol. 2018;105: 46–53. doi:10.1016/j.jinsphys.2018.01.003

96. Pegoraro M, Tauber E. Photoperiod-dependent expression of MicroRNA in Drosophila. bioRxiv. 2018; 464180. doi:10.1101/464180

97. Nagy D, Andreatta G, Bastianello S, Martín Anduaga A, Mazzotta G, Kyriacou CP, et al. A Semi-natural Approach for Studying Seasonal Diapause in Drosophila melanogaster Reveals Robust Photoperiodicity. J Biol Rhythms. 2018;33: 117– 125. doi:10.1177/0748730417754116

98. McCall K. Eggs over easy: cell death in the Drosophila ovary. Dev Biol. 2004;274: 3–14. doi:10.1016/j.ydbio.2004.07.017

99. Pegoraro M, Zonato V, Tyler ER, Fedele G, Kyriacou CP, Tauber E. Geographical analysis of diapause inducibility in European Drosophila melanogaster populations. J Insect Physiol. 2017;98: 238–244. doi:10.1016/j.jinsphys.2017.01.015

100. Rajpurohit S, Hanus R, Vrkoslav V, Behrman EL, Bergland AO, Petrov D, et al. Adaptive dynamics of cuticular hydrocarbons in Drosophila. J Evol Biol. 2017;30: 66–80. doi:10.1111/jeb.12988

101. Friedline CJ, Faske TM, Lind BM, Hobson EM, Parry D, Dyer RJ, et al. Evolutionary genomics of gypsy moth populations sampled along a latitudinal gradient. Mol Ecol. 2019;0. doi:10.1111/mec.15069

102. Bay RA, Palumbi SR. Multilocus Adaptation Associated with Heat Resistance in Reef-Building Corals. Curr Biol. 2014;24: 2952–2956. doi:10.1016/j.cub.2014.10.044

103. Exposito-Alonso M, Vasseur F, Ding W, Wang G, Burbano HA, Weigel D. Genomic basis and evolutionary potential for extreme drought adaptation in Arabidopsis thaliana. Nat Ecol Evol. 2018;2: 352. doi:10.1038/s41559-017-0423-0

104. Mansourian S, Enjin A, Jirle EV, Ramesh V, Rehermann G, Becher PG, et al. Wild African Drosophila melanogaster Are Seasonal Specialists on Marula Fruit. Curr Biol. 2018 [cited 13 Dec 2018]. doi:10.1016/j.cub.2018.10.033

105. Wilson TG. Determinants of oöcyte degeneration in Drosophila melanogaster. J Insect Physiol. 1985;31: 109–117. doi:10.1016/0022-1910(85)90015-0

106. Gruntenko NE, Rauschenbach IY. Interplay of JH, 20E and biogenic amines under normal and stress conditions and its effect on reproduction. J Insect Physiol. 2008;54: 902–908. doi:10.1016/j.jinsphys.2008.04.004

107. Nosil P, Villoutreix R, Carvalho CF de, Farkas TE, Soria-Carrasco V, Feder JL, et al. Natural selection and the predictability of evolution in Timema stick insects. Science. 2018;359: 765–770. doi:10.1126/science.aap9125

108. Barghi N, Tobler R, Nolte V, Jakši AM, Mallard F, Otte KA, et al. Genetic redundancy fuels polygenic adapta^ć^tion in Drosophila. PLOS Biol. 2019;17: e3000128. doi:10.1371/journal.pbio.3000128

109. Pitchers W, Pool JE, Dworkin I. Altitudinal Clinal Variation in Wing Size & Shape in African Drosophila melanogaster: One Cline or Many? Evol Int J Org Evol. 2013;67: 438–452. doi:10.1111/j.1558-5646.2012.01774.x

110. Klepsatel P, Gáliková M, Huber CD, Flatt T. Similarities and Differences in Altitudinal Versus Latitudinal Variation for Morphological Traits in Drosophila Melanogaster. Evolution. 2014;68: 1385–1398. doi:10.1111/evo.12351

111. Savolainen O, Lascoux M, Merilä J. Ecological genomics of local adaptation. Nat Rev Genet. 2013;14: 807–820. doi:10.1038/nrg3522

112. Hoban S, Kelley JL, Lotterhos KE, Antolin MF, Bradburd G, Lowry DB, et al. Finding the Genomic Basis of Local Adaptation: Pitfalls, Practical Solutions, and Future Directions. Am Nat. 2016;188: 379–397. doi:10.1086/688018

113. Bale JS, Hayward S a. L. Insect overwintering in a changing climate. J Exp Biol. 2010;213: 980–994. doi:10.1242/jeb.037911

114. Doležel D, Van ková H, Šauman I, Hodkova M. Is period gene causally involved in the photoper^ě^io^č^dic regulation of reproductive diapause in the linden bug, Pyrrhocoris apterus? J Insect Physiol. 2005;51: 655–659. doi:10.1016/j.jinsphys.2005.01.009

115. Han B, Denlinger DL. Mendelian Inheritance of Pupal Diapause in the Flesh Fly, Sarcophaga bullata. J Hered. 2009;100: 251–255. doi:10.1093/jhered/esn082

116. Kim Y, Krafsur ES, Bailey TB, Zhao S. Mode of inheritance of face fly diapause and its correlation with other developmental traits. Ecol Entomol. 1995;20: 359– 366. doi:10.1111/j.1365-2311.1995.tb00468.x

117. Ikten C, Skoda SR, Hunt TE, Molina-Ochoa J, Foster JE. Genetic Variation and Inheritance of Diapause Induction in Two Distinct Voltine Ecotypes of Ostrinia nubilalis (Lepidoptera: Crambidae). Ann Entomol Soc Am. 2011;104: 567–575. doi:10.1603/AN09149

118. Kozak GM, Wadsworth CB, Kahne SC, Bogdanowicz SM, Harrison RG, Coates BS, et al. Genomic Basis of Circannual Rhythm in the European Corn Borer Moth. Curr Biol. 2019;29: 3501–3509.e5. doi:10.1016/j.cub.2019.08.053

119. Mori A, Romero-Severson J, Severson DW. Genetic basis for reproductive diapause is correlated with life history traits within the Culex pipiens complex. Insect Mol Biol. 2007;16: 515–524. doi:10.1111/j.1365-2583.2007.00746.x

120. Pruisscher P, Nylin S, Gotthard K, Wheat CW. Genetic variation underlying local adaptation of diapause induction along a cline in a butterfly. Mol Ecol. 2018;27: 3613–3626. doi:10.1111/mec.14829

121. Grenier JK, Arguello JR, Moreira MC, Gottipati S, Mohammed J, Hackett SR, et al. Global diversity lines - a five-continent reference panel of sequenced Drosophila melanogaster strains. G3 Bethesda Md. 2015;5: 593–603. doi:10.1534/g3.114.015883

122. Behrman E, Howick H, Kapun M, Staubach F, Bergland B, Petrov D, et al. Rapid seasonal evolution in innate immunity of wild Drosophila melanogaster. Proc R Soc B Biol Sci. 2018;285: 20172599. doi:10.1098/rspb.2017.2599

123. Kao JY, Zubair A, Salomon MP, Nuzhdin SV, Campo D. Population genomic analysis uncovers African and European admixture in Drosophila melanogaster populations from the south-eastern United States and Caribbean Islands. Mol Ecol. 2015;24: 1499–1509. doi:10.1111/mec.13137

124. Fox J, Weisberg S. An R Companion to Applied Regression, Third Edition. Thousand Oaks, CA: Sage; 2019. Available: https://socialsciences.mcmaster.ca/jfox/Books/Companion/

125. Baym M, Kryazhimskiy S, Lieberman TD, Chung H, Desai MM, Kishony R. Inexpensive Multiplexed Library Preparation for Megabase-Sized Genomes. PLOS ONE. 2015;10: e0128036. doi:10.1371/journal.pone.0128036

126. Zhang J, Kobert K, Flouri T, Stamatakis A. PEAR: a fast and accurate Illumina Paired-End reAd mergeR. Bioinforma Oxf Engl. 2014;30: 614–620. doi:10.1093/bioinformatics/btt593

127. Li H, Durbin R. Fast and accurate short read alignment with Burrows-Wheeler transform. Bioinforma Oxf Engl. 2009;25: 1754–1760. doi:10.1093/bioinformatics/btp324

128. McKenna A, Hanna M, Banks E, Sivachenko A, Cibulskis K, Kernytsky A, et al. The Genome Analysis Toolkit: a MapReduce framework for analyzing next-generation DNA sequencing data. Genome Res. 2010;20: 1297–1303. doi:10.1101/gr.107524.110

129. Kessner D, Turner TL, Novembre J. Maximum likelihood estimation of frequencies of known haplotypes from pooled sequence data. Mol Biol Evol. 2013;30: 1145– 1158. doi:10.1093/molbev/mst016

130. Zheng C, Boer MP, van Eeuwijk FA. Reconstruction of Genome Ancestry Blocks in Multiparental Populations. Genetics. 2015;200: 1073–1087. doi:10.1534/genetics.115.177873

131. Comeron JM, Ratnappan R, Bailin S. The Many Landscapes of Recombination in Drosophila melanogaster. PLOS Genet. 2012;8: e1002905. doi:10.1371/journal.pgen.1002905

132. Zheng X, Levine D, Shen J, Gogarten SM, Laurie C, Weir BS. A high-performance computing toolset for relatedness and principal component analysis of SNP data. Bioinforma Oxf Engl. 2012;28: 3326–3328. doi:10.1093/bioinformatics/bts606

133. Kapun M, Fabian DK, Goudet J, Flatt T. Genomic Evidence for Adaptive Inversion Clines in Drosophila melanogaster. Mol Biol Evol. 2016;33: 1317–1336. doi:10.1093/molbev/msw016

134. Conomos M, Gogarten SM, Brown L, Chen H, Rice K, Sofer T, et al. GENetic EStimation and Inference in Structured samples (GENESIS): Statistical methods for analyzing genetic data from samples with population structure and/or relatedness. 2019. Available: https://github.com/UW-GAC/GENESIS

135. Conomos MP, Miller MB, Thornton TA. Robust Inference of Population Structure for Ancestry Prediction and Correction of Stratification in the Presence of Relatedness. Genet Epidemiol. 2015;39: 276–293. doi:10.1002/gepi.21896

136. Zeng Y, Breheny P. The biglasso Package: A Memory- and Computation-Efficient Solver for Lasso Model Fitting with Big Data in R. ArXiv170105936 Stat. 2018 [cited 24 Feb 2020]. Available: http://arxiv.org/abs/1701.05936

137. Cingolani P, Platts A, Wang LL, Coon M, Nguyen T, Wang L, et al. A program for annotating and predicting the effects of single nucleotide polymorphisms, SnpEff: SNPs in the genome of Drosophila melanogaster strain w1118; iso-2; iso-3. Fly (Austin). 2012;6: 80–92. doi:10.4161/fly.19695

138. International Schizophrenia Consortium, Purcell SM, Wray NR, Stone JL, Visscher PM, O’Donovan MC, et al. Common polygenic variation contributes to risk of schizophrenia and bipolar disorder. Nature. 2009;460: 748–752. doi:10.1038/nature08185

139. Marees AT, de Kluiver H, Stringer S, Vorspan F, Curis E, Marie-Claire C, et al. A tutorial on conducting genome-wide association studies: Quality control and statistical analysis. Int J Methods Psychiatr Res. 2018;27: e1608. doi:10.1002/mpr.1608

140. Rosenberg NA, Edge MD, Pritchard JK, Feldman MW. Interpreting polygenic scores, polygenic adaptation, and human phenotypic differences. Evol Med Public Health. 2019;2019: 26–34. doi:10.1093/emph/eoy036

141. Gautier M, Vitalis R. rehh: an R package to detect footprints of selection in genome-wide SNP data from haplotype structure. Bioinforma Oxf Engl. 2012;28: 1176–1177. doi:10.1093/bioinformatics/bts115

142. Gautier M, Klassmann A, Vitalis R. rehh 2.0: a reimplementation of the R package rehh to detect positive selection from haplotype structure. Mol Ecol Resour. 2017;17: 78–90. doi:10.1111/1755-0998.12634

143. R Core Team. R: A language and environment for statistical computing. Vienna, Austria: R Foundation for Statistical Computing; Available: http://www.R-project.org/

144. Wickham H. ggplot2: Elegant Graphics for Data Analysis. New York: Springer-Verlag; 2016.

145. Wilke CO. cowplot: Streamlined Plot Theme and Plot Annotations for “ggplot2.” 2019. Available: https://CRAN.R-project.org/package=cowplot

146. Dowle M, Srinivasan A. data.table: Extension of ‘data.framè. 2019. Available: https://CRAN.R-project.org/package=data.table

147. Microsoft, Weston S. foreach: Provides Foreach Looping Construct for R. 2017. Available: https://CRAN.R-project.org/package=foreach

148. Revolution Analytics, Weston S. doMC: Foreach Parallel Adaptor for “parallel.” 2017. Available: Adaptor for ’parallel’. R package version 1.3.5. https://CRAN.R-project.org/package=doMC

149. Clarke E, Sherrill-Mix S. ggbeeswarm: Categorical Scatter (Violin Point) Plots. 2017. Available: https://CRAN.R-project.org/package=ggbeeswarm

150. Grolemund G, Wickham H. Dates and Times Made Easy with lubridate. J Stat Softw. 2011;40: 1–25. doi:10.18637/jss.v040.i03

151. Garnier S. viridis: Default Color Maps from “matplotlib.” 2018. Available: https://CRAN.R-project.org/package=viridis

